# The genetic architecture of human brainstem structures and their involvement in common brain disorders

**DOI:** 10.1101/811711

**Authors:** Torbjørn Elvsåshagen, Shahram Bahrami, Dennis van der Meer, Ingrid Agartz, Dag Alnæs, Deanna M. Barch, Ramona Baur-Streubel, Alessandro Bertolino, Mona K. Beyer, Giuseppe Blasi, Stefan Borgwardt, Birgitte Boye, Jan Buitelaar, Erlend Bøen, Elisabeth Gulowsen Celius, Simon Cervenka, Annette Conzelmann, David Coynel, Pasquale Di Carlo, Srdjan Djurovic, Sarah Eisenacher, Thomas Espeseth, Helena Fatouros-Bergman, Lena Flyckt, Barbara Franke, Oleksandr Frei, Barbara Gelao, Hanne Flinstad Harbo, Catharina A. Hartman, Asta Håberg, Dirk Heslenfeld, Pieter Hoekstra, Einar A. Høgestøl, Rune Jonassen, Erik G. Jönsson, Karolinska Schizophrenia Project (KaSP) consortium, Peter Kirsch, Iwona Kłoszewska, Trine Vik Lagerberg, Nils Inge Landrø, Stephanie Le Hellard, Klaus-Peter Lesch, Luigi A. Maglanoc, Ulrik F. Malt, Patrizia Mecocci, Ingrid Melle, Andreas Meyer-Lindenberg, Torgeir Moberget, Jan Egil Nordvik, Lars Nyberg, Kevin S. O’Connell, Jaap Oosterlaan, Marco Papalino, Andreas Papassotiropoulos, Paul Pauli, Giulio Pergola, Karin Persson, Dominique de Quervain, Andreas Reif, Jarek Rokicki, Daan van Rooij, Alexey A. Shadrin, André Schmidt, Emanuel Schwarz, Geir Selbæk, Hilkka Soininen, Piotr Sowa, Vidar M. Steen, Magda Tsolaki, Bruno Vellas, Lei Wang, Eric Westman, Georg Ziegler, Mathias Zink, Ole A. Andreassen, Lars T. Westlye, Tobias Kaufmann

**Author notes:** @ Corresponding authors: Torbjørn Elvsåshagen, M.D., Ph.D, Shahram Bahrami, Ph.D, & Tobias Kaufmann, Ph.D, Postal address: Norwegian Centre for Mental Disorders Research, Oslo University Hospital, PoBox 4956 Nydalen, Norway. These authors contributed equally to this work (Torbjørn Elvsåshagen, Shahram Bahrami).

## Abstract

Brainstem regions support critical bodily functions, yet their genetic architectures and involvement in brain disorders remain understudied. Here, we examined volumes of brainstem structures using magnetic resonance imaging in 43,353 individuals. In 27,034 genotyped healthy participants, we identified 16 genetic loci associated with whole brainstem volume and 10, 23, 3, and 9 loci associated with volumes of the midbrain, pons, superior cerebellar peduncle, and medulla oblongata, respectively. These loci were mapped to 305 genes, including genes linked to brainstem development and common brain disorders. We detected genetic overlap between the brainstem volumes and eight psychiatric and neurological disorders. Using imaging data from 16,319 additional individuals, we observed differential volume alterations in schizophrenia, bipolar disorder, multiple sclerosis, mild cognitive impairment, dementia, and Parkinson’s disease. Together, our results provide new insights into the genetic underpinnings of brainstem structures and support their involvement in common brain disorders.

## Main

The brainstem is a critical regulator of vital bodily functions and includes the midbrain, pons, and the medulla oblongata^1, 2^. Regions of the brainstem subserve emotions and behavior and are implicated in the pathophysiology of psychiatric and neurological diseases^3–6^. The monoaminergic brainstem nuclei may play central roles in mood, psychotic, and autism spectrum disorders^7–10^. Atrophy and lesions of brainstem structures are hallmarks of neurodegenerative and other neurological diseases^5, 11^. Despite their importance in human health and disease, the brainstem structures remain markedly understudied.

Magnetic resonance imaging (MRI) studies have revealed cortical and subcortical structural alterations in psychiatric and neurological disorders^12–15^, and the discovery of genetic contributions to brain structure variation has begun^16–18^. However, no large-scale neuroimaging study has focused on the genetic architecture of brainstem regions and their involvement in common brain disorders. The unprecedented availability of large imaging genetics resources^19^ and recent development of a Bayesian brainstem segmentation algorithm^20^ allowed us to estimate the volumes of midbrain, pons, medulla oblongata, superior cerebellar peduncle (SCP, which interconnects the pons and the cerebellum), and the whole brainstem in a large sample. We employed three complementary approaches to increase our knowledge of the genetic underpinnings of brainstem structures and their roles in common brain disorders. First, we conducted genome-wide association studies (GWAS) in healthy individuals to identify genetic loci associated with volumes of the brainstem structures. Second, we used summary statistics from recent large-scale GWAS of common brain disorders to assess genetic overlap between the disorders and volumes of the brainstem regions. Finally, we examined volumes of the brainstem structures in individuals with psychiatric or neurological illnesses in comparison to healthy controls.

## Results

We obtained raw T1 3D brain MRI data from a total of *n* = 49,815 individuals, collected through collaborations, data sharing platforms, and from in-house samples (Supplementary Tables 1-2). The MRI data was segmented into the whole brainstem, midbrain, pons, SCP, and medulla oblongata using Freesurfer 6.0^21^ and Bayesian brainstem segmentation, robust to differences in MRI scanners and pulse sequence details^20^. We assessed the delineations in all 49,815 data sets by visually inspecting twelve sagittal view figures of the segmentations for each participant (Supplementary Fig. 1). This procedure was conducted blind to case-control status and excluded 13% (*n* = 6,462) of the data sets, mainly due to insufficient field of view, image quality, and segmentation errors in the clinical samples. The final study sample of *n* = 43,353 participants (Supplementary Table 3) comprised healthy participants (*n* = 38,299, age range 3-95 years) and individuals with psychiatric or neurological disorders (*n* = 5,054, age range 5-96 years).

### GWAS reveals 61 genetic loci associated with brainstem volumes

The study sample included 27,034 genotyped healthy individuals aged 40-70 years from the UK Biobank^22^. Using MRI and single-nucleotide polymorphism (SNP) data from these participants, we conducted GWAS with PLINK v2.0^23^ on volumes of the midbrain, pons, SCP, medulla oblongata, and whole brainstem. All GWAS accounted for age, age², sex, scanning site, intracranial volume (ICV), genotyping batch, and the first ten genetic principal components to control for population stratification. In addition, the GWAS for the midbrain, pons, SCP, and medulla oblongata accounted for whole brainstem volume, thus revealing genetic signals beyond commonality in volume, analogous to a recent study of hippocampal subfields^24^. The GWAS for the brainstem structures were also run without covarying for whole brainstem volume.

SNP-based heritability estimated using LD score regression^25^ on the GWAS summary statistics was 32% for the whole brainstem, 29% for the midbrain, 31% for pons, 15% for SCP, and 23% for the medulla oblongata (all s.e. < 5%), illustrating the substantial genetic influence on brainstem volumes (Fig. 1a). We found genome-wide significant hits (*P* < 5e-8) for all brainstem volumes and identified a total of 125 independent significant SNPs across structures located in 61 genomic loci, using the Functional Mapping and Annotation of GWAS (FUMA) platform v1.3.3c^26^ (Fig. 1b-c and Supplementary Table 4). Sixteen genetic loci were associated with whole brainstem volume and 10, 23, 3, and 9 loci were associated with volumes of the midbrain, pons, SCP, and medulla oblongata, respectively. Sixteen loci were associated with more than one brainstem volume, thus resulting in 45 unique brainstem-associated genetic regions. Individual Manhattan and quantile-quantile (Q-Q) plots for each brainstem volume are provided in Supplementary Figs. 2-3. Supplementary Fig. 4 shows regional plots for the most significant genetic locus for each brainstem volume. Heritability estimates and GWAS hits for the brainstem regions without covarying for whole brainstem volume are provided in Supplementary Figs. 5-6 and Supplementary Table 5.

**Fig. 1.**
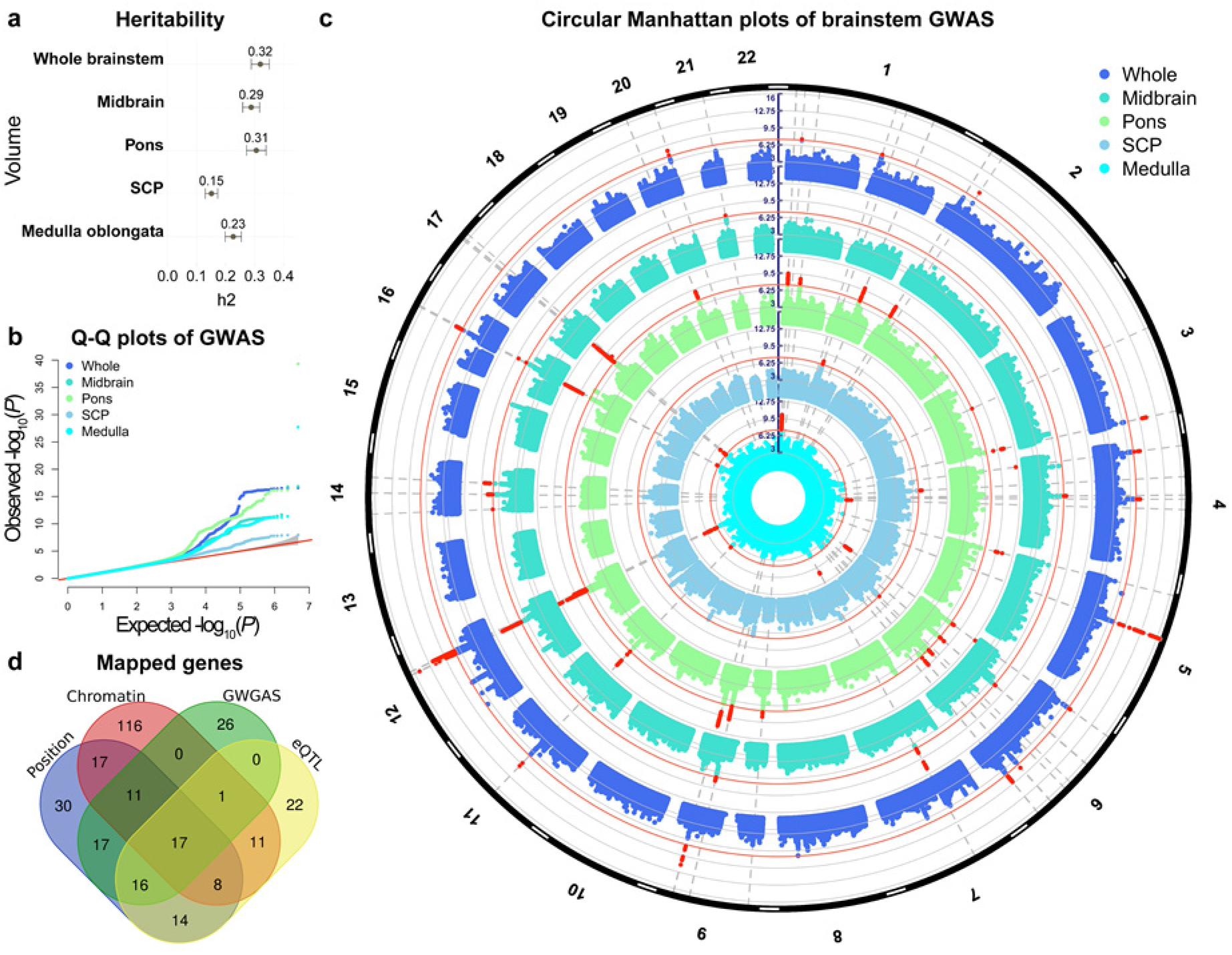
GWAS identifies 61 loci associated with brainstem volumes. **a**, Heritability estimates for the brainstem volumes of *n* = 27,034 healthy individuals. All brainstem volumes showed substantial heritability, with highest estimates for the whole brainstem (*h_2_ =* 0.32) and pons (*h_2_ =* 0.31) and lowest for the medulla oblongata (*h_2_ =* 0.23) and SCP (*h_2_ =* 0.15). **b**, Q-Q plots for the brainstem volumes. **c**, Circular Manhattan plots of GWAS for brainstem volumes. The outermost plot in blue reflects the GWAS of whole brainstem volume, whereas, from the periphery to center, the turquoise, green, grey/blue, and cyan plots indicate the GWAS of the midbrain, pons, SCP, and medulla oblongata volumes, respectively. Red circular lines indicate genome-wide significance and the red radial lines are significant loci. **d**, Venn diagram showing number of genes mapped by the four different strategies, i.e., positional gene, expression quantitative trait loci (eQTL), and chromatin interaction mapping, and identification by the GWGAS. Seventeen genes were identified by all four approaches. Whole; whole brainstem. SCP; superior cerebellar peduncle. Medulla; medulla oblongata. GW(G)AS; genome-wide (gene-based) association analyses.

We functionally annotated SNPs across the brainstem volumes that were in linkage disequilibrium (*r*^2^ ≥ 0.6) with one of the independent significant SNPs using FUMA. A majority of these SNPs were intronic (60.3%) or intergenic (23.7%) and 1.5% were exonic (Supplementary Tables 6-10). About 94% of the SNPs had a minimum chromatin state of 1 to 7, thus suggesting they were in open chromatin regions^27^. Supplementary Fig. 7 provides information for functional SNP categories for each brainstem volume. Two of the lead SNPs were exonic and associated with medulla oblongata (rs13107325) and whole brainstem (rs13388394) volumes. The combined annotation-dependent depletion (CADD) scores of those SNPs were 23.1 (rs13107325) and 17.7 (rs13388394), thus indicating deleterious protein effects^28^. rs13107325 is located in *SLC39A8* and has previously been associated with multiple traits, including schizophrenia (SCZ) and Parkinson’s disease (PD)^29^.

### Implicated genes and genome-wide gene-based associations

We used positional, expression quantitative trait loci (eQTL), and chromatin interaction mapping in FUMA^26^ to map the 125 independent significant SNPs to genes. These three strategies identified 280 unique genes, where 130, 89, and 181 genes were mapped by positional, eQTL, and chromatin interaction mapping, respectively. 168 of these were implicated by one mapping strategy, 68 genes by two strategies, and 25 of the genes were implicated by three strategies (Fig. 1d, and Supplementary Table 11). Supplementary Fig. 8 provides visualisation of mapped genes for each brainstem volume in Circos plots.

We then conducted genome-wide gene-based association analyses (GWGAS) using MAGMA^30^ and detected 87 unique genes across the brainstem volumes (Fig. 2 and Supplementary Table 12). Thirty-six were associated with whole brainstem volume and 22, 37, 10, and 17 genes were associated with volumes of the midbrain, pons, SCP, and the medulla oblongata, respectively. Twenty-two of the genes were only associated with whole brainstem volume, whereas 13, 14, 6, 5 genes were only significant for midbrain, pons, SCP, and the medulla oblongata volumes. The most strongly associated gene for each volume identified by the GWGAS was *RFX4* (*P* = 2.8e-15), *PARPBP* (*P* = 1.7e-11), *DRAM1* (*P* = 6.2e-15), *LMX1A* (*P* = 1.7e-10), and *HOXB3* (*P* = 2.0e-12) for the whole brainstem, midbrain, pons, SCP, and medulla oblongata, respectively. Supplementary Fig. 9 provides Q-Q plots for these GWGAS. We also found that 25 of the genes identified by GWGAS were not mapped by the GWAS analyses, resulting in a total number of 305 brainstem-linked genes identified by either GWAS or GWGAS. Moreover, supporting robustness, seventeen of the 87 genes identified by the GWGAS were also implicated by all three FUMA mapping strategies (Fig. 1d, Supplementary Table 13).

**Fig. 2.**
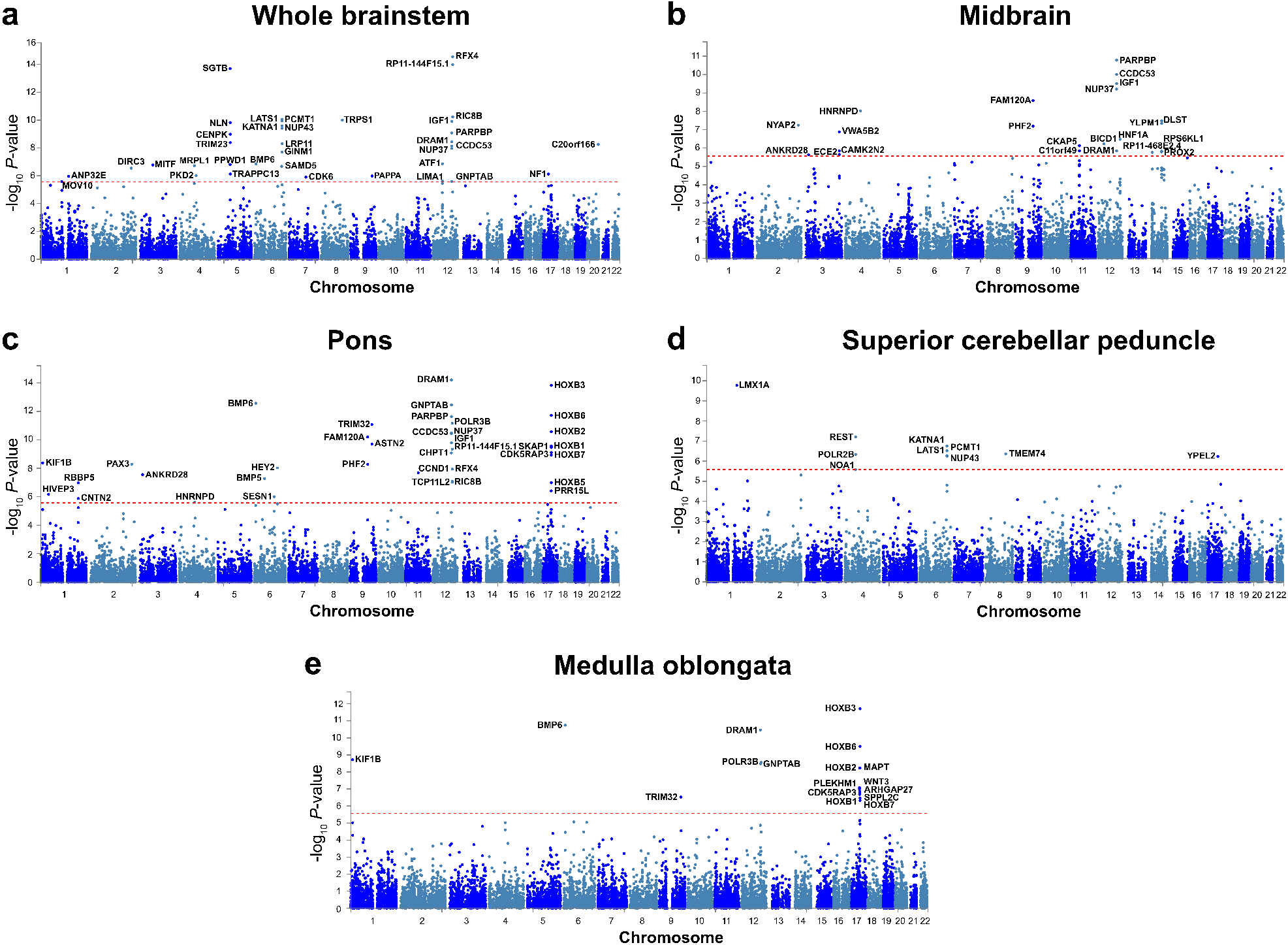
Manhattan plots from the genome-wide gene-based association analyses for volumes of the whole brainstem (**a**), midbrain (**b**), pons (**c**), superior cerebellar peduncle (**d**), and medulla oblongata (**e**). Thirty-six genes were associated with whole brainstem volume and 22, 37, 10, and 17 genes were associated with volumes of the midbrain, pons, superior cerebellar peduncle, and the medulla oblongata, respectively. Twenty-two of the genes were only associated with whole brainstem volume, whereas 13, 14, 6, 5 genes were only significant for volumes of the midbrain, pons, superior cerebellar peduncle, and the medulla oblongata. The red horizontal lines indicate genome-wide significance.

### Gene sets implicated by the significant genes

We conducted gene sets analyses and identified 78 Gene Ontology sets significantly associated with whole brainstem volume, and 34, 58, 6, and 56 gene sets associated with volumes of the midbrain, pons, SCP, and medulla oblongata, respectively (Supplementary Table 14). The most significant gene set for whole brainstem volume was *natural killer cell mediated immunity* (*P* = 2.47e-10), *positive regulation of epithelial cell proliferation* for midbrain (*P* = 8.97e-06), *anterior posterior pattern specification* for pons (*P* = 1.68e-11), *imp biosynthetic process* for SCP (*P* = 4.96e-06), and *embryonic skeletal system development* for medulla oblongata (*P* = 2.66e-14). Notably, *HOX* genes, which encode transcription factors with central roles in nervous system development^31, 32^ were included in the nine most significant gene sets for pons and in the 24 gene sets most strongly associated with medulla oblongata. We also employed the ConsensusPathDB^33^ to identify over-represented pathways for the mapped genes and found 13, 1, 25, and 58 significant pathways for volumes of the whole brainstem, pons, SCP, and medulla oblongata, respectively (Supplementary Table 15).

### Genetic overlap between brainstem volumes and common brain disorders

To further examine the polygenic architecture of brainstem volumes and the potential genetic overlap between brainstem regions and common brain disorders, we used GWAS summary statistics for attention-deficit/hyperactivity disorder (ADHD), autism spectrum disorder (ASD), bipolar disorder (BD), major depression (MD), SCZ, Alzheimer’s disease (AD), multiple sclerosis (MS), and PD, as outlined in Methods. We then generated conditional Q-Q plots^34–36^ for the brainstem regions and the eight clinical conditions. The conditional Q-Q plots compare the association with one trait (e.g., whole brainstem volume) across all SNPs and within SNPs strata determined by the significance of their association with another trait (e.g., SCZ). Polygenic overlap exists if the proportion of SNPs associated with the first trait increases as a function of the strength of association for the second trait and is visualized as a successive leftward deflection from the null distribution^34^. The conditional Q-Q plots for brainstem volumes and the clinical conditions showed successive increments of SNP enrichment for whole brainstem, midbrain, pons, SCP, and medulla oblongata (Supplementary Fig. 10), consistent with polygenic overlap across volumes and disorders. Conditional Q-Q plots illustrating the genetic overlap between whole brainstem volume and SCZ, BD, and PD are provided in Fig. 3a-c.

**Fig. 3.**
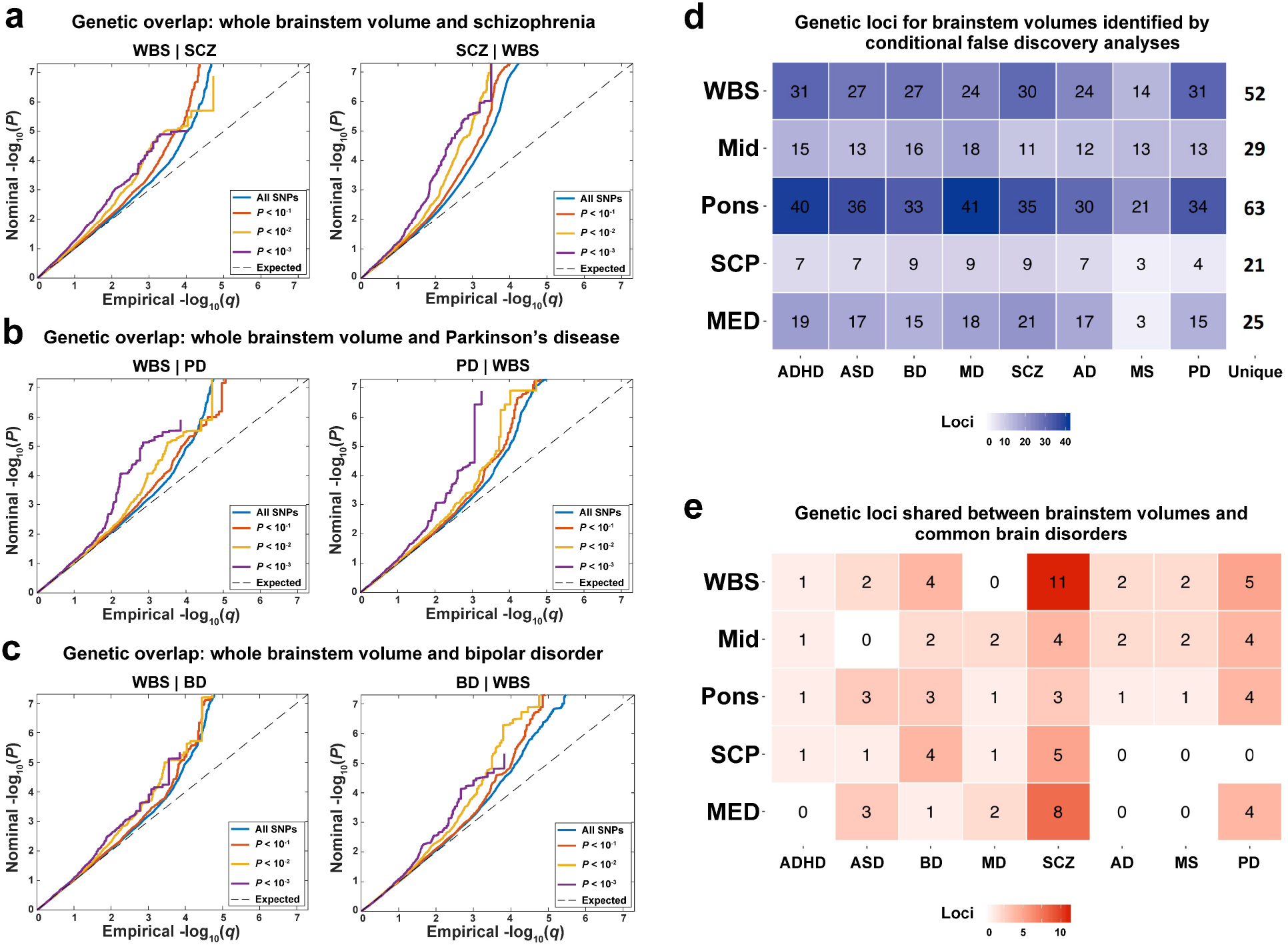
**a**, Conditional Q-Q plots for whole brainstem volume conditioned on SCZ (left) and vice versa (right), demonstrating genetic overlap. **b**, Conditional Q-Q plots for whole brainstem volume conditioned on PD (left) and vice versa (right), showing genetic overlap between these phenotypes. **c**, Conditional Q-Q plots for whole brainstem volume conditioned on BD (left) and vice versa (right), demonstrating genetic overlap. **d**, Enhanced discovery of genetic loci for each of the brainstem volumes when conditional false discovery rate analyses were run for each of the brainstem volumes conditioned on the eight brain disorders. These analyses revealed a total of 208 genetic loci for whole brainstem volume, and 111, 270, 55, and 125 loci for volumes of the midbrain, pons, SCP, and medulla oblongata, respectively. These genetic regions were located in 52 unique genetic loci for whole brainstem volume, and 29, 63, 21, and 25 unique loci for volumes of the midbrain, pons, SCP, and medulla oblongata. **e**, conjunctional false discovery rate analysis detected shared genetic loci across brainstem volumes and the eight clinical conditions. The largest numbers of shared loci were found for SCZ (31), BD (14), and PD (17), whereas 8, 4, 6, 9, and 5 genetic loci were jointly shared for ASD, ADHD, MD, AD, and MS, respectively, and the brainstem volumes. WBS; whole brainstem. MID; midbrain. SCP; superior cerebellar peduncle. MED; medulla oblongata. ADHD; attention-deficit/hyperactivity disorder. ASD; autism spectrum disorders. BD; bipolar disorder. MD; major depression. SCZ; schizophrenia. AD; Alzheimer’s disease. MS; multiple sclerosis. PD; Parkinson’s disease.

We leveraged the genetic overlap to discover more of the genetic underpinnings of brainstem volumes by employing conditional false discovery rate (FDR) statistics^37, 38^. The conditional FDR builds on an empirical Bayesian statistical framework, combines summary statistics from a trait of interest with those of a conditional trait, and thus increases power to detect genetic variants associated with the primary trait. We ran the conditional FDR analyses for each of the brainstem volumes conditioned on the eight disorders and discovered a total of 208 genetic loci for the whole brainstem, and 111, 270, 55, and 125 loci for the midbrain, pons, SCP, and medulla oblongata, respectively. These regions were located in 52 unique genetic loci for whole brainstem volume, and 29, 63, 21, and 25 unique loci for volumes of the midbrain, pons, SCP, and medulla oblongata, respectively (Fig. 3d, Supplementary Tables 16-20). The loci identified by the conditional FDR included all brainstem-associated genetic regions discovered by the GWAS. Supplementary Fig. 11 provides Manhattan plots for the genetic loci detected by the conditional FDR analyses for each brainstem region.

To further characterize the genetic overlap between brainstem volumes and the eight clinical conditions, we performed conjunctional FDR analyses, which enable detection of genetic loci shared between two phenotypes^34–36^. These analyses revealed shared loci across the brainstem structures and the clinical conditions (Fig. 3e). We found the largest number of loci shared between brainstem volumes and SCZ (31), BD (14), and PD (17). For ASD, ADHD, MD, AD, and MS, there were 9, 4, 6, 5, and 5 genetic loci jointly associated with the brainstem volumes and the disorders, respectively (Fig. 3e). Notably, the shared genetic loci exhibited a mixed pattern of allelic effect directions, i.e., disorder-linked genetic variants were associated with both larger and smaller brainstem volumes. Manhattan plots and details for the genetic loci shared between the eight clinical conditions and the brainstem volumes are provided in Fig. 4a-h and in Supplementary Table 21.

**Fig. 4.**
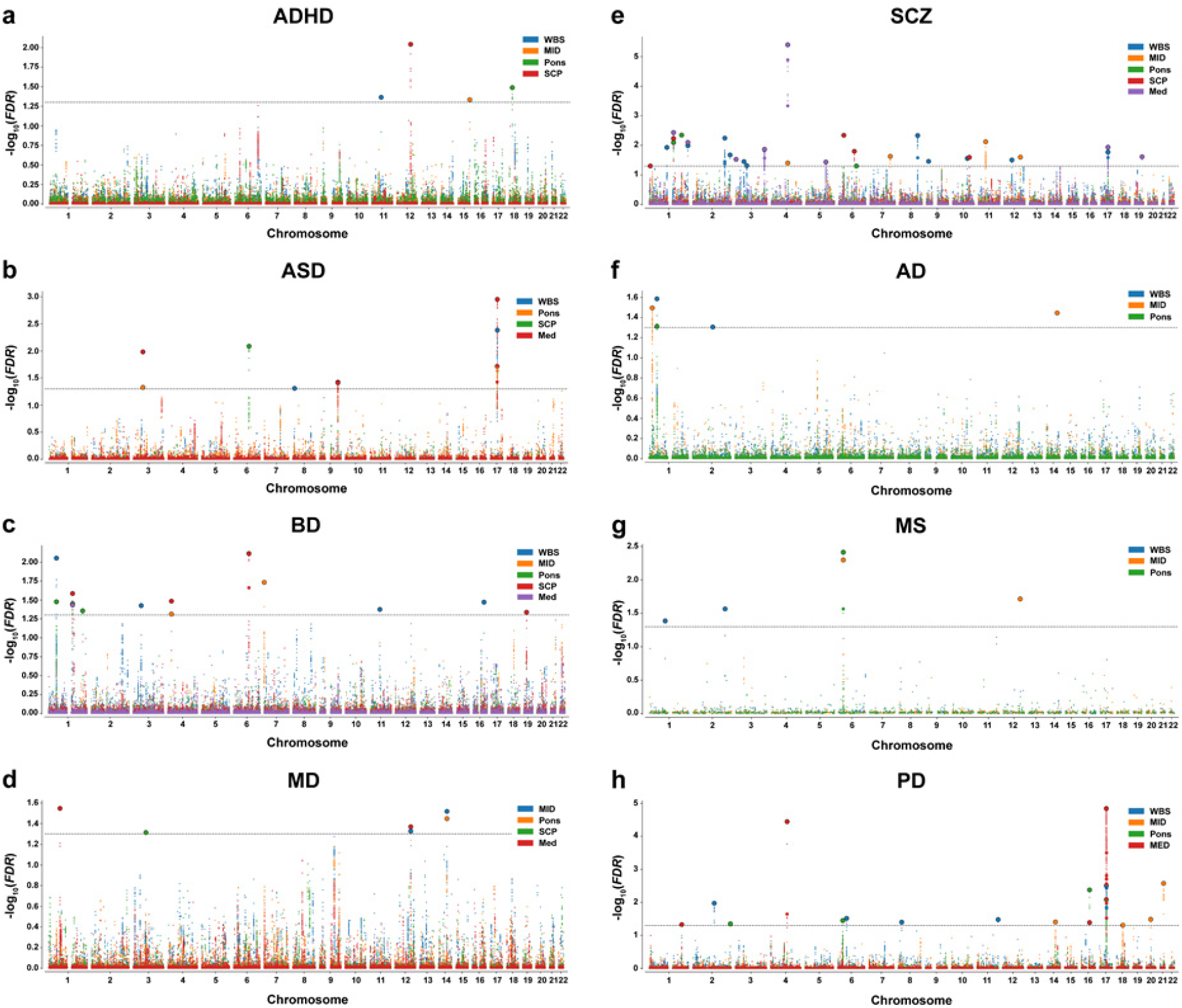
Manhattan plots for genetic loci shared between brainstem volumes and eight common brain disorders: **a**, 4 shared loci in ADHD, **b**, 9 shared loci in ASD, **c**, 14 shared loci in BD, **d**, 6 shared loci in MD, **e**, 31 shared loci in SCZ, **f**, 5 shared loci in AD, **g**, 5 shared loci in MS, and **h**, 17 shared loci in PD. WBS; whole brainstem. MID; midbrain. SCP; superior cerebellar peduncle. MED; medulla oblongata. ADHD; attention-deficit/hyperactivity disorder. ASD; autism spectrum disorders. BD; bipolar disorder. MD; major depression. SCZ; schizophrenia. AD; Alzheimer’s disease. MS; multiple sclerosis. PD; Parkinson’s disease.

We ran Gene Ontology gene sets analyses for genes nearest to the shared loci across the brainstem regions for each disorder and found 33 significant gene sets for SCZ, mainly involving central nervous system, neuronal, and cellular developmental processes (Supplementary Table 22). There were no significant gene sets for the other disorders.

We also examined genetic correlations between brainstem volumes and the common brain disorders using LD score regression^25^ (Supplementary Fig. 12). There were correlations with uncorrected *P* < 0.05, including positive associations between brainstem volumes and PD, yet these were not significant after multiple testing corrections.

### Brainstem volumes in common brain disorders

We compared brainstem volumes between individuals with common brain disorders and healthy controls (HC) (age range 5-96 years): ADHD (*n* = 681 patients/*n* = 992 HC), ASD (*n* = 125/*n* = 140), BD (*n* = 464/*n* = 1,513), major depressive disorder (MDD; *n* = 211/*n* = 93), SCZ (*n* = 1,044/*n* = 2,079), prodromal SCZ or at risk mental state (SCZRISK; *n* = 91/*n* = 402), non-SCZ psychosis spectrum diagnoses (PSYMIX; *n* = 308/*n* = 1,430), dementia (*n* = 756/*n* = 1,921), mild cognitive impairment (MCI; *n* = 987/*n* = 1,655), MS (*n* = 257/*n* = 1,053), and PD (*n* = 138/*n* = 67). Supplementary Tables 1-3 provide information on the individual cohorts. Linear models were run covarying for sex, age, age², ICV, and scanner site using R^39^. The analyses for volumes of midbrain, pons, SCP, and medulla oblongata were run both with and without covarying for whole brainstem volume, and were adjusted for multiple testing using FDR (Benjamini-Hochberg, accounting for all 99 tests). Fig. 5 depicts the resulting case-control differences.

**Fig. 5.**
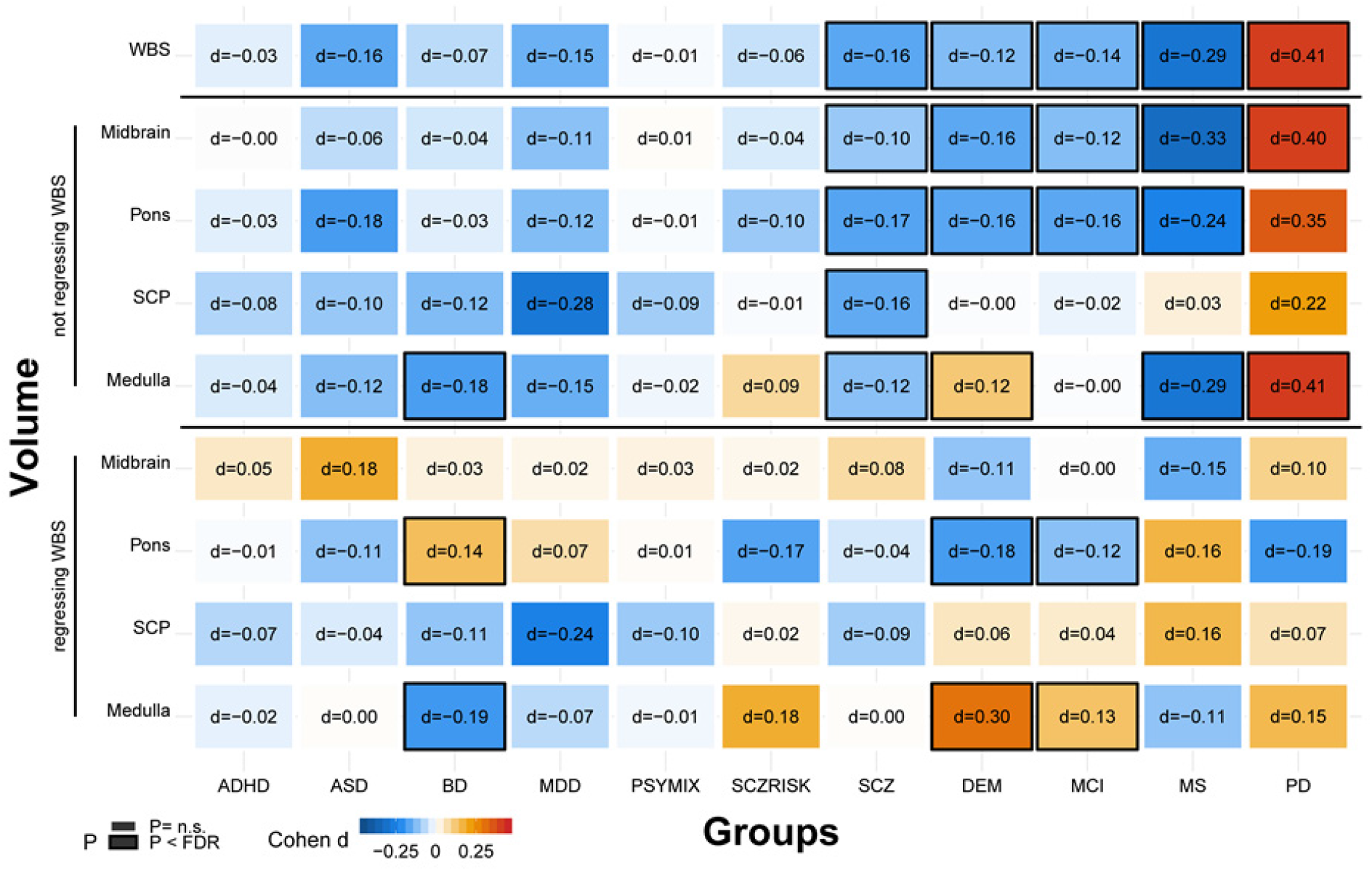
Volumes of brainstem structures in individuals with common brain disorders compared to healthy controls. There were differential volumetric alterations in individuals with BD, SCZ, DEM, MCI, MS, and PD after adjusting for multiple testing. ADHD; attention-deficit/hyperactivity disorder. ASD; autism spectrum disorders. BD; bipolar disorder. MDD; major depressive disorder. PSYMIX; non-SCZ psychosis spectrum diagnoses. SCZRISK; prodromal SCZ or at risk mental state. SCZ; schizophrenia. DEM; dementia. MCI; mild cognitive impairment. MS; multiple sclerosis. PD; Parkinson’s disease. WBS; whole brainstem. SCP; superior cerebellar peduncle. Medulla; medulla oblongata.

BD was associated with smaller medulla oblongata volume and larger pons volume, when accounting for the whole brainstem. Individuals with SCZ showed smaller volumes of all brainstem structures compared to HC, but not significantly for the midbrain, pons, and medulla oblongata when regressing out whole brainstem volume, consistent with a general effect across the brainstem regions. Volumes of whole brainstem, midbrain, and pons were smaller in the individuals with dementia compared to HC, whereas medulla oblongata volume was larger. A highly similar pattern was found for individuals with MCI, with smaller volumes of the whole brainstem, midbrain, and pons, and larger medulla oblongata volume when accounting for whole brainstem. Individuals with MS showed smaller volumes of the whole brainstem, midbrain, pons, and medulla oblongata, whereas individuals with PD had larger volume of the whole brainstem, midbrain, and medulla oblongata.

We ran further analyses of associations between brainstem volumes and clinical characteristics in the individuals with MCI, dementia, MS, SCZ, and PD and details of these analyses are provided in Supplementary Figs. 13-14. There were significant associations between Mini-Mental State Examination^40^ scores and brainstem volumes in dementia and MCI, indicating smaller pons and larger medulla oblongata volumes in more severely affected individuals (all *P* < 2e-04). In MS, there were brainstem volume decreases also in the subgroup of patients without infratentorial lesions (*n* = 91; all *P* < 0.05) and significant negative associations between the Expanded Disability Status Scale^41^ scores and brainstem volumes in patients with infratentorial lesions (*n* = 153; *P* < 0.05). There was no significant association between the Global Assessment of Functioning scale^42^ or Positive and Negative Syndrome Scale^43^ scores and brainstem volumes in individuals with SCZ. We found no evidence for tremor severity influencing brainstem volumes in individuals with PD.

## Discussion

The midbrain, pons, and medulla oblongata have central roles in human health and disease, yet no large-scale neuroimaging study has focused on their structure and genetic underpinnings. Here, we discovered novel genetic loci associated with brainstem volumes and found genetic overlap with eight psychiatric and neurological disorders, revealing that the brainstem may play important roles in common brain disorders. Indeed, leveraging clinical imaging data we found differential alterations of brainstem volumes in individuals with SCZ, BD, MS, dementia, MCI, and PD.

We identified 61 genetic loci associated with brainstem volumes using GWAS. Sixteen of these loci were associated with more than one volume, thus resulting in 45 unique brainstem-associated genetic regions. There is to our knowledge no previous study of the genetic underpinnings of midbrain, pons, SCP, and medulla oblongata volumes, yet a recent landmark study of ∼8,400 individuals in UK Biobank identified four SNPs on chromosomes four (rs10027331), nine (rs10983069), 11 (rs10792032), and 12 (rs11111090) associated with Freesurfer-based volume of the whole brainstem^44^. These SNPs are within four of the genetic loci linked to whole brainstem volume in the present study.

The brainstem volume-associated genetic loci detected by the GWAS of this study were linked to 305 genes. Seventeen of these genes were identified by both the GWGAS and by all three FUMA mapping strategies (Supplementary Table 13). Among these genes, *MAPT*, *KIF1B*, *KATNA1*, *NLN,* and *SGTB* are notable. *MAPT* encodes tau protein, which is produced throughout neurons of the brain^45^. Accumulation of tau is a hallmark of the neurodegenerative tauopathies, including AD and frontotemporal dementia^45^. *MAPT* has also been linked to PD^46^ and rare mutations and common variants of *MAPT* increase progressive supranuclear palsy risk, where brainstem volume loss is a central disease characteristic^47^. *KIF1B* is involved in axonal transport of mitochondria and synaptic vesicles and plays important roles in development of myelinated axons^48^. Loss of the gene results in impaired development of brainstem nuclei and impaired formation of synapses in the mouse spinal cord^49^. *KATNA1* is implicated in axon outgrowth regulation^50^ and neuronal migration during development^51^. *NLN* regulates neurotensin signaling^52^, which has been linked to the pathophysiology of psychiatric and neurological disorders, including SCZ and PD^53, 54^. The most significant locus for whole brainstem volume was mapped to *SGTB*, which is expressed at high levels in the brain and promotes neuronal differentiation and neurite outgrowth^55^. Although further studies are needed to clarify the relationship between these genes and brainstem structures, their implication by both the GWGAS and the three FUMA mapping approaches are suggestive of a role in brainstem volume variation.

The Gene Ontology gene sets analyses of the GWAS findings showed that *HOX* genes were included in the nine most significant gene sets for pons and in the 24 gene sets most strongly associated with medulla oblongata volume. In addition, nine *HOX* genes (*HOXB1-9*) were associated with volumes of both pons and medulla oblongata in the GWGAS. *HOX* genes encode Hox proteins, which are transcription factors with central roles in nervous system development^31, 32^. The *HOXB1-4* genes are critical for the development of the embryonic hindbrain, which gives rise to the pons, the medulla oblongata, and the cerebellum^32^. For example, *HOXB1* mutations can cause congenital bilateral facial palsy, hearing loss, and strabismus^56^. The *HOX* genes are not, however, expressed in the embryonic midbrain, which develops into the midbrain. Consistent with the embryonic genetic division between the hindbrain and the midbrain, *HOX* genes were not associated with the midbrain in the gene sets or in the GWGAS analyses of the current study.

There was polygenic overlap between the brainstem regions and the eight psychiatric and neurological disorders of the present study. We leveraged the genetic overlap to uncover more of the genetic architecture of the brainstem volumes and identified 52, 29, 63, 21, and 25 loci associated with volumes of the whole brainstem, midbrain, pons, SCP, and medulla oblongata, respectively, using conditional FDR. These loci included all brainstem-associated genetic regions identified by the GWAS. The polygenic overlap also indicates a role for brainstem regions in common brain disorders and gene sets analyses implicated cellular and neurodevelopmental processes in the genetic loci shared with SCZ.

Further studies of how the overlapping genetic regions influence brainstem structure and the risk for common brain disorders are warranted, yet several of the shared loci are noteworthy. The most significant shared locus for SCZ and the second-most significant shared locus for PD was rs13107325, which was associated with midbrain volume in SCZ and medulla oblongata volume in both disorders. rs13107325 is located in the metal ion transporter gene *SLC39A8*. We also found that rs4845679 was jointly associated with volumes of pons, SCP, and medulla oblongata and both SCZ and BD. The nearest gene for rs4845679 is *KCNN3*, which is expressed at high levels in the adult brain and encodes a protein that contributes to the afterhyperpolarization in neurons^57^. The most significant locus for ASD was rs9891103, which was jointly associated with whole brainstem volume, and its nearest gene was *MAPT*. rs8070942 and rs3865315 were shared between ASD and SCZ, respectively, and medulla oblongata volume. The nearest gene for these SNPs was *KANSL1*, which is expressed in the brain and encodes a nuclear protein involved in histone acetylation^58^.

We also found that the genetic loci shared between brainstem structures and the brain disorders exhibited a mixed pattern of allelic effect directions, i.e., disorder-linked genetic variants were associated with both larger (same effect direction) and smaller (opposite effect direction) brainstem volumes. A consistent direction of effect across overlapping genetic loci is a requirement for a significant genetic correlation as assessed using LD score regression^25^. For example, a recent study showed that SCZ and educational attainment may share >8K causal genetic variants, yet their genetic correlation is close to zero due to shared variants with opposite effect directions^59^. Thus, a mixed pattern of allelic effect directions might be one explanation for the lack of robust genetic correlations between the brainstem volumes and the disorders in the present study.

We detected brainstem volume differences between individuals with SCZ, BD, dementia, MCI, MS, and PD and their respective HC groups. The monoaminergic nuclei of the brainstem are implicated in psychotic and mood disorders^4, 60–63^, yet there are few volumetric studies of brainstem regions in these illnesses. The results of the present study suggest a general volume decrease across brainstem regions in SCZ, consistent with previous studies of the whole brainstem^64, 65^. BD, on the other hand, was associated with reduced volume of the medulla oblongata and a relative sparing or even increase of pons volume in the current study. Whether brainstem differences in SCZ and BD are genetically mediated and involved in the development of these disorders or illness effects that emerge during the course of the diseases mandates future studies.

Compared to healthy peers, individuals with dementia showed smaller volumes of the midbrain and pons and larger relative volume of medulla oblongata. Notably, we found a highly similar pattern in individuals with MCI. To our knowledge, there is no previous study showing reduced brainstem volumes in MCI, although one recent report found greater whole brainstem volume reduction over one year in individuals with MCI that converted to dementia than in those who did not convert^66^. There is a scarcity of structural brainstem studies in dementia, yet the results of the present study are consistent with a few previous findings suggesting volume decreases mainly in midbrain and pons in dementia^20, 67, 68^. Here, we extend these findings to MCI, thus suggesting that structural midbrain and pons alterations could be present in the early phases of dementia. The smaller volumes of whole brainstem, midbrain, pons, and the medulla oblongata in individuals with MS are consistent with the limited number of previous volumetric brainstem studies of the disorder^69–71^.

We found larger volumes of the whole brainstem, midbrain, and medulla oblongata in the individuals with PD. There was no indication that tremor severity could explain the volume increases. Notably, some previous studies detected enlargement of the brainstem and other brain structures in PD^72–74^ and the individuals with PD of the present study were in the early phase of the disorder and none used anti-Parkinson drugs. However, the PD sample was small and replication studies are needed to further explore how clinical characteristics, such as disorder phase and medication use, and potential confounds, including within-scanner motion, may factor into measurements of brainstem volumes in PD.

The resolution of the MRI data of the present study does not allow for analyses of individual brainstem nuclei. We also note that the effect sizes for the brainstem changes in the brain disorders of this study were small to moderate. However, larger effect sizes might be revealed in future studies of brainstem nuclei and the effects observed in the present study should not be interpreted as clinically insignificant. Rather, the findings of this study highlight the potential importance of the brainstem across psychiatric and neurological disorders and should stimulate research efforts to further clarify the roles of brainstem subregions in the etiologies and treatments of common brain disorders.

In summary, the current study provides new insights into the genetic architecture of brainstem regions, identifies the first genetic loci linked to volumes of the midbrain, pons, SCP, and the medulla oblongata, and shows genetic and imaging evidence for an involvement of brainstem regions in common brain disorders. Altogether, these findings encourage further studies of brainstem structures in human health and disease.

## Methods

Methods are available at http://…..

## Acknowledgments

The author list between I.A. and M.Z. is in alphabetic order. The authors were funded by the South-Eastern Norway Regional Health Authority 2015-078 (T.E.), 2013-123 (O.A.A.), 2014-097 (L.T.W.), 2015-073 (L.T.W.) and 2016-083 (L.T.W.), by the Research Council of Norway (276082 LifespanHealth (T.K.), 213837 (O.A.A), 223273 NORMENT (O.A.A.), 204966 (L.T.W.), 229129 (O.A.A.), 249795 (L.T.W.), 273345 (L.T.W.) and 283798 SYNSCHIZ (O.A.A.)), Stiftelsen Kristian Gerhard Jebsen, the European Research Council (ERCStG 802998 BRAINMINT (L.T.W.)), NVIDIA Corporation GPU Grant (T.K.), the Ebbe Frøland foundation, and a research grant from Mrs. Throne-Holst. The data used in this study was gathered from various sources and a detailed overview of the included cohorts and acknowledgement of their respective funding sources and cohort-specific details is provided in Supplementary Table 1. Data used in preparation of this article were obtained from the Alzheimer’s Disease Neuroimaging Initiative (ADNI) database (adni.loni.usc.edu), from the AddNeuroMed consortium, and from the Pediatric Imaging, Neurocognition and Genetics Study (PING) database (www.chd.ucsd.edu/research/ping-study.html, now shared through the NIMH Data Archive (NDA)). The investigators within the ADNI and PING contributed to the design and implementation of ADNI/PING and/or provided data but did not participate in analysis or writing of this report. This publication is solely the responsibility of the authors and does not necessarily represent the views of the National Institutes of Health or PING investigators. Complete listings of participating sites and study investigators can be found at http://adni.loni.usc.edu/wp-content/uploads/how_to_apply/ADNI_Acknowledgement_List.pdf and https://ping-dataportal.ucsd.edu/sharing/Authors10222012.pdf. The AddNeuroMed consortium was led by Simon Lovestone, Bruno Vellas, Patrizia Mecocci, Magda Tsolaki, Iwona Kłoszewska, and Hilkka Soininen.

We would like to thank the research participants and employees of the ADHD, ASD, SCZ, BD, and MD Working Groups of the Psychiatric Genomics Consortium, the International Genomics of Alzheimer’s Project, the International Multiple Sclerosis Genetics Consortium, International Parkinson Disease Genomics Consortium, and 23andMe, Inc. for granting us access to their GWAS summary statistics, and the many people who provided DNA samples for their studies. Data used in the preparation of this article were obtained from the Parkinson’s Progression Markers Initiative (PPMI) database (www.ppmi-info.org/data). For up-to-date information on the study, visit www.ppmi-info.org. This work was performed on the TSD (Tjeneste for Sensitive Data) facilities, owned by the University of Oslo, operated and developed by the TSD service group at the University of Oslo, IT-Department (USIT) and on resources provided by UNINETT Sigma2 - the National Infrastructure for High Performance Computing and Data Storage in Norway.

## Author contributions

T.E., S.B., L.T.W., and T.K. conceived the study, T.K., L.T.W., and S.B. preprocessed all MRI and genetic data, T.E. and T.K. performed quality control of the MRI data, T.E., S.B., and T.K. performed the analyses, and T.E., S.B., D.v.d.M., O.A.A., L.T.W., and T.K. interpreted the results. All remaining authors contributed to data collection at their respective sites as well as sample-specific tasks. T.E., S.B., and T.K. wrote the first draft of the paper and all authors contributed to and approved the final manuscript.

## Competing interests

Some authors received speaker’s honoraria from Lundbeck (T.E., G.B., and O.A.A.), Janssen Cilag (T.E.), Merck (E.H.), Sanofi Genzyme (E.H.), and Synovion (O.A.A). A.B. received speaker’s honoraria from Lundbeck, Otsuka, and Janssen Cilag and consultation fees from Biogen and Roche. J.K.B. has been a consultant to, member of advisory board of, and/or speaker for Shire, Roche, Medice, and Servier. E.G.C. has received personal fees from Almirall, Biogen, Merck, Roche and Teva, and grants and personal fees from Novartis and Sanofi. S.C. has received grant support from AstraZeneca as a coinvestigator and has served as a speaker for Otsuka. H.F.F. has received travel support, honoraria for advice or lecturing from Biogen Idec, Sanofi Genzyme, Merck, Novartis, Roche, and Teva and an unrestricted research grant from Novartis. N.I.L. has received consultation fees and travel support from Lundbeck. H.S. has received fees for advisory boards from ACImmune and Merck. P.S. has received honoraria for lecturing and travel support from Merck. M.T. has been member of advisory boards for Merck, IASIS Healthcare, ELPEN and FarmaSyn. M.Z. has received speaker fees for lectures, travel support and membership in advisory boards from Janssen Cilag, Lundbeck, Otsuka, Ferrer Pharma, Trommsdorff, Servier, and Roche. None of these external parties had any role in the analysis, writing or decision to publish this work. All other authors declare no competing interests.

## Additional information

**Supplementary information** is available for this paper at …

## Karolinska Schizophrenia Project (KaSP)

Members of the Karolinska Schizophrenia Project (KaSP): L Farde^6^, L Flyckt^6^, G Engberg^58^, S Erhardt S^58^, H Fatouros-Bergman^6^, S Cervenka^6^, L Schwieler^58^, F Piehl^59^, I Agartz^1,5,6^, K Collste^6^, P Victorsson^6^, A Malmqvist^58^, M Hedberg^58^, F Orhan^58^, C M Sellgren^58^

## Online methods

### Samples

We collected data from cohorts of participants with common brain disorders and healthy individuals through collaborations, data sharing platforms, and from in-house samples (*n* = 49,815). All included samples have been part of previously published works and data collection for each sample was performed with participants’ written informed consent and with approval by the respective local Institutional Review Boards. Supplementary Table 1 provides details for each sample and refers to previously published works from the included samples.

### Preprocessing of MRI data, brainstem segmentations, and quality control procedures

Raw T1-weighted MRI data for all individuals was stored and analyzed locally at the University of Oslo. The whole brainstem, midbrain, pons, SCP, and medulla oblongata were then delineated using Freesurfer 6.0^21^ and Bayesian brainstem segmentation^20^. The brainstem segmentation method is based on a probabilistic atlas and Bayesian inference and is robust to changes in MRI scanners and pulse sequence details^20^. We then manually assessed the delineations in all MRI data sets (*n* = 49,815) by visually inspecting twelve sagittal view figures of the segmentations for each participant, as shown in Supplementary Fig. 1. This visual quality control (QC) procedure for each data set was conducted blind to case-control status. Data sets were excluded from the study if one of the following requirements was not met: 1. the field of view included the whole brainstem, 2. the superior boundary of the midbrain approximated an axial plane through the mammillary body and the superior edge of the quadrigeminal plate, 3. the boundary between mibrain and pons approximated an axial plane through the superior pontine notch and the inferior edge of the quadrigeminal plate, 4. the boundary between between pons and medulla oblongata approximated an axial plane at the level of the inferior potine notch, 5. the inferior boundary of the medulla oblongata approximated an axial plane at the level of the posterior rim of the foramen magnum, 6. there were no substantial segmentation errors for the anterior and posterior boundaries of midbrain, pons, and medulla oblongata, and 7. the superior boundary of the SCP approximated the inferior boundary of the midbrain tectum, the inferior boundary of the SCP was defined by the merging with the cerebellum, and the anterior boundary of the SCP was defined by the posterior boundary of the pons. This QC procedure excluded 13% (*n* = 6,462) of the data sets, mainly due to insufficient field of view (e.g., not fully covering the inferior part of the medulla oblongata), insufficient data quality, and segmentation errors in the clinical samples, resulting in a final sample size of *n* = 43,353 (Supplementary Table 3).

### Genome-wide association studies for brainstem volumes and identification of genomic loci

The genetic analyses for the brainstem volumes were based on MRI and genetic data from healthy individuals of the UK Biobank Resource (sample size *n* = 27,034 after the QC procedures). We restricted all genetic analyses to individuals with White European ancestry, as determined by the UK Biobank study team. We applied standard quality control procedures to the UK Biobank v3 imputed genetic data, removing SNPs with an imputation quality score < 0.5, a minor allele frequency < 0.05, missing in more than 5% of individuals, and failing the Hardy Weinberg equilibrium tests at a *P* < 1e-6.

We performed GWAS on the brainstem volumes in the 27,034 healthy adults using PLINK 2.0^23^. All GWAS accounted for age, age², sex, scanning site, ICV, genetic batch, and the first ten genetic principal components to account for population stratification. In addition, the GWAS for the midbrain, pons, SCP, and medulla oblongata accounted for whole brainstem volume. The MHC region was excluded from the analysis.

We identified genetic loci related to brainstem volumes using the FUMA platform v1.3.3c^26^. Independent significant SNPs were identified by the genome-wide significant threshold (*P* < 5e-8) and by their independency (*r*^2^ ≤ 0.6 within a 1 mb window). Independent significant SNPs with *r*^2^ < 0.1 within a 1 mb window were defined as lead SNPs. Genomic risk loci were found by merging lead SNPs if they were closer than 250 kb. Candidate SNPs were defined as all SNPs in LD (*r*^2^ ≥ 0.6) with one of the independent significant SNPs in the genetic loci.

### Functional annotation, gene-based association, and gene-set analysis

We functionally annotated all candidate SNPs of brainstem volumes that were in linkage disequilibrium (*r*^2^ ≥ 0.6) with one of the independent significant SNPs using the FUMA platform v1.3.3c^26^. FUMA is based on information from 18 biological repositories and tools and functionally annotates GWAS results. The platform prioritizes the most likely causal SNPs and genes by combining positional, eQTL, and chromatin interaction mapping^26^. FUMA annotates significantly associated SNPs with functional categories, combined CADD scores^28^, RegulomeDB scores^77^, and chromatin states^26^. A CADD score above 12.37 is suggestive of a deleterious protein effect^28^. The RegulomeDB score indicates the regulatory functionality of SNPs based on eQTLs and chromatin marks, whereas the chromatin state indicates the accessibility of genomic regions accessibility using 15 categorical states, as predicted by ChromHMM based on 5 chromatin marks for 127 epigenomes^78, 79^.

We conducted genome-wide gene-based association and gene-set analyses using MAGMA^30^ in FUMA on the complete GWAS input data. MAGMA performs multiple linear regression to obtain gene-based *P*-values and the Bonferroni-corrected significant threshold was *P* = 0.05/18158 genes = 2.75e-6. We performed a MAGMA^30^ gene-set analysis for curated gene sets and GO terms obtained from MsigDB^80^. To identify over-represented pathways for the mapped genes, we used the ConsensusPathDB^33^. ConsensusPathDB is a database system that integrates functional interactions, including binary and complex protein-protein, genetic, metabolic, signaling, gene regulatory and drug-target interactions, as well as biochemical pathways^33^.

### Analyses of genetic overlap between brainstem volumes and eight brain disorders

To further examine the genetic architecture of brainstem volumes and the genetic relationships between brainstem regions and common brain disorders, we obtained GWAS summary statistics for ADHD^81^, ASD, SZ, and BD from the Psychiatric Genomics Consortium^82–84^, for MD from the Psychiatric Genomics Consortium and 23andMe^85, 86^, for AD from the International Genomics of Alzheimer’s Project^87^, for MS from the International Multiple Sclerosis Genetics Consortium^88^, and for PD from the International Parkinson Disease Genomics Consortium^46, 89^. We then employed conditional Q-Q plots^90^ and conditional FDR and conjunctional FDR statistics^34, 91^ to assess polygenic overlap between brainstem volumes and the eight brain disorders.

The conditional Q-Q plots compare the association with a primary trait across all SNPs and within SNPs strata determined by their association with the secondary trait. Genetic overlap exists if the proportion of SNPs associated with a phenotype increases as a function of the strength of the association with a secondary phenotype^90^. In conditional Q-Q plots, this enrichment is visualized as successive leftward deflections from the null distribution, and can be directly interpreted in terms of the true discovery rate (1−FDR)^34–36^. In this work, we plotted the empirical cumulative distribution of nominal *P*-values in one phenotype (e.g., whole brainstem volume) for all SNPs and for subsets of SNPs with significance levels in another phenotype (e.g., SCZ) below the indicated cut-offs (*P* ≤ 1, *P* ≤ 0.1, *P* ≤ 0.01, and *P* ≤ 0.001).

The conditional FDR statistical framework applies genetic association summary statistics from a trait of interest together with those of a conditional trait to estimate the posterior probability that a SNP has no association with the primary trait, given that the *P*-values for that SNP in both the primary and conditional traits are lower than the observed *P*-value^34–36^. This method can enhance the detection of genetic variants associated inserted the primary trait via re-ranking SNPs compared to nominal *P*-value based ranking. Here, we used an FDR level of 0.05 per pairwise comparison for conditional FDR.

To detect genetic jointly associated with the brainstem volumes and the eight clinical conditions, we used the conjunctional FDR method at a threshold of 0.05^34–36^. The conjunctional FDR is an extension of conditional FDR and is defined by the maximum of the two conditional FDR values for a specific SNP. This method estimates a posterior probability that a SNP is null for either trait or both at the same time, given that the *P* values for both phenotypes are as small, or smaller, than the *P*-values for each trait individually. Manhattan plots were constructed based on the ranking of the conjunctional FDR to show the genomic location of the shared genetic risk loci. The empirical null distribution in GWASs is affected by global variance inflation and all p-values were therefore corrected for inflation using a genomic inflation control procedure. All analysis was performed after excluding SNPs in the major extended histocompatibility complex (hg19 location Chr 6: 25119106–33854733) and 8p23.1 regions (hg19 location Chr 8: 7242715– 12483982) for all cases and *MAPT* and *APOE* regions for PD and AD, respectively, since complex correlations in regions with intricate LD can bias the FDR estimation. We also ran pairwise genetic correlations between brainstem volumes and the eight psychiatric and neurological disorders using LD score regression^25^. Here, the SNPs were pruned using a pairwise correlation coefficient approximation to LD (*r*²), where SNPs were disregarded at *r²* < 0.2 and pruning performed with 20 iterations^90^.

### Statistical analysis of brainstem volumes, brain disorders, and clinical variables

Statistical analyses for group comparisons were conducted using linear models in R statistics^39^. We included all healthy individuals that were imaged on the same scanners as the patients they were compared with, in the respective control groups. For clinical conditions where patients were imaged on multiple scanners, we included scanner site as a covariate in the analyses. For each of the clinical conditions, we ran linear models covarying for sex, age, age-orthogonalized age², ICV, and adjusted for multiple testing using FDR (Benjamini-Hochberg). The group analyses for volumes of midbrain, pons, SCP, and medulla oblongata were run both with and without covarying for whole brainstem volume.

Information concerning illness severity was available from individuals with MCI, dementia, MS, SCZ, and PD. 1610 individuals with MCI or dementia had MMSE score^40^, whereas 190 individuals with MS had EDSS scores^41^. Linear models were run to examine the relationships between the clinical variables and brainstem volumes covarying for sex, age, age-orthogonalized age², ICV, and scanner site. Two neuroradiologists assessed the imaging data from the individuals with MS and found that *n* = 153 participants had infratentorial MS lesions detectable with MRI, whereas *n* = 91 did not. 384 individuals with SCZ had function scores of the Global Assessment of Functioning scale^42^, whereas 264 individuals had symptom scores from the scale. 616 and 614 individuals with SCZ had positive and negative scores, respectively, from the Positive and Negative Syndrome Scale^43^. 128 individuals with PD had Unified Parkinson’s Disease Rating Scale III scores^75^ and the Hoehn and Yahr Stage score^76^. To examine whether tremor level might influence the measurements of brainstem volumes in PD, we used the self-report tremor item 2.10 of the Unified Parkinson’s Disease Rating Scale III^75^ and examined brainstem volumes across these tremor scores using linear models.

### Code availability

The code needed to reproduce the results is available from the authors upon request.

## Supplementary figures

**Supplementary Fig. 1.**
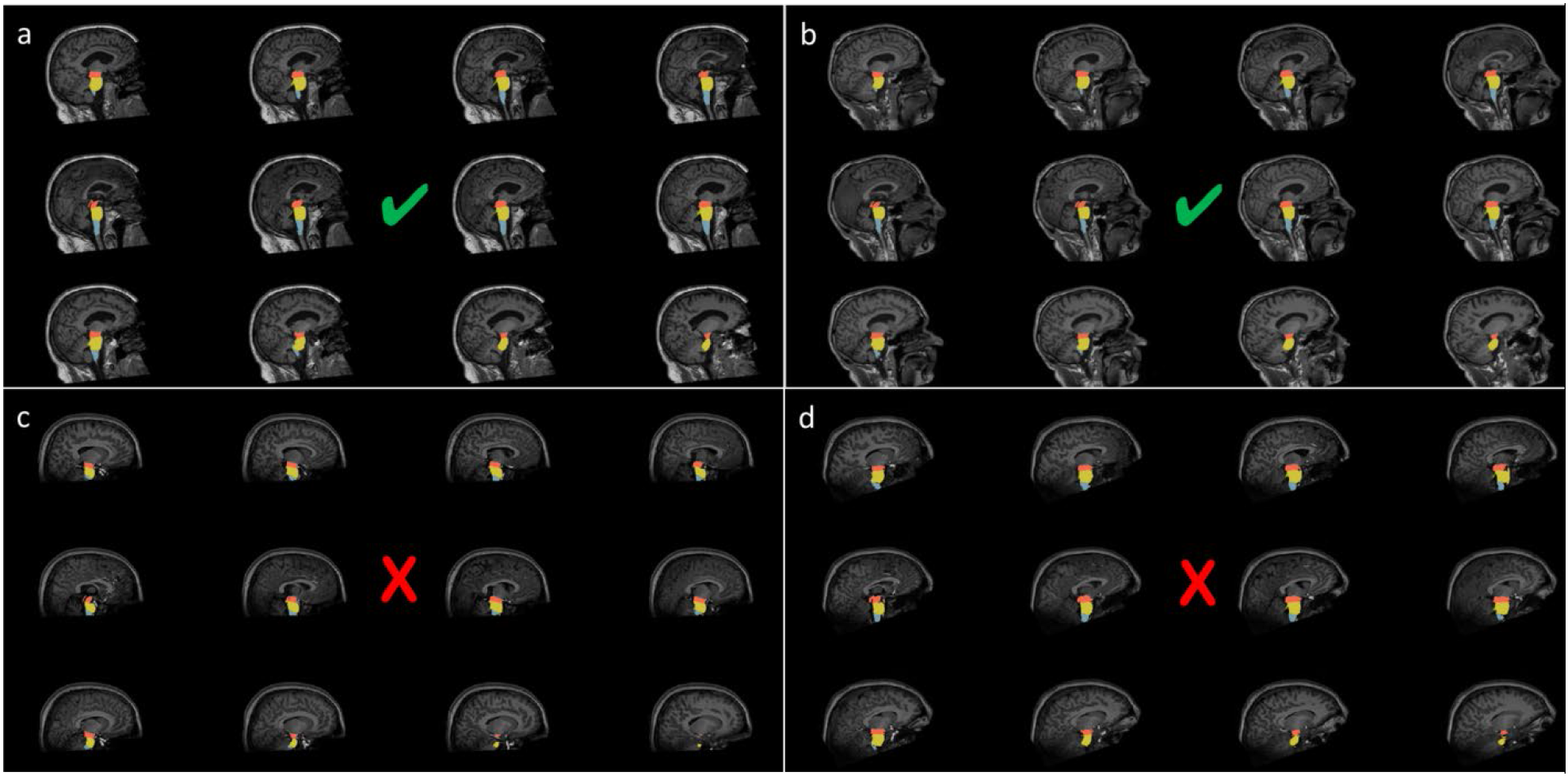
We manually assessed the delineations in all magnetic resonance imaging data sets (*n* = 49,815) by visually inspecting twelve sagittal view figures of the segmentations for each participant, as illustrated in **a**-**d**. **a** and **b** are examples of two datasets included in the study, whereas **c** and **d** are data sets excluded due to insufficient field of view (FOV). Data sets were excluded from the study if one of the following requirements was not met: 1. the field of view included the whole brainstem, 2. the superior boundary of the midbrain approximated an axial plane through the mammillary body and the superior edge of the quadrigeminal plate, 3. the boundary between mibrain and pons approximated an axial plane through the superior pontine notch and the inferior edge of the quadrigeminal plate, 4. the boundary between between pons and medulla oblongata approximated an axial plane at the level of the inferior potine notch, 5. the inferior boundary of the medulla oblongata approximated an axial plane at the level of the posterior rim of the foramen magnum, 6. there were no substantial segmentation errors for the anterior and posterior boundaries of midbrain, pons, and medulla oblongata, and 7. the superior boundary of the SCP approximated the inferior boundary of the midbrain tectum, the inferior boundary of the SCP was defined by the merging with the cerebellum, and the anterior boundary of the SCP was defined by the posterior boundary of the pons.This visual quality control procedure excluded 13% (*n* = 6,462) of the data sets, mainly due to insufficient FOV, image quality, and segmentation errors in the clinical samples. SCP; superior cerebellar peduncle.

**Supplementary Fig. 2.**
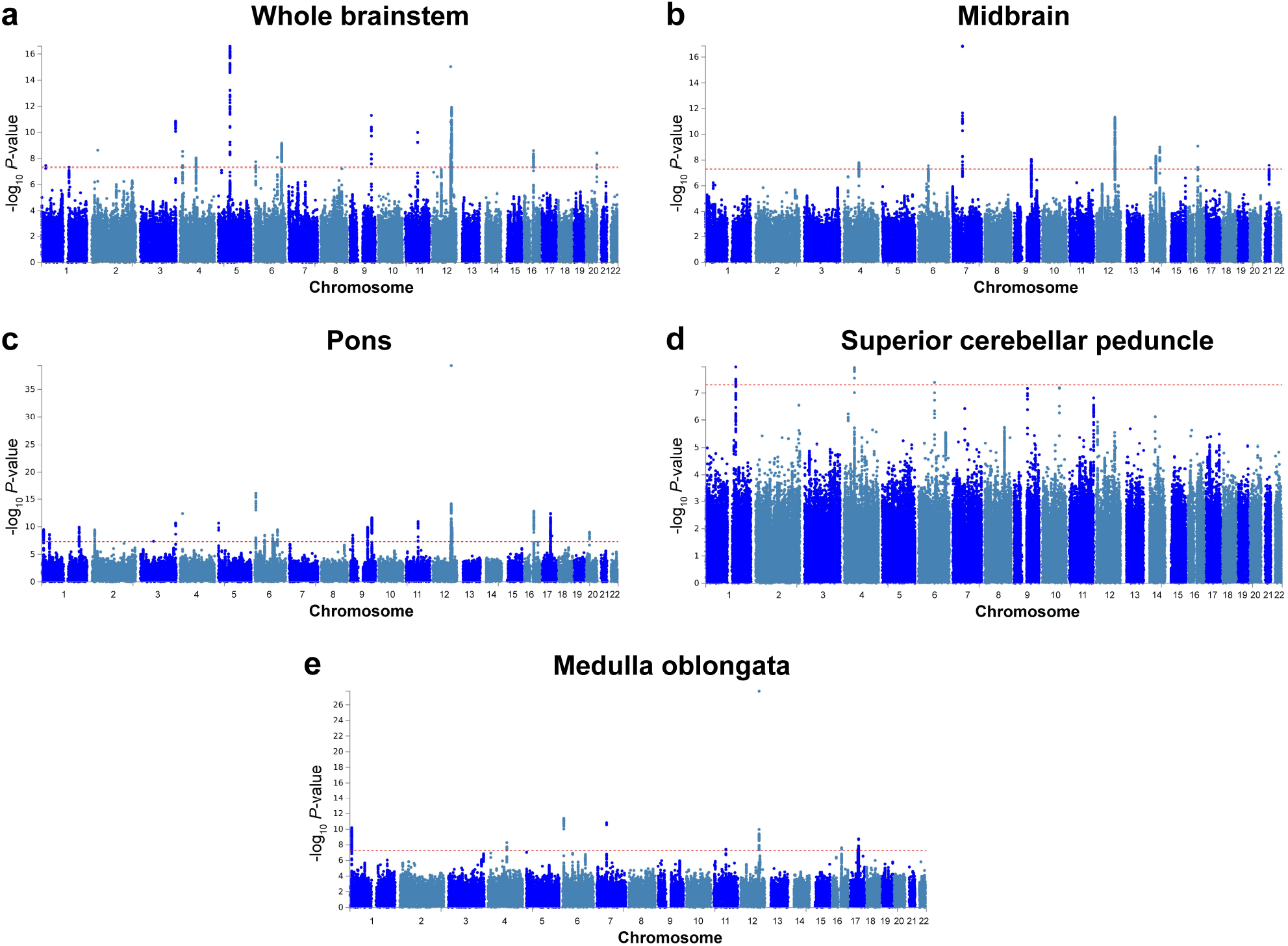
Manhattan plots for volumes of the whole brainstem (**a**), midbrain (**b**), pons (**c**), superior cerebellar peduncle (d), and medulla oblongata (e) from the genome-wide association studies. 16 genetic loci were associated with whole brainstem volume and 10, 23, 3, and 9 loci were associated with volumes of the midbrain, pons, superior cerebellar peduncle, and medulla oblongata, respectively. The red horizontal lines indicate genome-wide significance.

**Supplementary Fig. 3.**
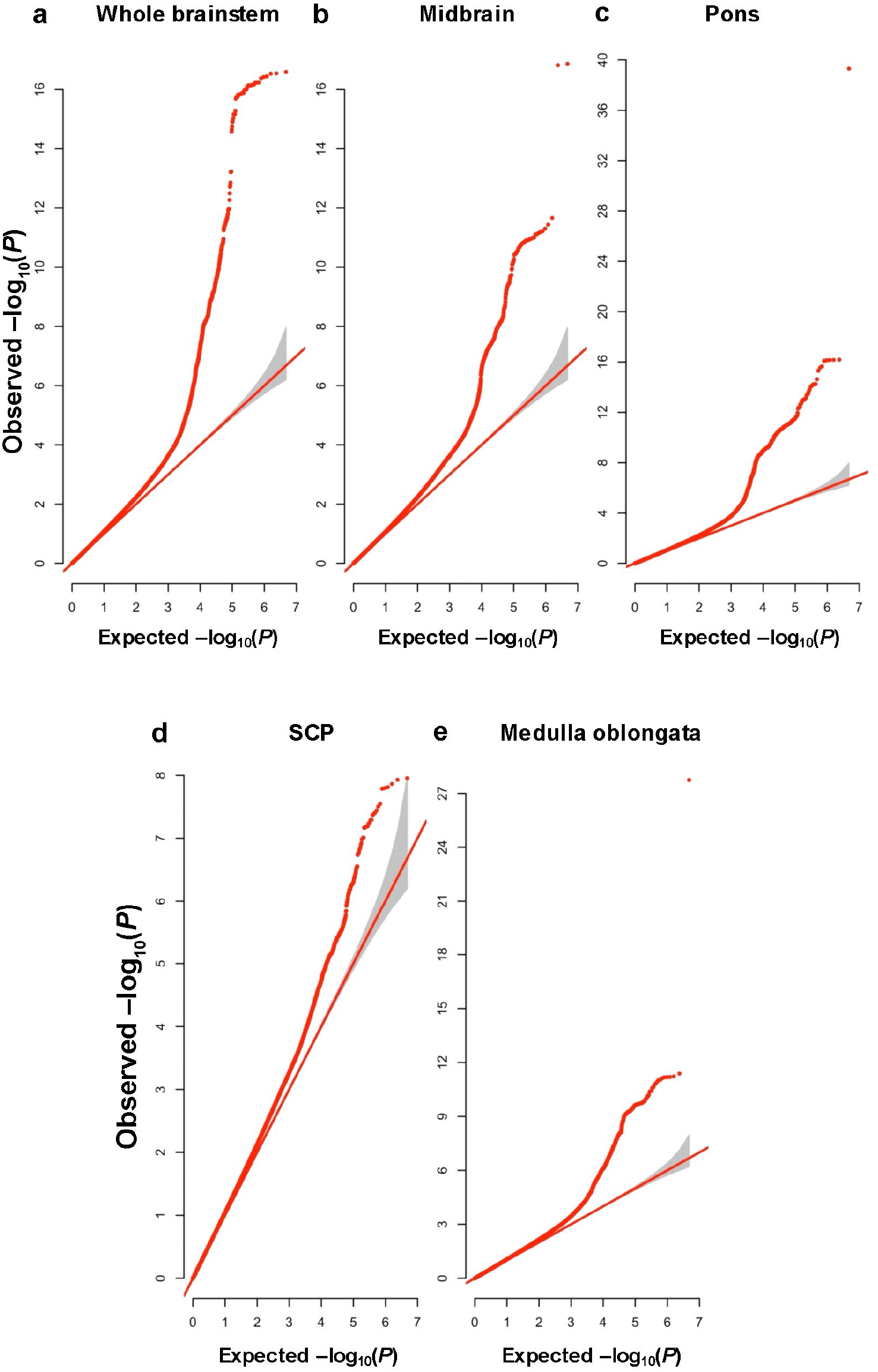
Q-Q plots for volumes of the whole brainstem (**a**), midbrain (**b**), pons (**c**), superior cerebellar peduncle (**d**), and medulla oblongata (**e**) from the genome-wide association studies. SCP; superior cerebellar peduncle.

**Supplementary Fig. 4.**
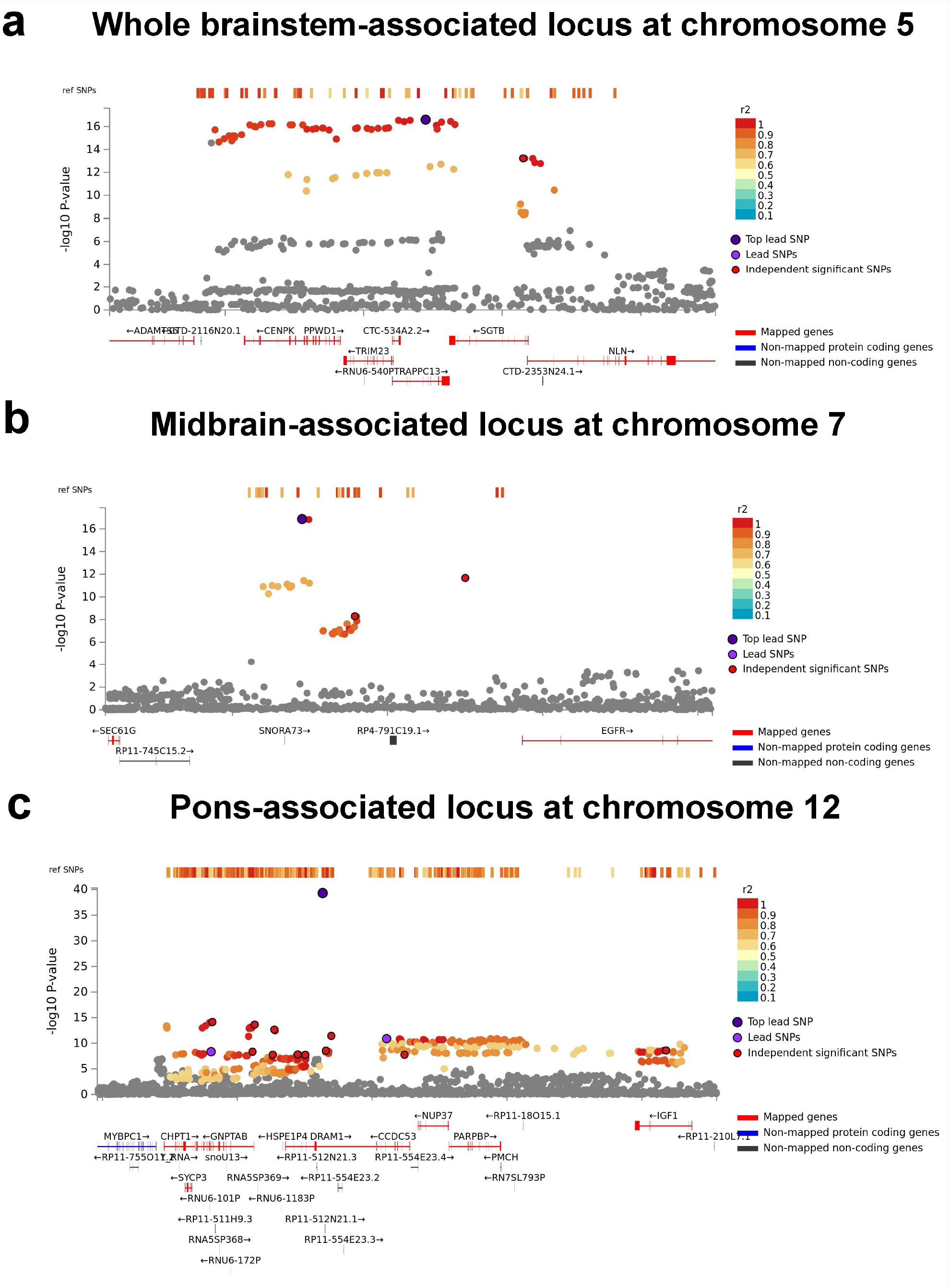
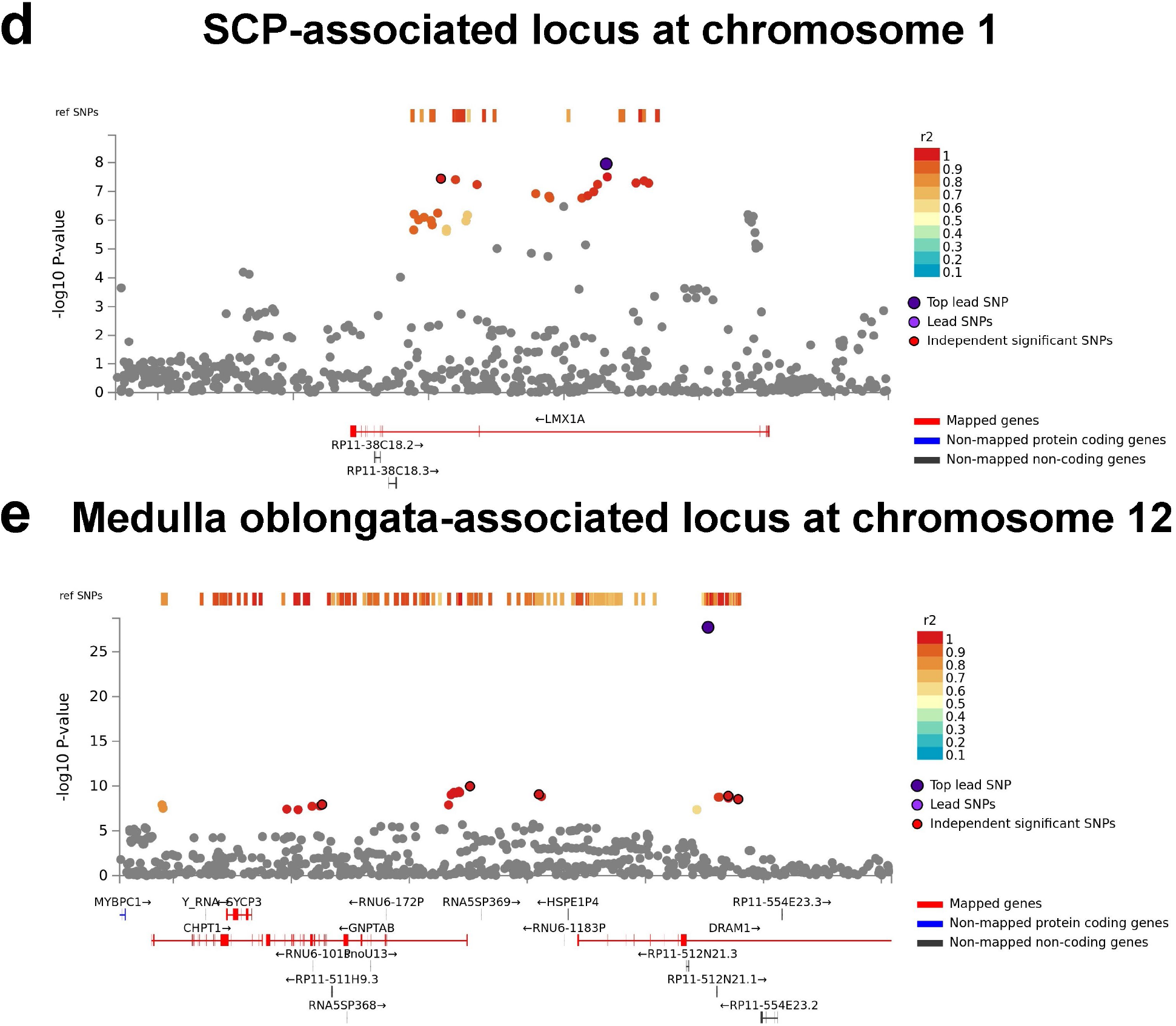
Regional plots for the most significant genetic locus from the genome-wide association studies for volumes of the whole brainstem at chromosome 5 (**a**), midbrain at chromosome 7 (**b**), pons at chromosome 12 (**c**), superior cerebellar peduncle at chromosome 1 (**d**), and medulla oblongata at chromosome 12 (**e**). SCP; superior cerebellar peduncle.

**Supplementary Fig. 5.**
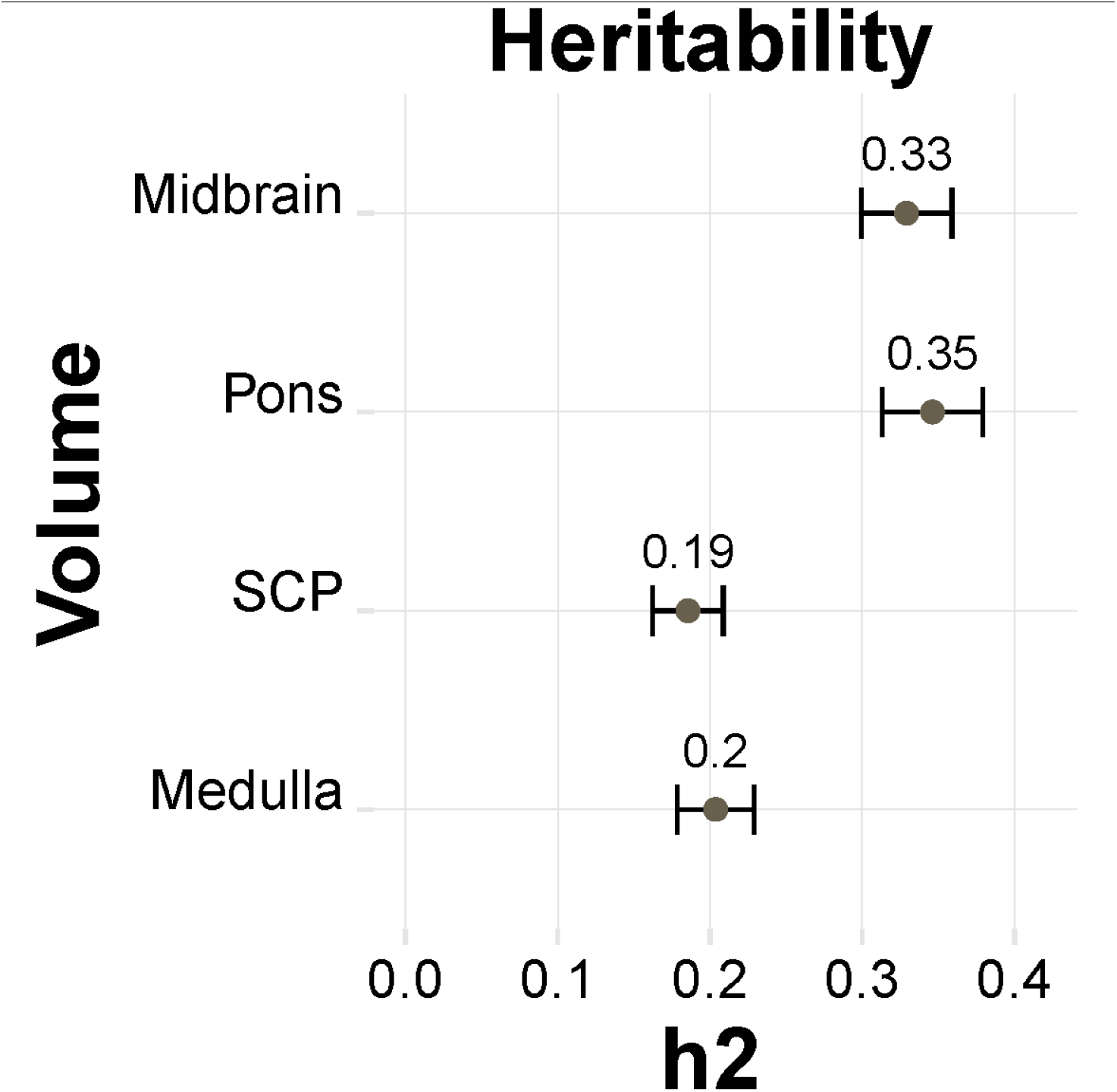
Heritability estimates for the brainstem volumes of *n* = 27,034 healthy individuals when not accounting for the whole brainstem. All brainstem volumes showed substantial heritability, which highest estimates for the midbrain (*h_2_ =* 0.33) and pons (*h_2_ =* 0.35) and lowest for the medulla oblongata (*h_2_ =* 0.20) and SCP (*h_2_ =* 0.19). SCP; superior cerebellar peduncle.

**Supplementary Fig. 6.**
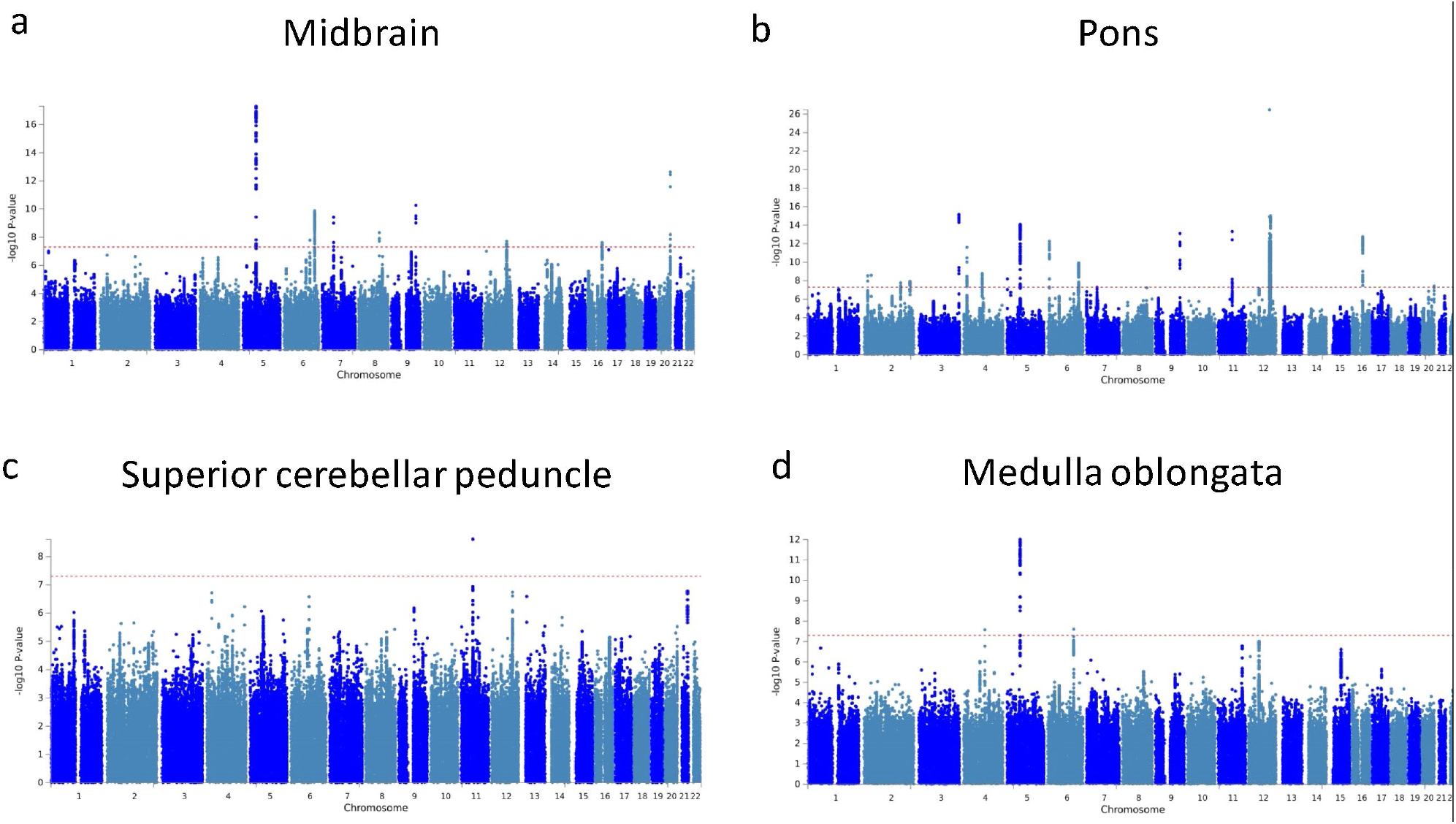
Manhattan plots for volumes of the midbrain (**a**), pons (**b**), superior cerebellar peduncle (**c**), and medulla oblongata (**d**) from the genome-wide association studies when not accounting for whole brainstem volume. The red horizontal lines indicate genome-wide significance. Additional information for the significant loci is provided in Supplementary Table 5.

**Supplementary Fig. 7.**
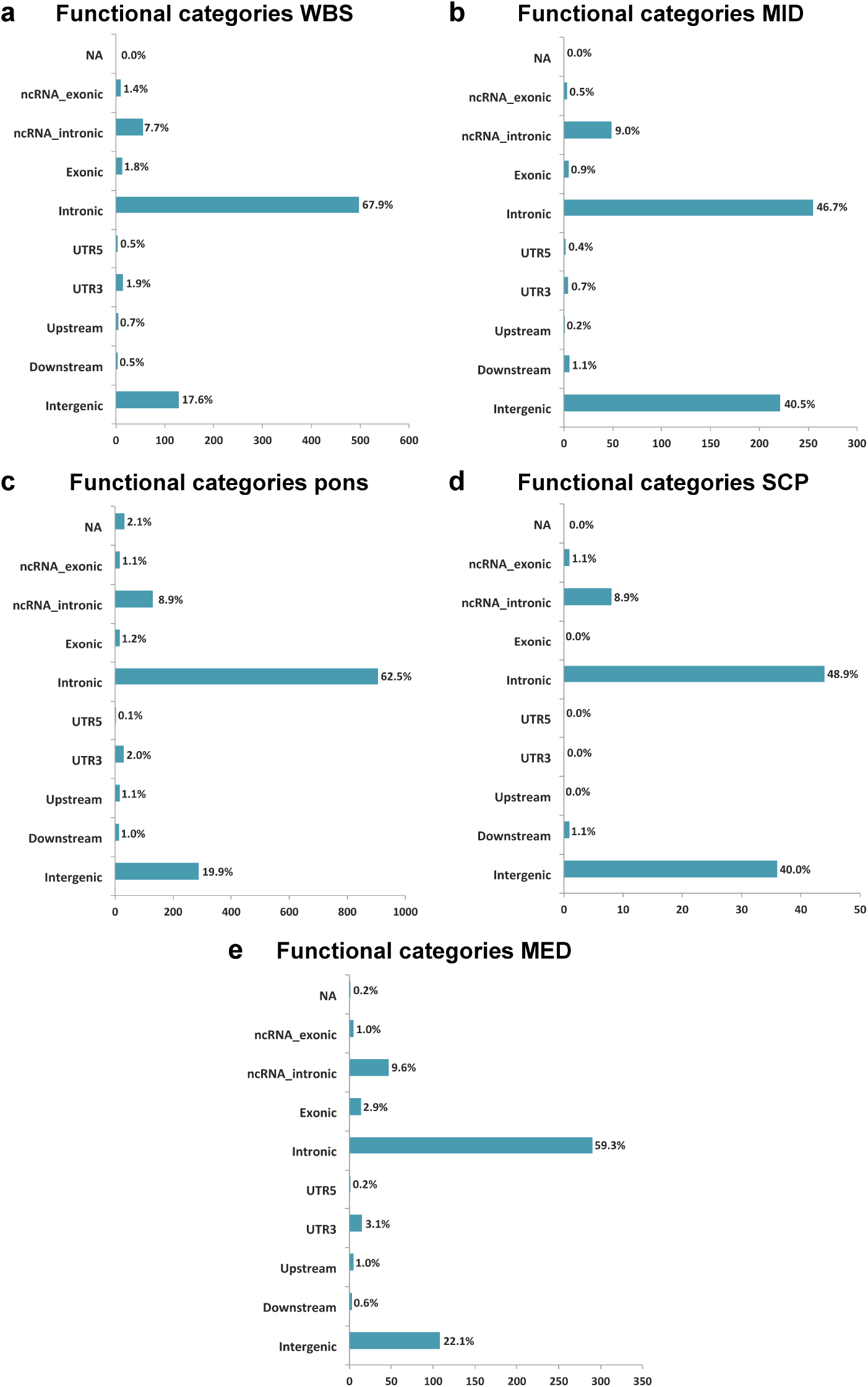
Functional single-nucleotide polymorphism categories from the genome-wide association studies for volumes of the whole brainstem (**a**), midbrain (**b**), pons (**c**), superior cerebellar peduncle (**d**), and medulla oblongata (**e**). WBS; whole brainstem. MID; midbrain. SCP; superior cerebellar peduncle. MED; medulla oblongata. UTR; untranslated region.

**Supplementary Fig. 8.**
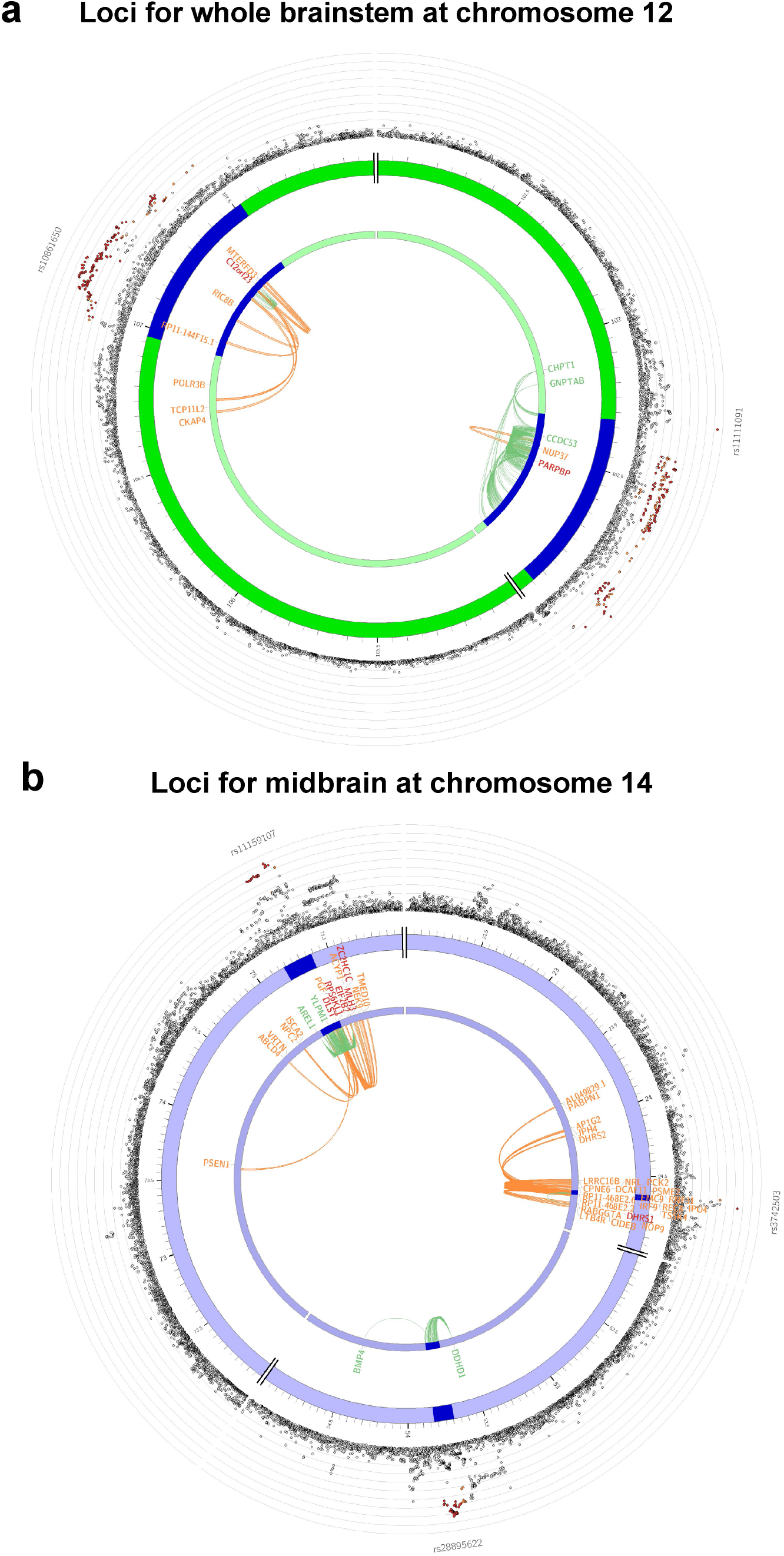
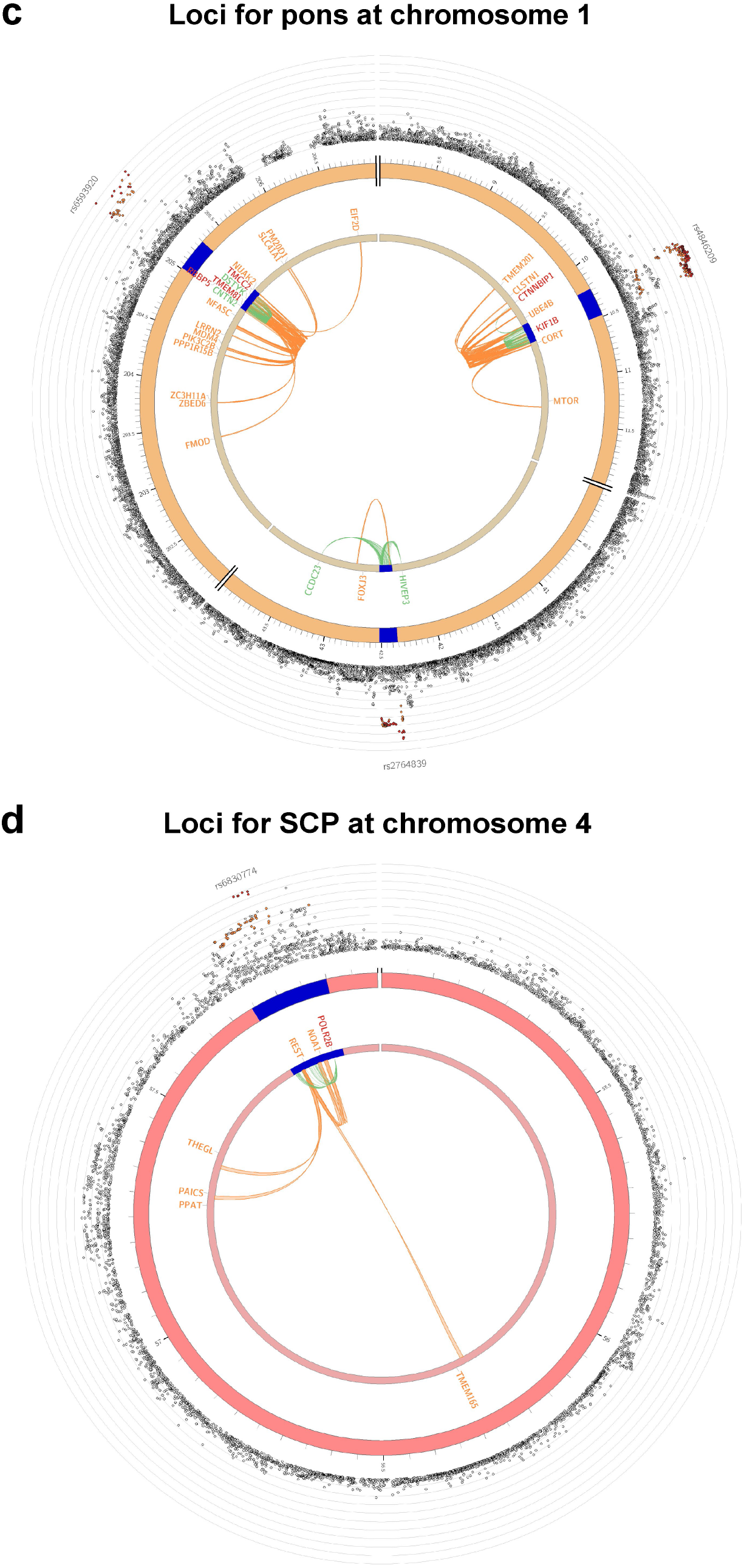
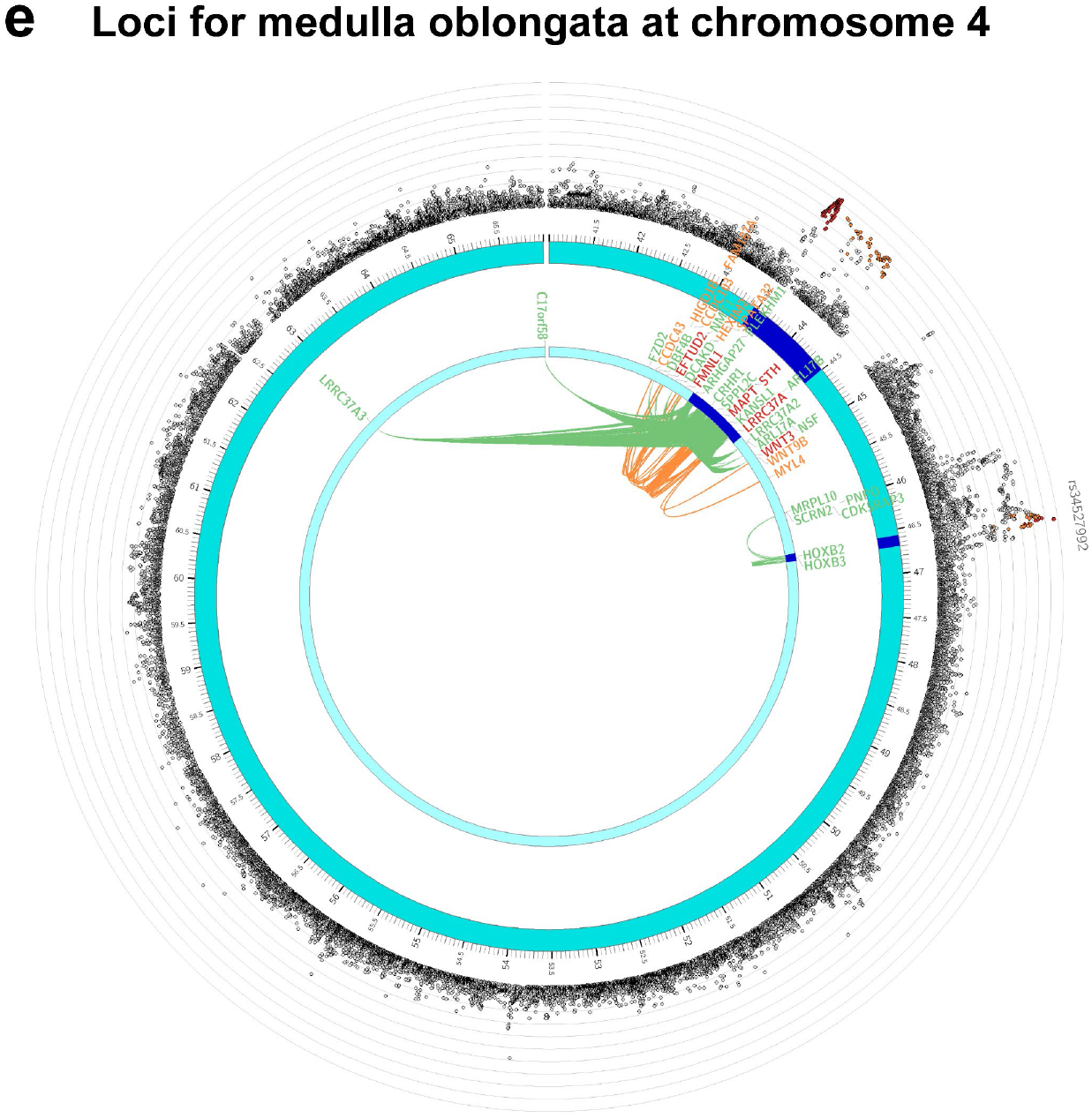
Examples of Circos plots of mapped genes for the whole brainstem at chromosome 12 (**a**), midbrain at chromosome 14 (**b**), pons at chromosome 1 (**c**), superior cerebellar peduncle at chromosome 4 (**d**), and medulla oblongata at chromosome 4 (**e**). The plots show mapped genes of significant genetic loci from the genome-wide association studies of brainstem volumes (blue regions). The genes were linked to the loci by eQTL mapping (green lines) and chromatin interactions (orange lines). Green color indicates genes implicated by eQTLs, orange color indicates genes mapped by chromatin interactions, and genes implicated by both strategies are in red color. The outer layers show the Manhattan plots of single nucleotide polymorphisms from the genome-wide association studies.

**Supplementary Fig. 9.**
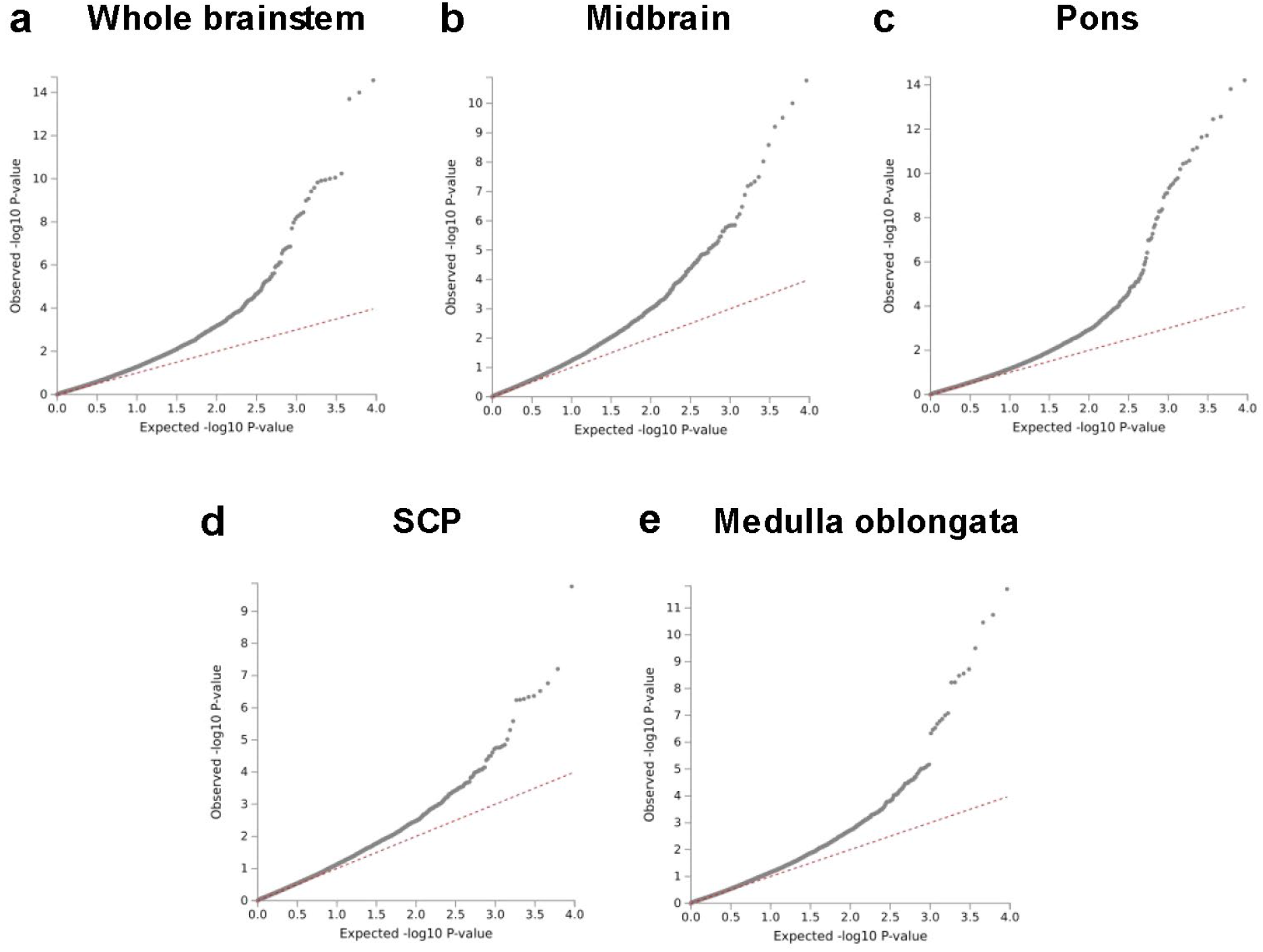
Q-Q plots for volumes of the whole brainstem (**a**), midbrain (**b**), pons (**c**), superior cerebellar peduncle (**d**), and medulla oblongata (**e**) from the genome-wide gene-based association analyses. SCP; superior cerebellar peduncle.

**Supplementary Fig. 10.**
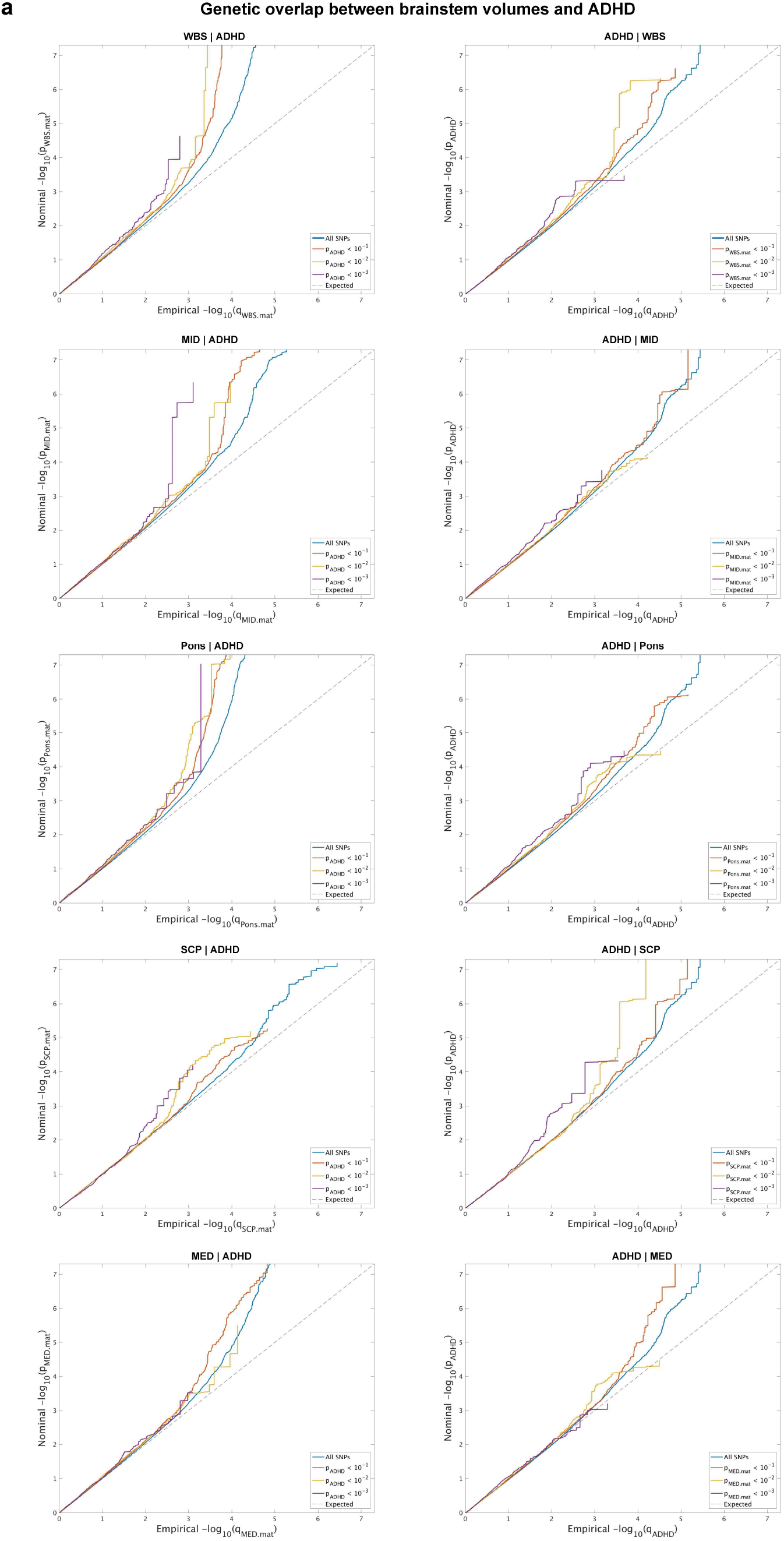
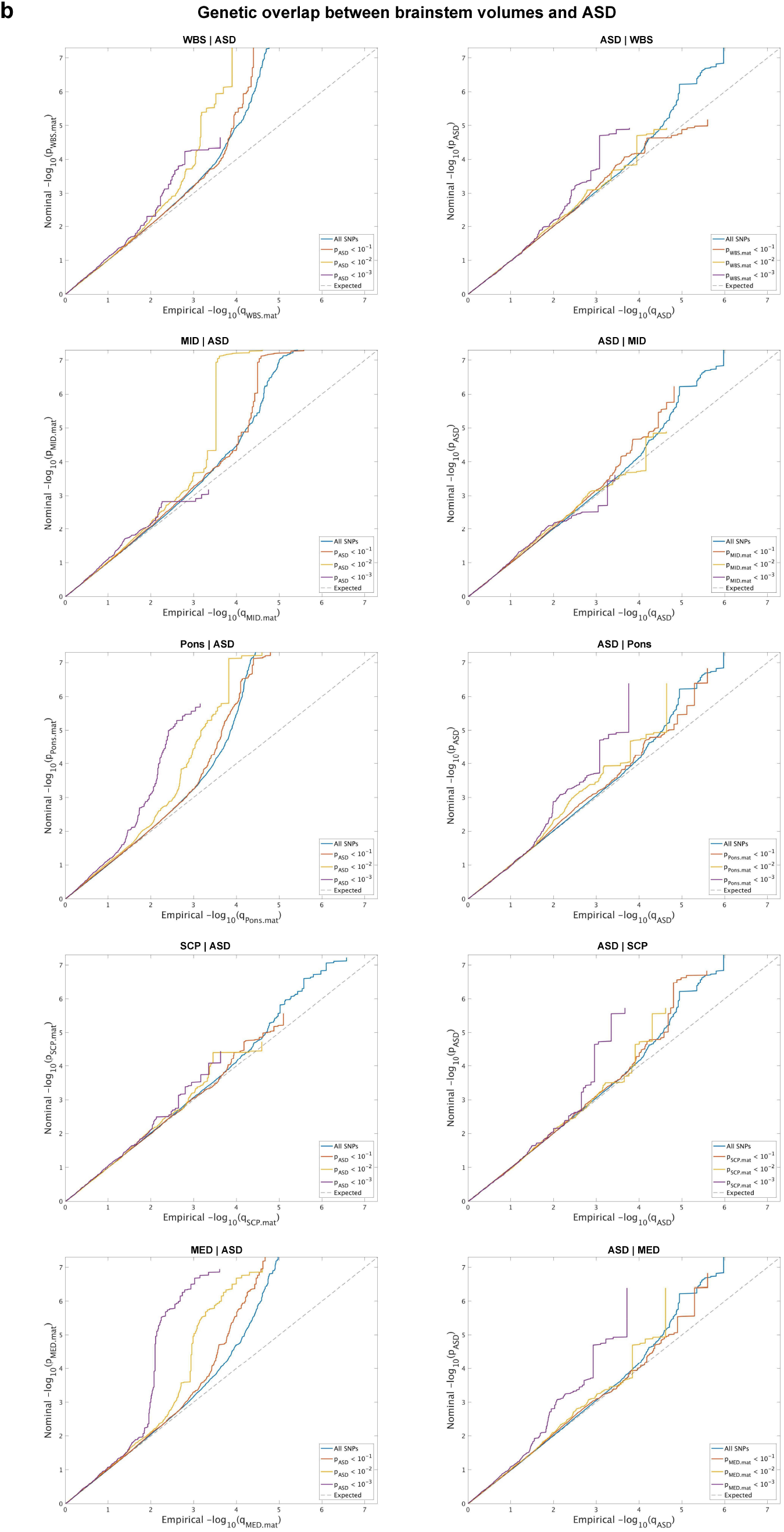
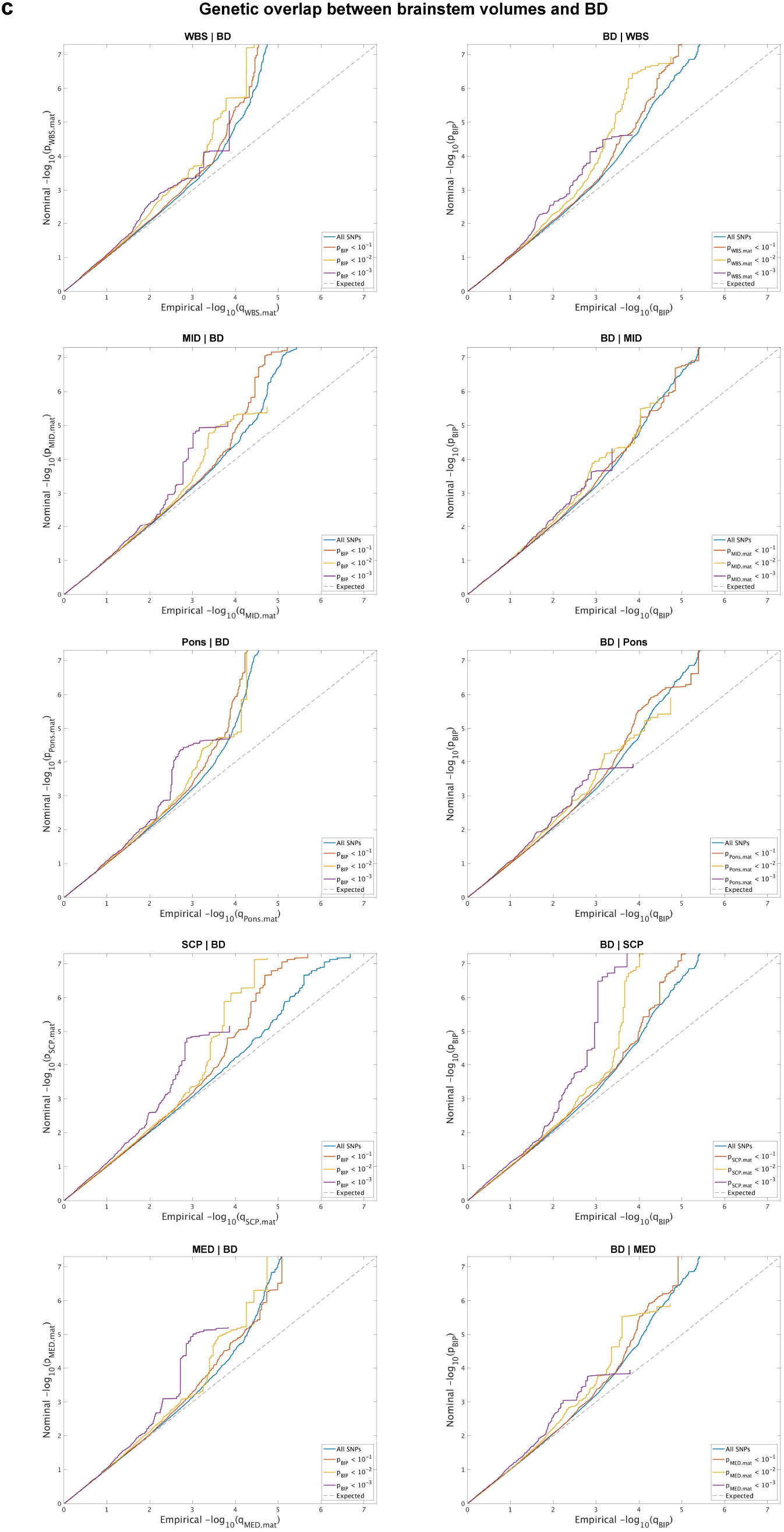
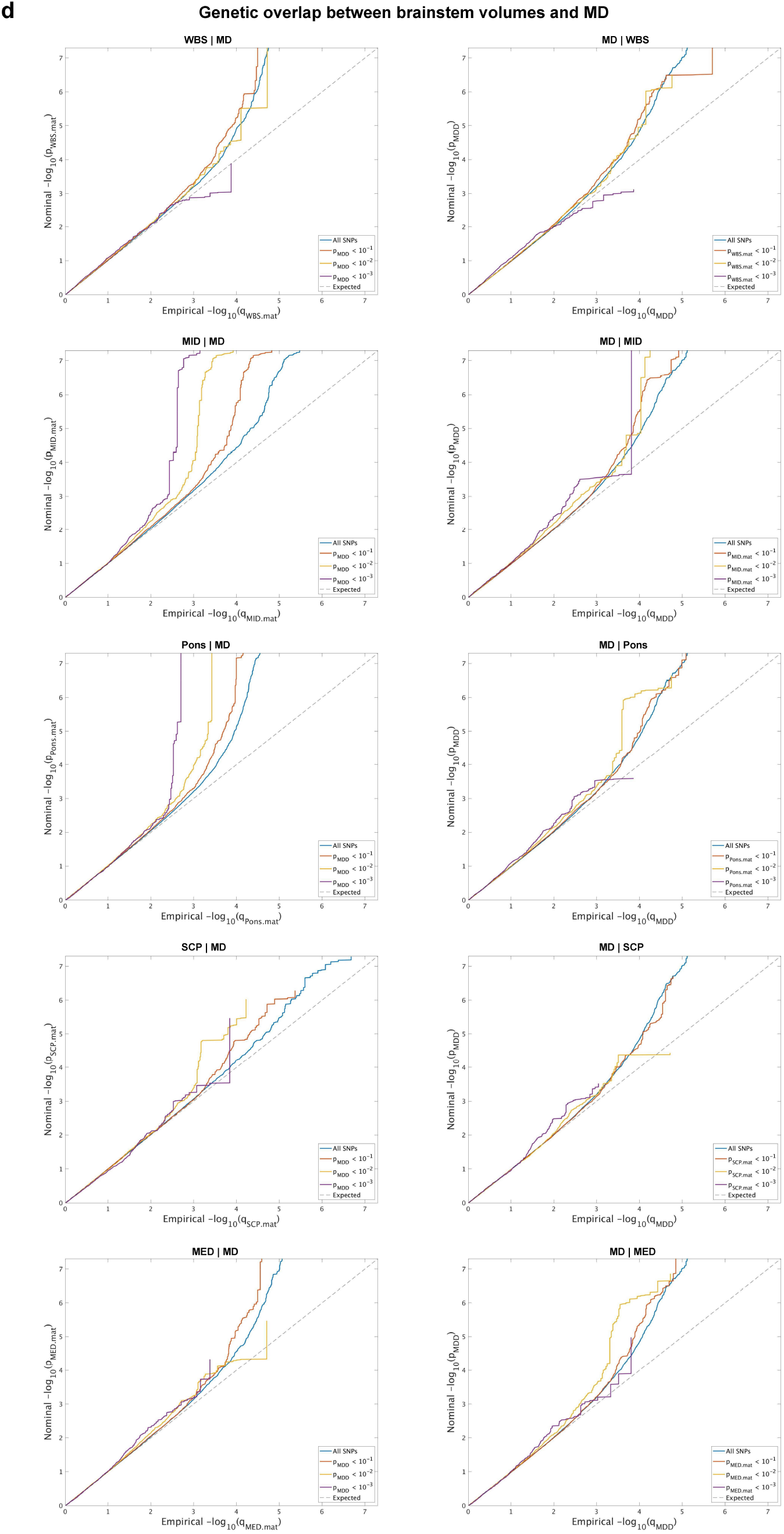
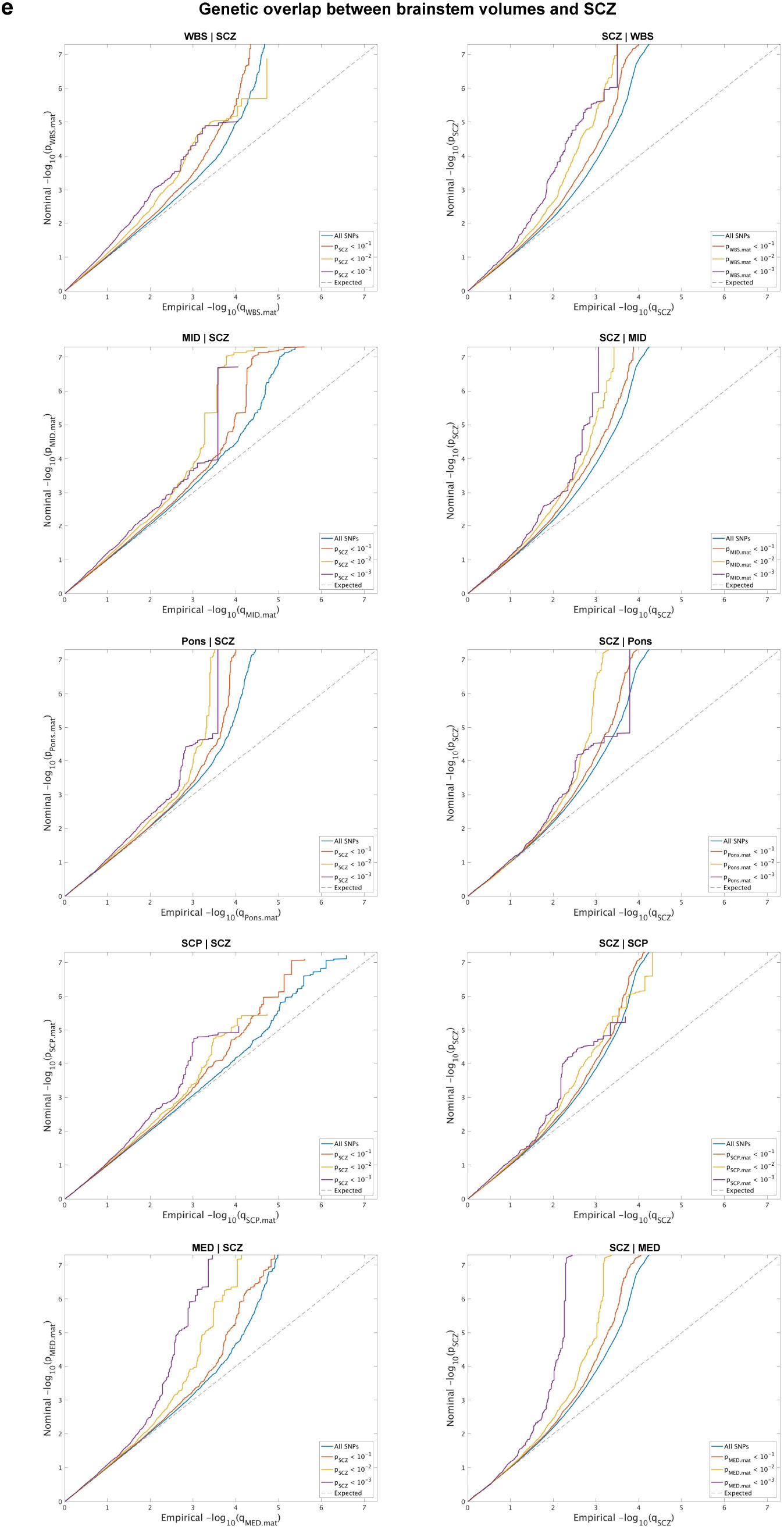
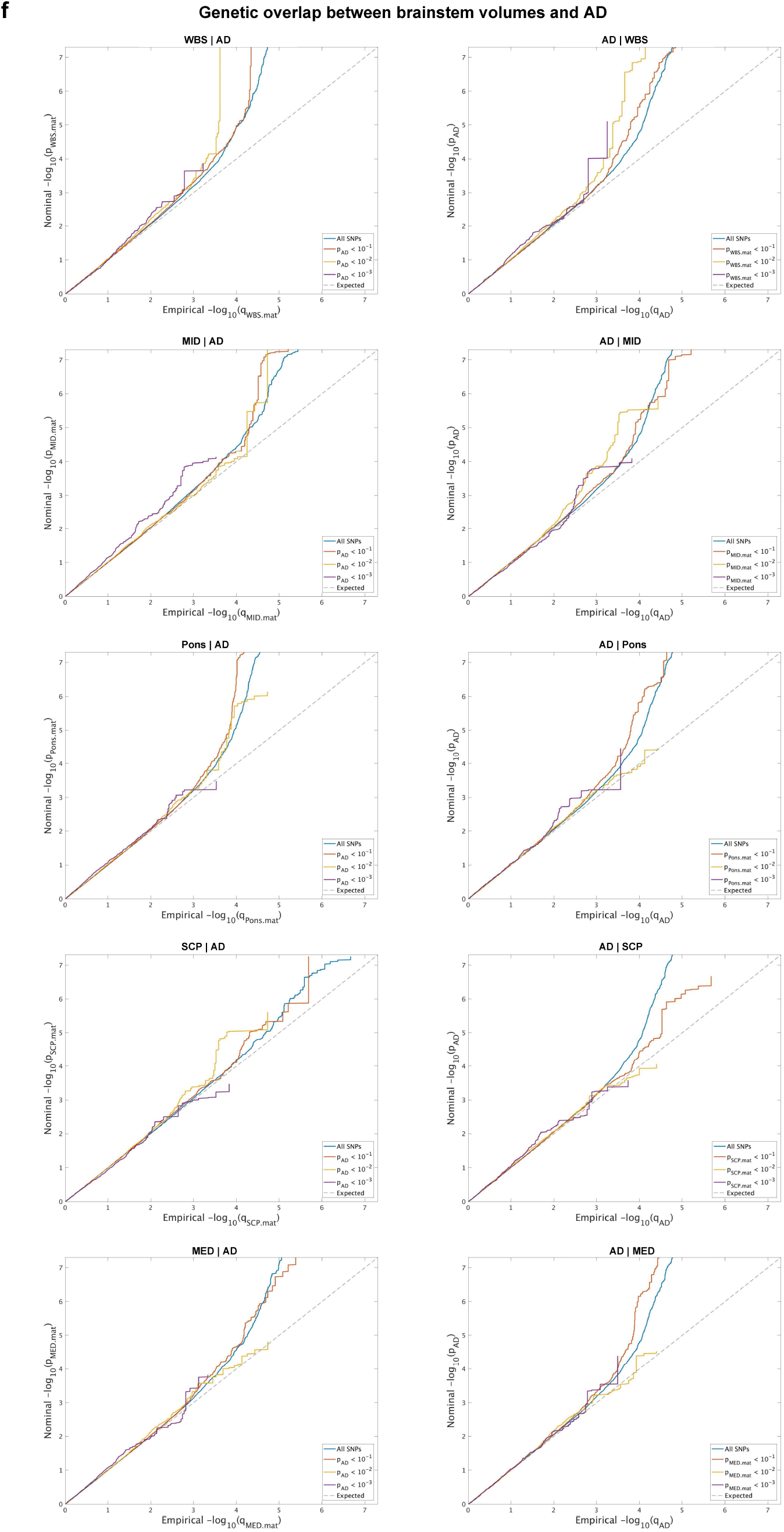
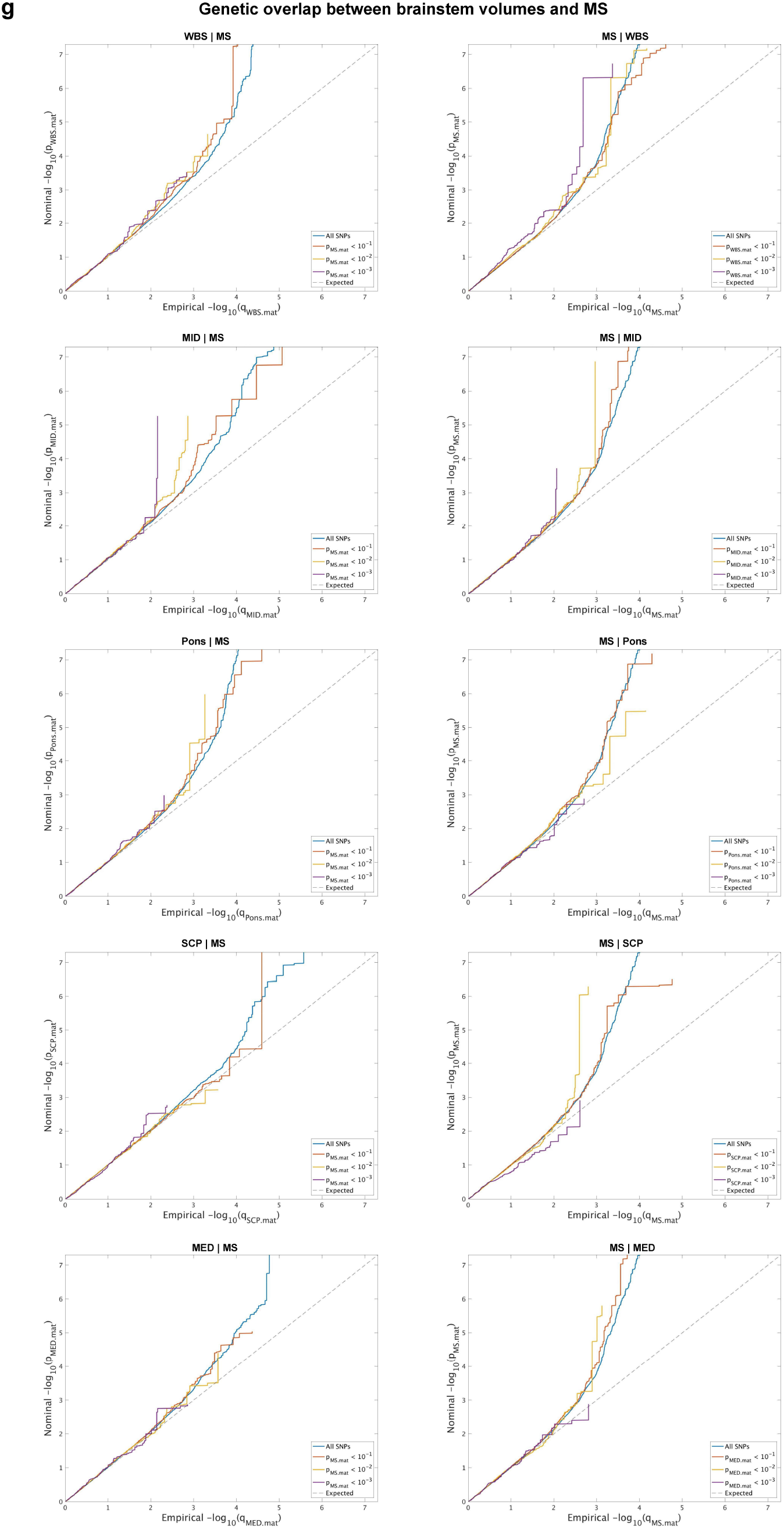
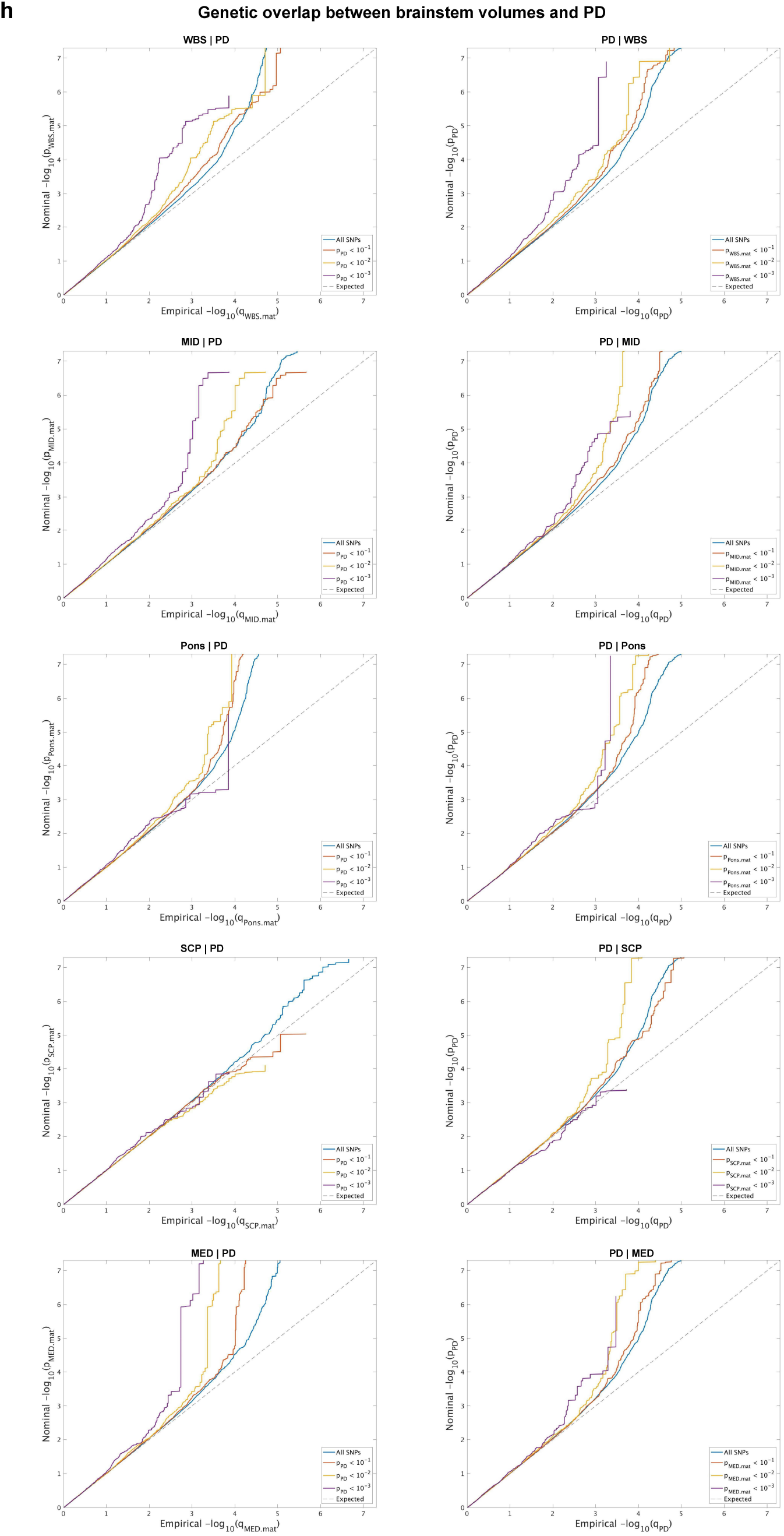
Conditional Q-Q plots for brainstem volumes given associations with the disorder (left figures) and vice versa (right figures), for attention deficit hyperactivity disorder (**a**), autism spectrum disorder (**b**), bipolar disorder (**c**), major depression (**d**), schizophrenia (**e**), Alzheimer’s disease (**f**), multiple sclerosis (**g**), and Parkinson’s disease (**h**). ASD; autism spectrum disorders. BD; bipolar disorder. MD; major depression. SCZ; schizophrenia. AD; Alzheimer’s disease. MS; multiple sclerosis. PD; Parkinson’s disease.

**Supplementary Fig. 11.**
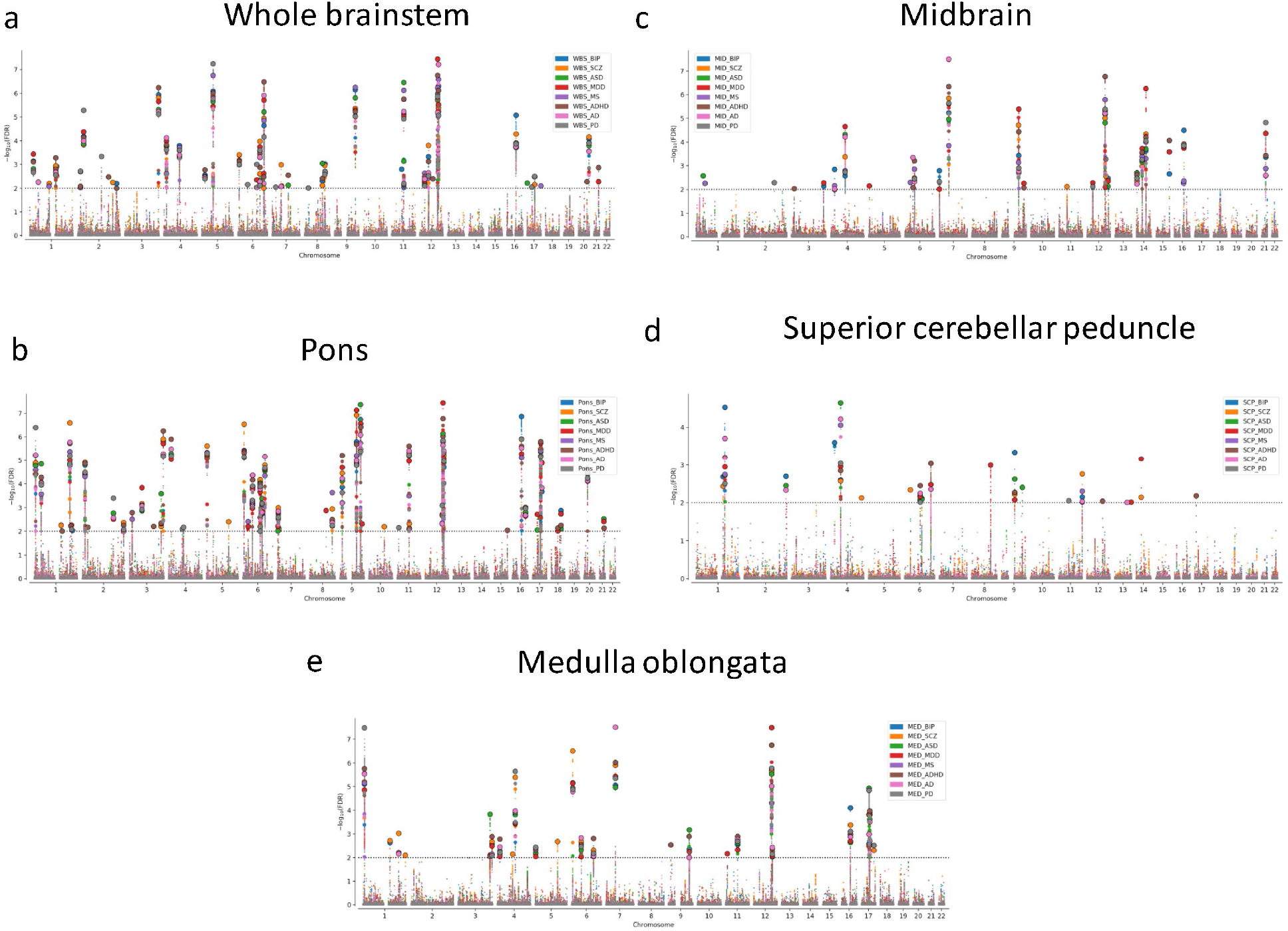
Manhattan plots of genetic loci for each brainstem region identified by the condition false discovery rate analyses when conditioned on the eight brain disorders. These analyses revealed a total of 208 independent significant singlenucleotide polymorphisms (SNPs) for whole brainstem volume (**a**), 111 SNPs for midbrain volume (**b**), 270 SNPs for pons volume (**c**), 55 SNPs for superior cerebellar peduncle volume (**d**), and 125 SNPs for medulla oblongata volume (**e**). ASD; autism spectrum disorders. BD; bipolar disorder. MD; major depression. SCZ; schizophrenia. AD; Alzheimer’s disease. MS; multiple sclerosis. PD; Parkinson’s disease.

**Supplementary Fig. 12.**
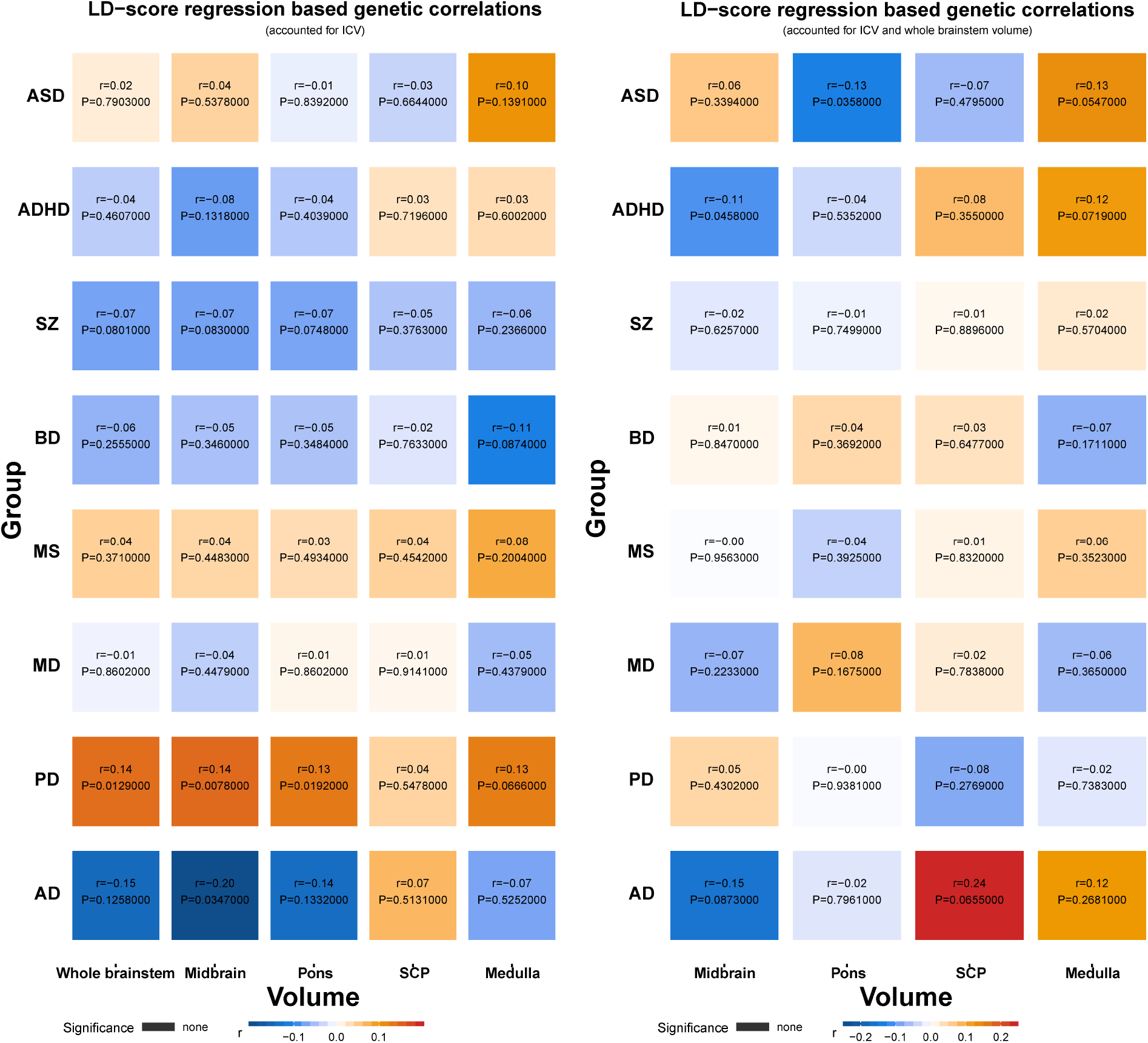
Genetic correlations between brainstem volumes and eight brain disorders. There were correlations between brainstem volumes and ASD, ADHD, and PD with uncorrected *P* < 0.05, yet these were not significant after multiple testing corrections. ADHD; attention deficit hyperactivity disorder. ASD; autism spectrum disorder. BD; bipolar disorder. MD; major depression. MS; multiple sclerosis. PD; Parkinson’s disease. SCZ; schizophrenia.

**Supplementary Fig. 13.**
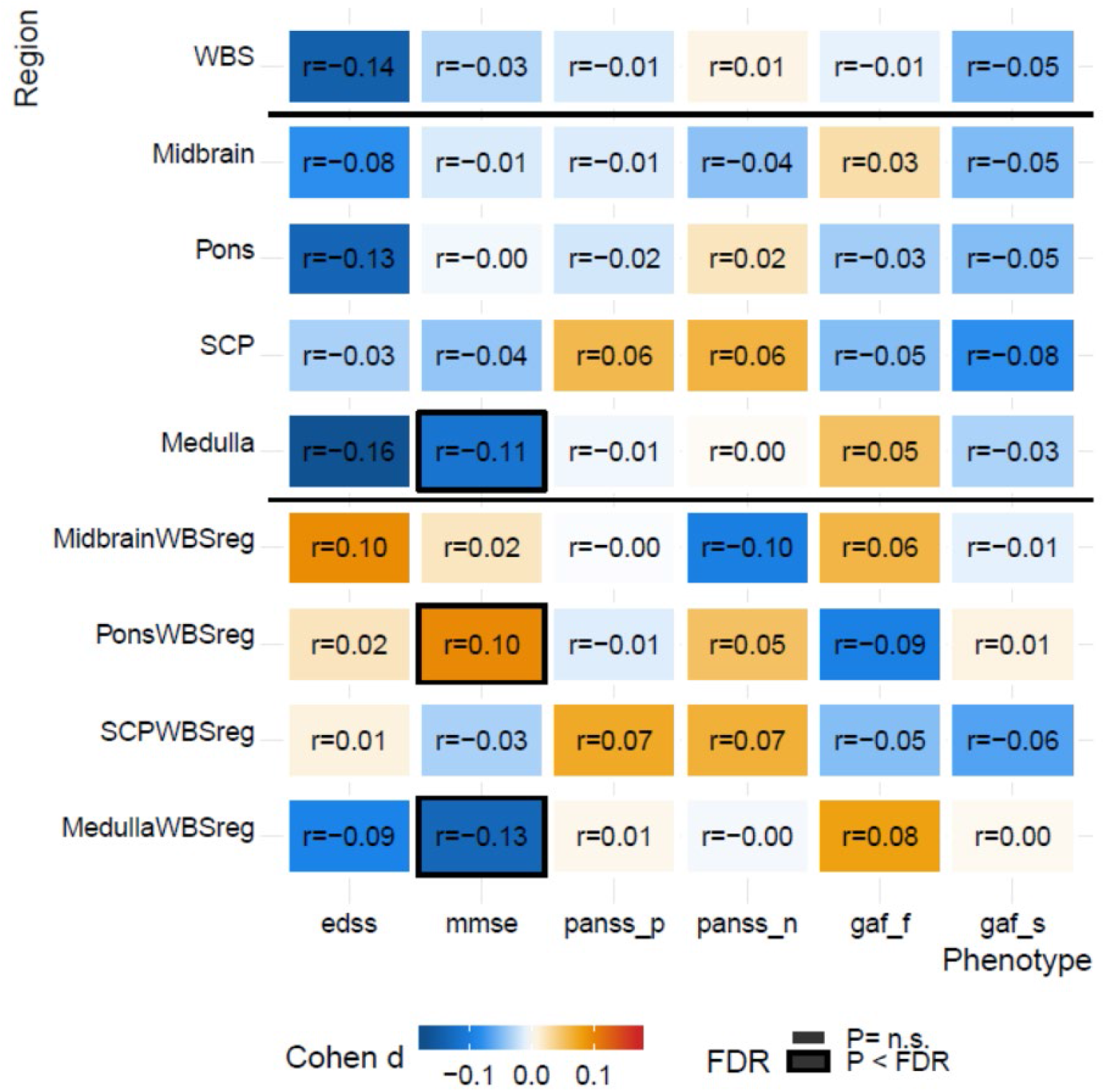
Associations between brainstem volumes and clinical variables. We ran analyses of associations between brainstem volumes and clinical characteristics in the individuals with MCI, DEM, MS, SCZ, and PD. Across individuals with MCI and DEM (*n* = 1610), there were negative associations between Mini-Mental State Examination (MMSE) scores^40^ and medulla oblongata volume before (*r* = −0.11, *P =* 1.8e-05) and after (*r* = −0.13, *P =* 3.5e-07) accounting for whole brainstem volume. In addition, there was a significant positive association between MMSE and pons volume when adjusted for the whole brainstem (*r* = 0.10, *P =* 1.7e-04). All MRIs from the individuals with MS were examined by two neuroradiologists and then divided into two groups according to presence of infratentorial lesions. There was no significant difference in brainstem volumes between patients with (*n* = 153) and without (*n* = 91) infratentorial lesions (all *P* > 0.05; results not shown in the figure). Patients without lesions had reduced volumes relative to the controls of the whole brainstem (Cohen’s *d* = −0.23, *P* = 0.03), midbrain (Cohen’s *d* = - 0.26, *P* = 0.01), and medulla oblongata (Cohen’s *d* = −0.22, *P* = 0.03; results not shown in the figure). There were significant reductions in volumes of patients with infratentorial lesions relative to controls for the whole brainstem (Cohen’s *d* = - 0.30, *P* = 3.4e-04), the midbrain (Cohen’s *d* = −0.36, *P* = 1.9e-05), the pons (Cohen’s *d* = −0.24, *P* = 3.9e-03), and medulla

**Supplementary Fig. 14.**
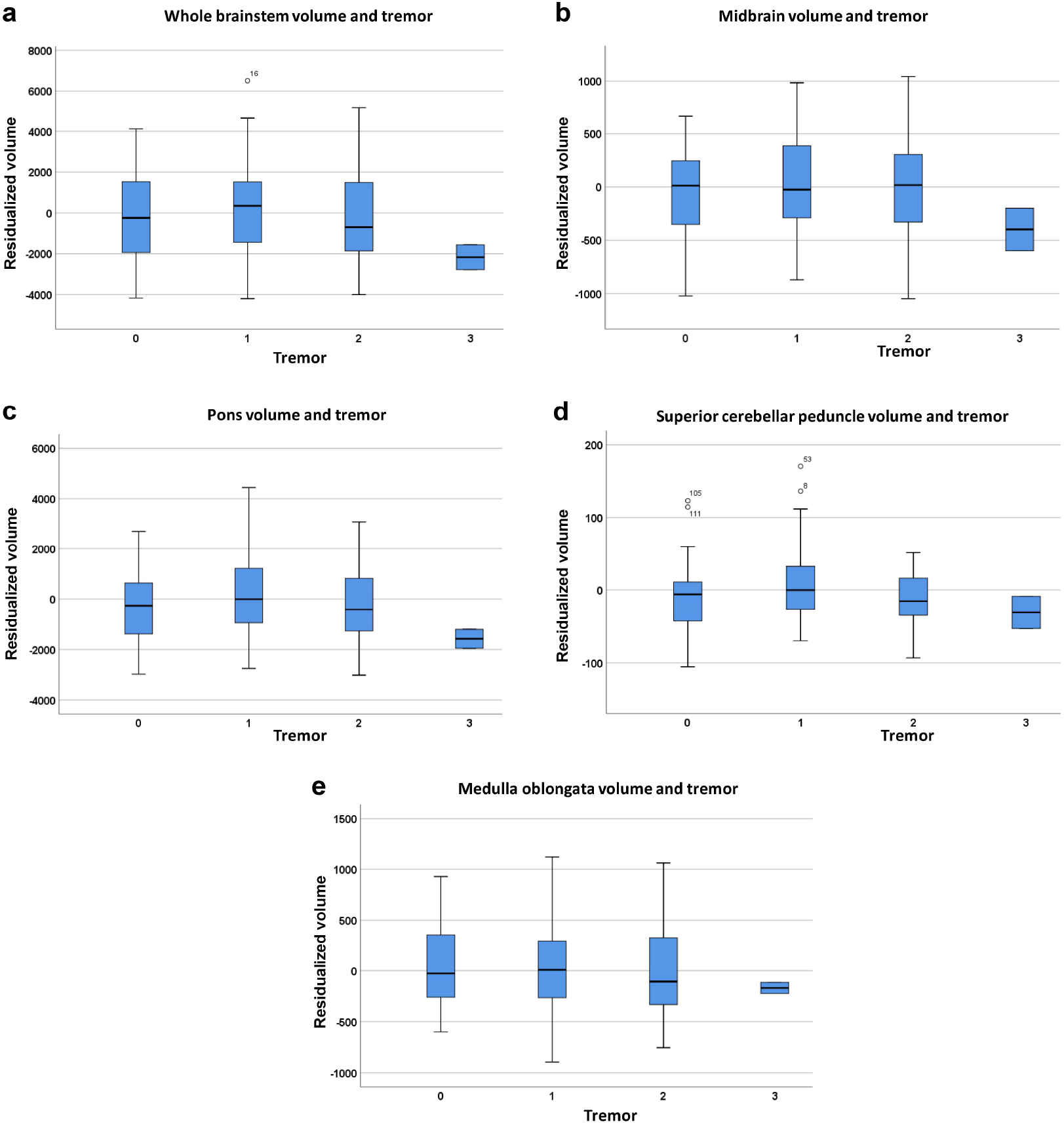
Tremor and brainstem volumes in individuals with Parkinson’s disease (PD).We examined whether the volume increases in the individuals with PD were related to tremor, which could cause increased within-scanner motion and confound the brainstem segmentation. Here, we used item 2.10 of the Unified Parkinson’s Disease Rating Scale III^75^: “Over the past week, have you usually had shaking or tremor? 0: Normal: Not at all. I have no shaking or tremor; 1: Slight: Shaking or tremor occurs but does not cause problems with any activities.; 2: Mild: Shaking or tremor causes problems with only a few activities; 3: Moderate: Shaking or tremor causes problems with many of my daily activities; and 4: Severe: Shaking or tremor causes problems with most or all activities.” Thirty individuals had a tremor score of 0, 74 individuals had a score of 1, 22 individuals had a score of 2, 2 individuals had a score of 3, and none had a score of 4. We then grouped the individuals according to the tremor level and compared brainstem volumes between these groups using a linear model, covarying for gender, intracranial volume, scanner, age, and age^2^. There were no significant effects of tremor group on brainstem volumes (all *P* > 0.13).

## Supplementary Tables

**Supplementary Table 1:**
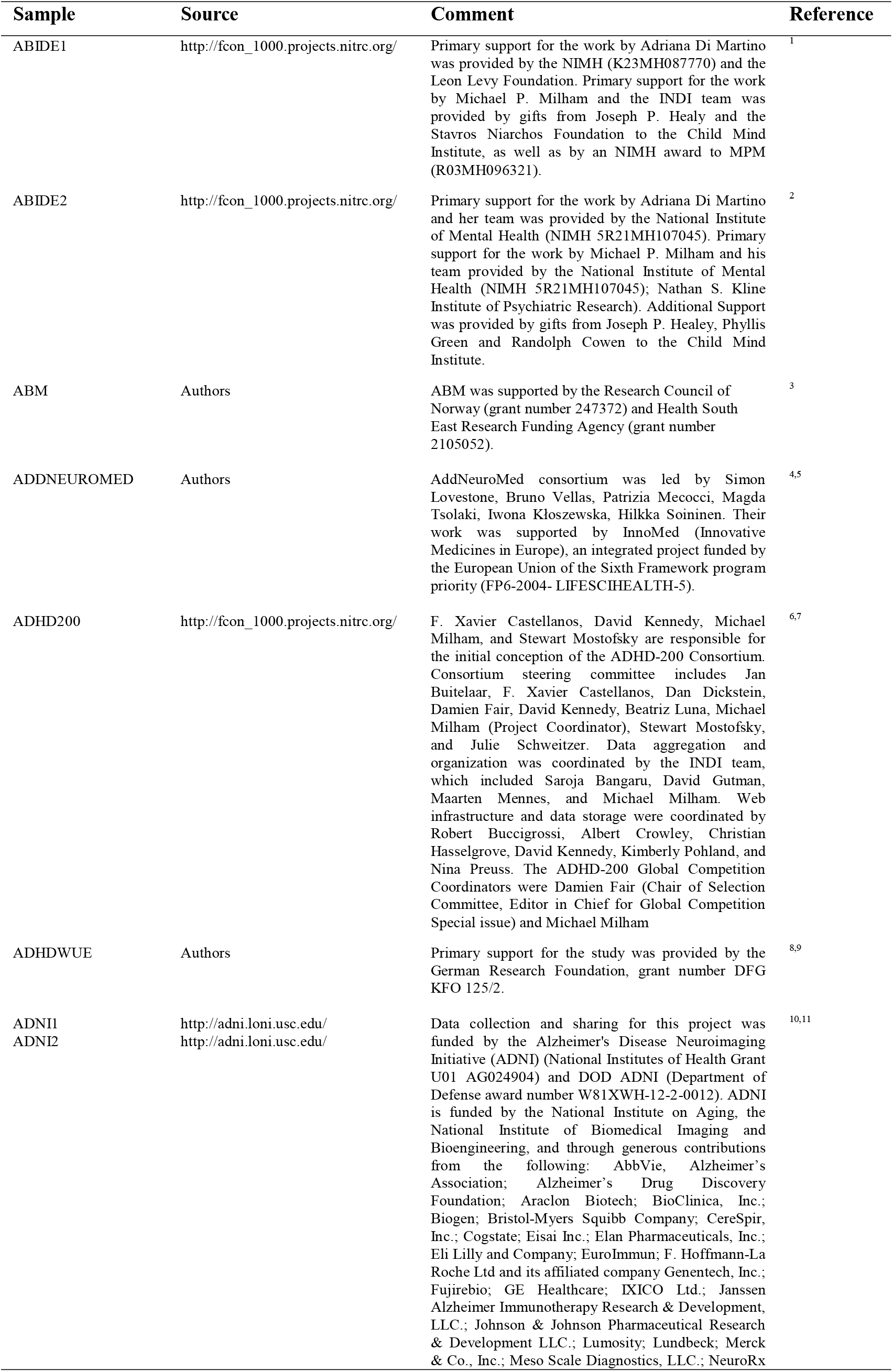
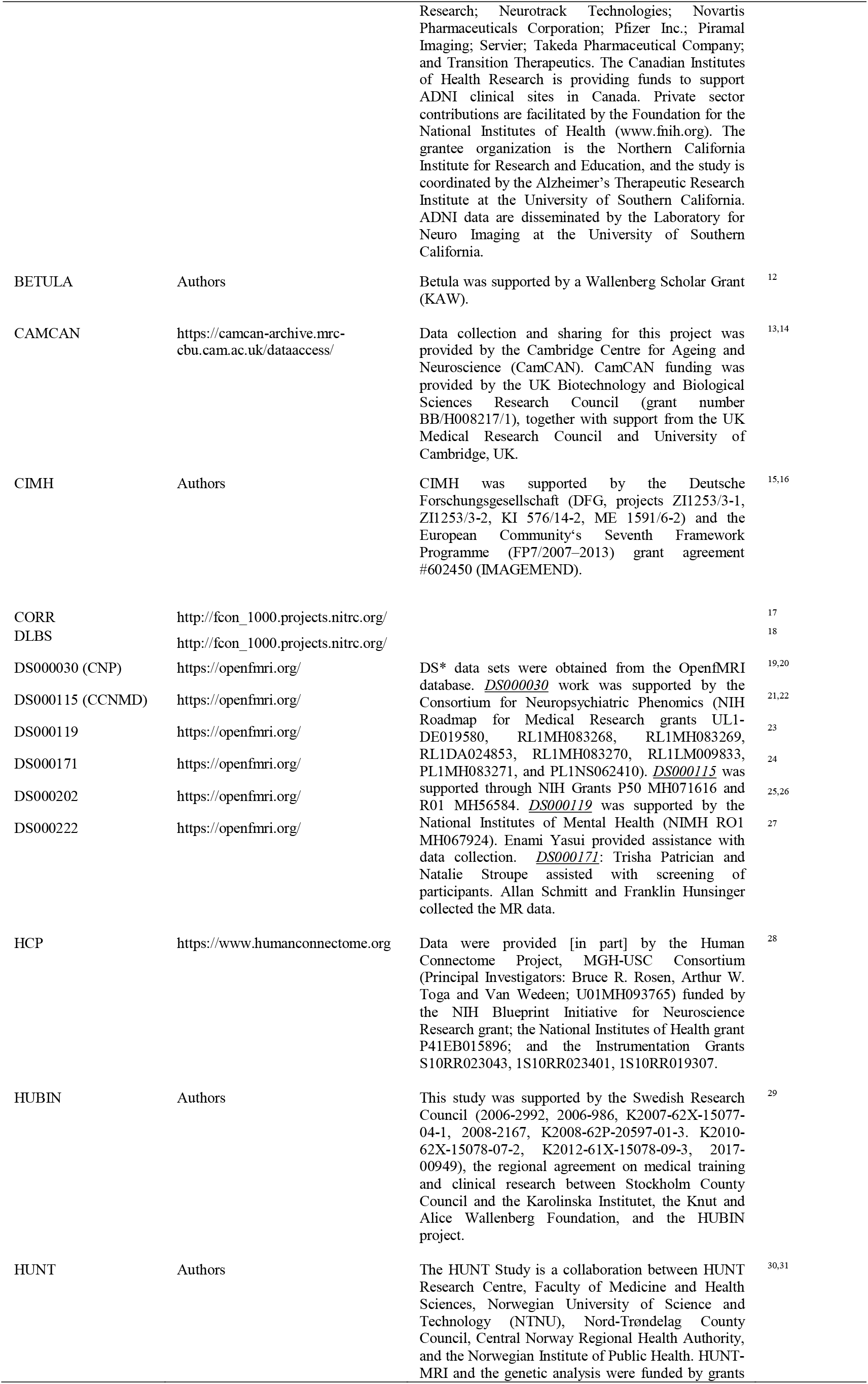
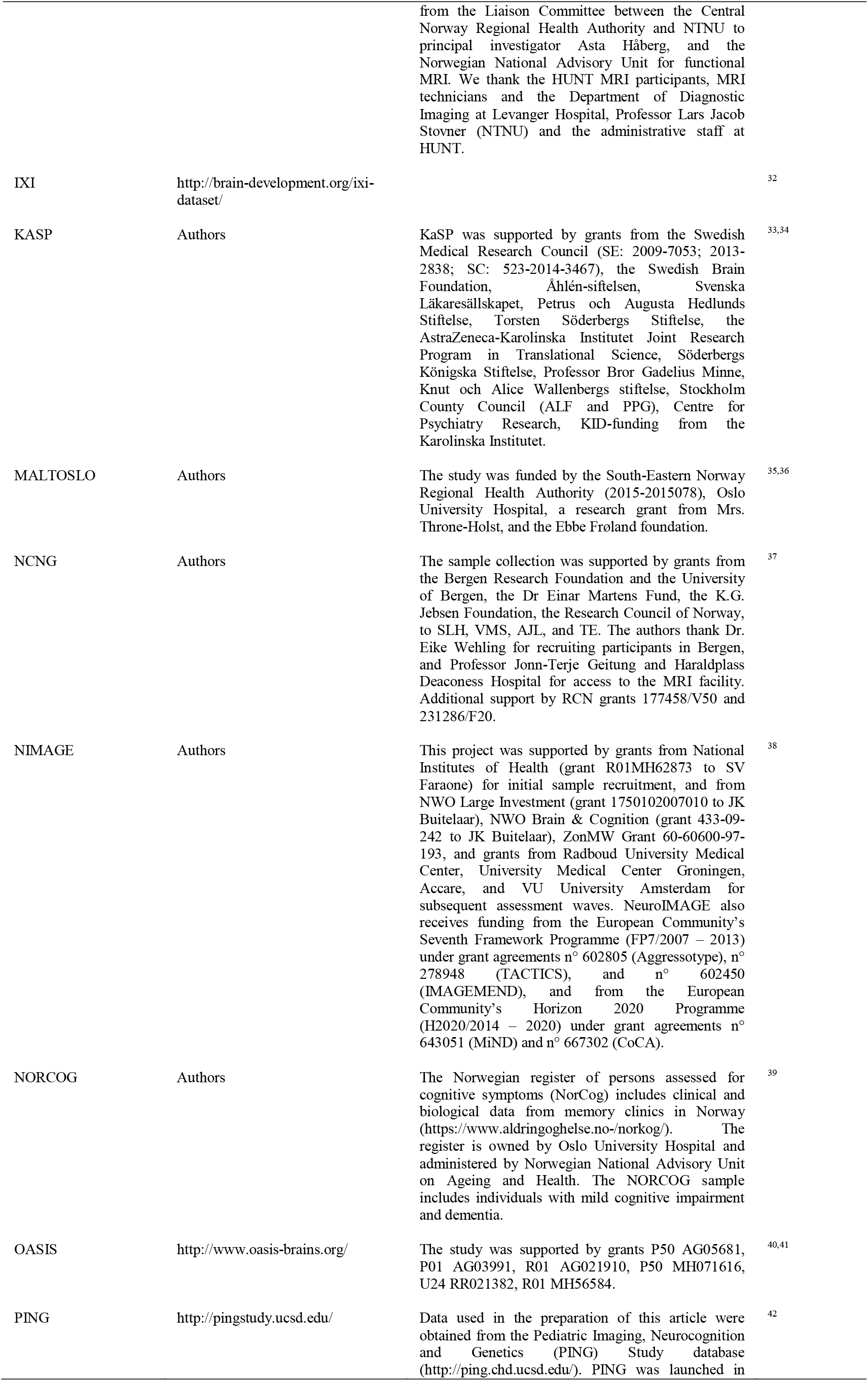
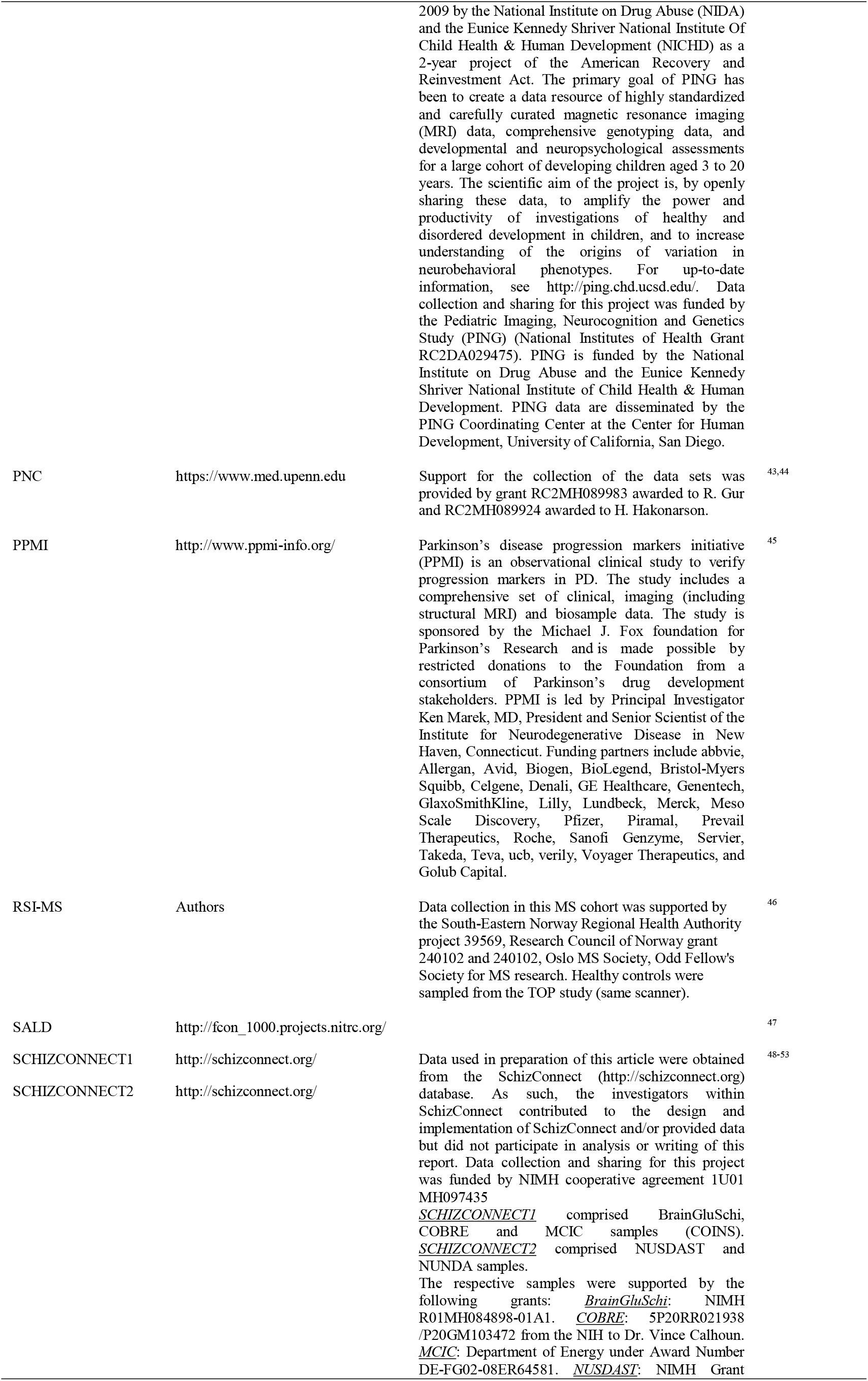
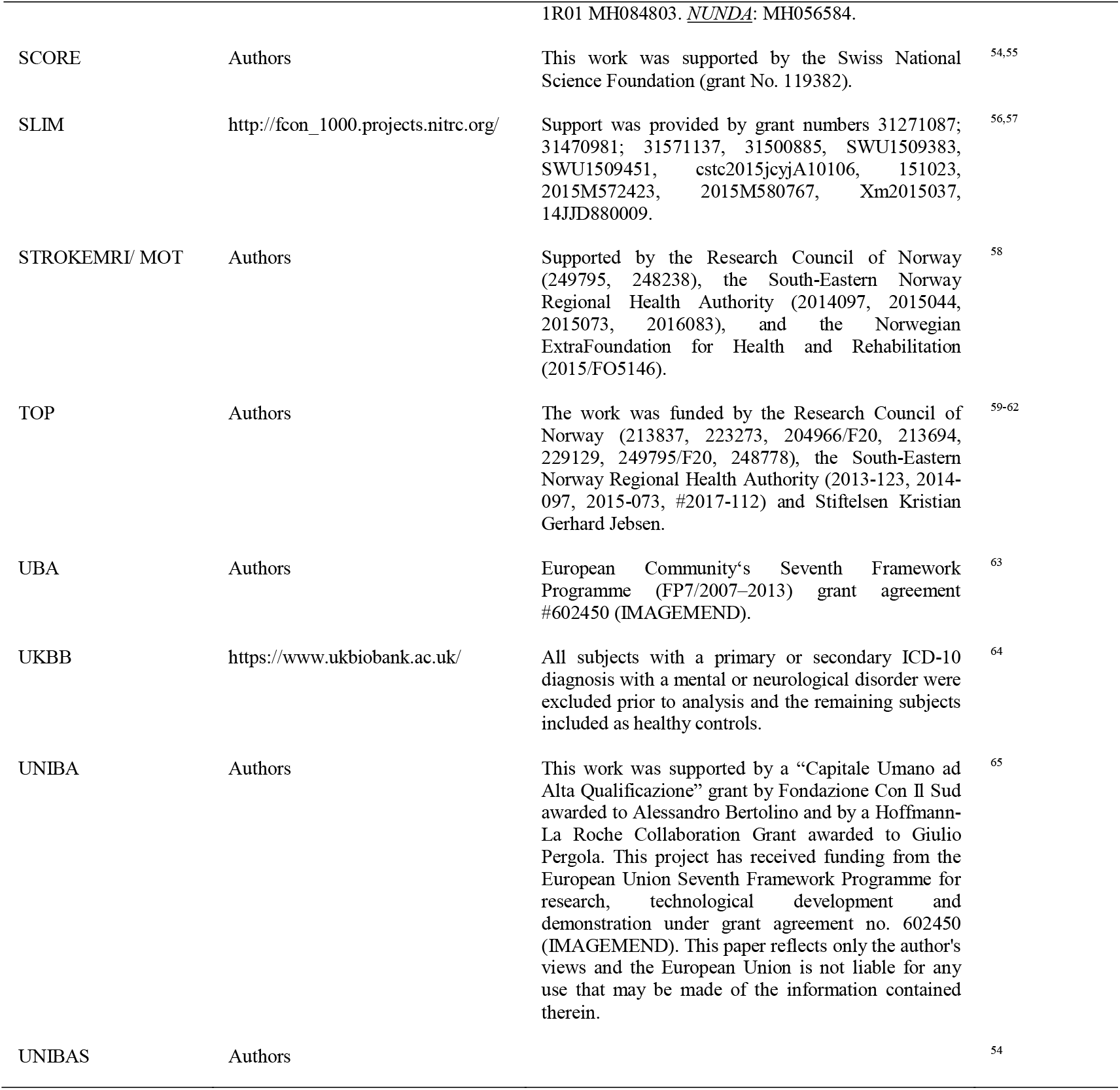
Summary of included samples.

**Supplementary Table 2:**
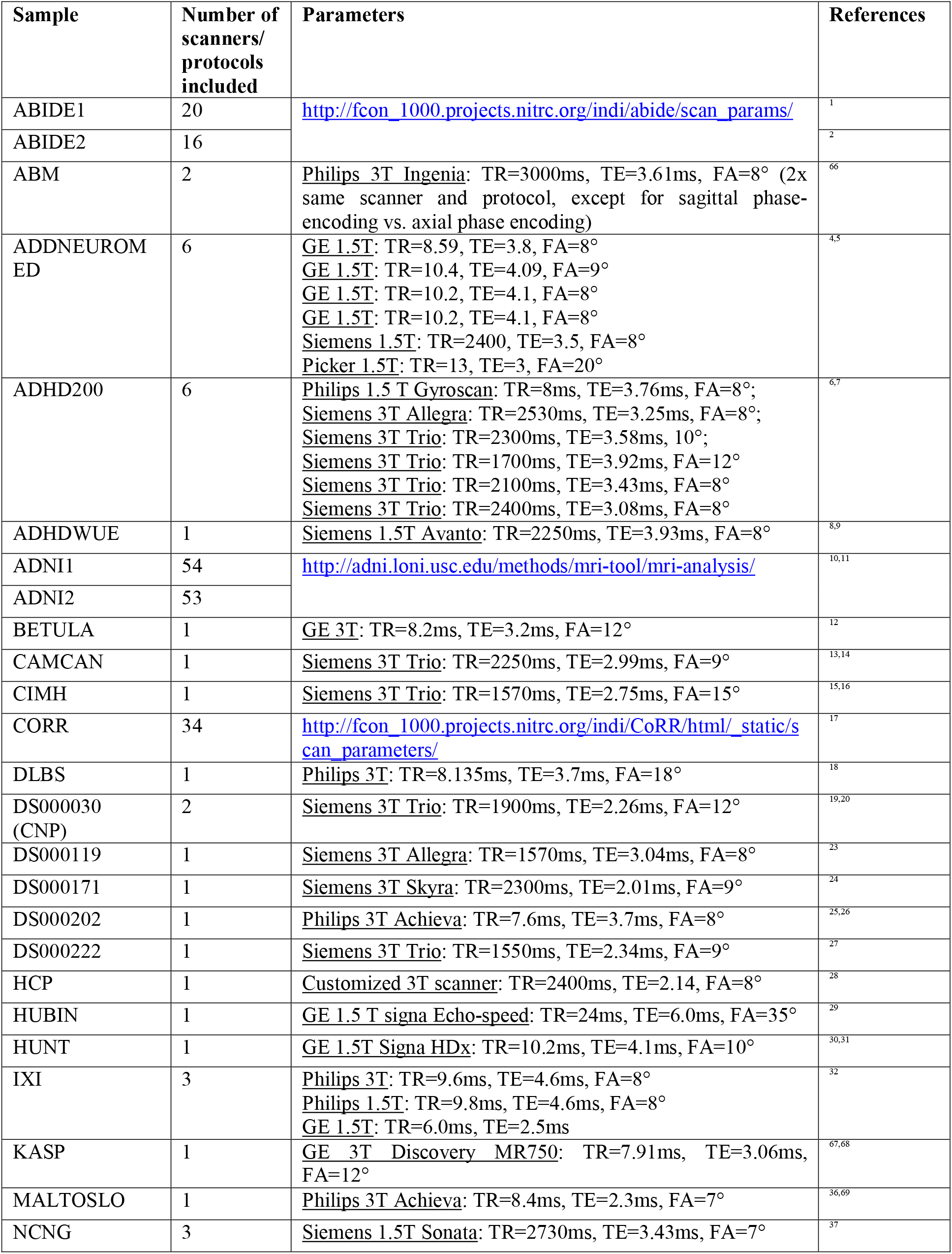
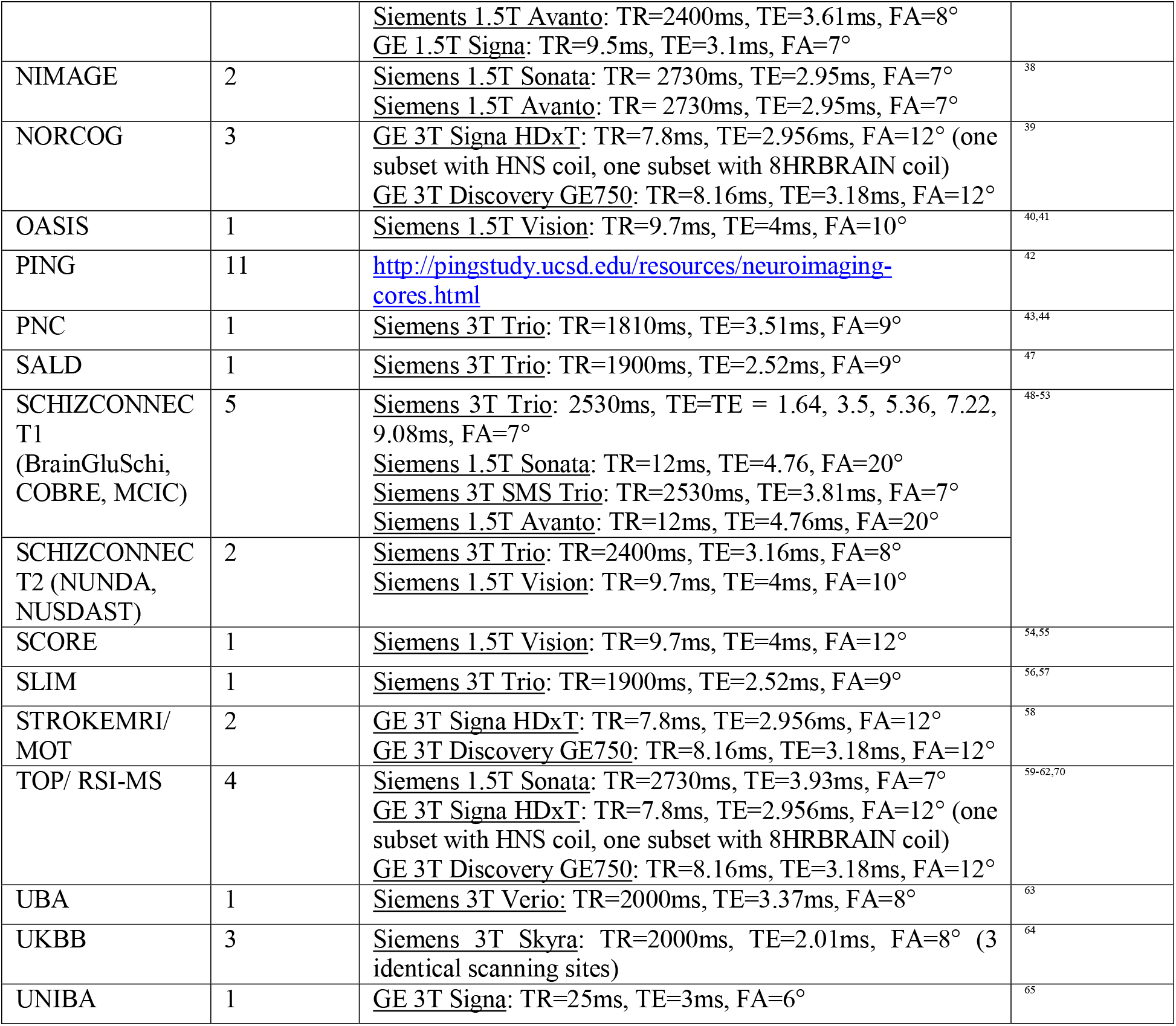
Summary of magnetic resonance imaging characteristics of included samples.

**Supplementary Table 3:** Size and demographic information of final study samples after quality control procedures.

| Sample | Number of subjects per group | Age: mean ± sd in years (group) | Sex: f/m |
| --- | --- | --- | --- |
| ADHD | 681 patients; 992 HC | 17.5±9.9 (ADHD); 17.4±9.7 (HC) | 702/971 |
| ASD | 125 patients; 140 HC | 18.7±9.7 (ASD); 17.1±9.8 (HC) | 34/231 |
| BD | 464 patients; 1,531 HC | 33.7±10.7 (BD); 39.1±15.9 (HC) | 1,031/946 |
| MDD | 211 patients; 93 HC | 38.8±13.5 (MDD); 39.5±13.9 (HC) | 201/103 |
| PSYMIX | 308 patients; 1,430 HC | 28.9±9.3 (PSYMIX); 39.3±16.2 (HC) | 852/886 |
| SCZ | 1,044 patients; 2,079 HC | 33.6±11.0 (SCZ); 37.7±15.2 (HC) | 1,323/1,800 |
| SCZRISK | 91 patients; 402 HC | 23.7±5.0 (SCZRISK); 31.2±11.7 (HC) | 223/270 |
| DEM | 756 patients; 1,921 HC | 75.4±7.7 (DEM); 51.4±22.0 (HC) | 1,434/1,243 |
| MCI | 987 patients; 1,655 HC | 72.2±8.4 (MCI); 53.3±21.3 (HC) | 1,264/1,378 |
| MS | 257 patients; 1,053 HC | 40.9±10.0 (MS); 41.4±17.6 (HC) | 745/565 |
| PD | 138 patients; 67 HC | 61.2±9.1 (PD); 60.2±11.3 (HC) | 73/132 |
| UK Biobank sample | 27,034 HC | 55.6±7.4 (HC) | 13,931/13,103 |
| All HC | 38,299 HC | 50.0±15.7 (HC) | 19,963/18,336 |
| All participants | 43,353 |  |  |

ADHD; attention-deficit/hyperactivity disorder. ASD; autism spectrum disorders. BD; bipolar disorder. HC; healthy controls. MDD; major depression. PSYMIX; non-SCZ psychosis spectrum diagnoses. SCZ; schizophrenia.
SCZRISK; prodromal SCZ or at risk mental state. DEM; dementia. MCI; mild cognitive impairment. MS; multiple sclerosis. PD; Parkinson's disease.

## References

1. Guyenet, P. G. The sympathetic control of blood pressure. Nature reviews Neuroscience 7, 335–346 (2006).

2. Del Negro, C. A., Funk, G. D. Breathing matters. Nature reviews Neuroscience 19, 351–367 (2018).

3. Damasio, A., Carvalho, G. B. The nature of feelings: evolutionary and neurobiological origins. Nature reviews Neuroscience 14, 143–152 (2013).

4. Fisman, M. The brain stem in psychosis. The British journal of psychiatry: the journal of mental science 126, 414–422 (1975).

5. Williams, D. R., Lees, A. J. Progressive supranuclear palsy: clinicopathological concepts and diagnostic challenges. The Lancet Neurology 8, 270–279 (2009).

6. Lang, A. E., Lozano, A. M. Parkinson’s disease. First of two parts. The New England journal of medicine 339, 1044–1053 (1998).

7. Krishnan, V., Nestler, E. J. The molecular neurobiology of depression. Nature 455, 894–902 (2008).

8. Sara, S. J. The locus coeruleus and noradrenergic modulation of cognition. Nature reviews Neuroscience 10, 211–223 (2009).

9. Nakamura, K. et al. Brain serotonin and dopamine transporter bindings in adults with high-functioning autism. Archives of general psychiatry 67, 59–68 (2010).

10. Grace, A. A. Dysregulation of the dopamine system in the pathophysiology of schizophrenia and depression. Nature reviews Neuroscience 17, 524–532 (2016).

11. Przedborski, S. The two-century journey of Parkinson disease research. Nature reviews Neuroscience 18, 251–259 (2017).

12. van Erp, T. G. et al. Subcortical brain volume abnormalities in 2028 individuals with schizophrenia and 2540 healthy controls via the ENIGMA consortium. Molecular psychiatry 21, 547–553 (2016).

13. Hibar, D. P. et al. Cortical abnormalities in bipolar disorder: an MRI analysis of 6503 individuals from the ENIGMA Bipolar Disorder Working Group. Molecular psychiatry 23, 932–942 (2018).

14. Whelan, C. D. et al. Structural brain abnormalities in the common epilepsies assessed in a worldwide ENIGMA study. Brain: a journal of neurology 141, 391–408 (2018).

15. Schmaal, L. et al. Cortical abnormalities in adults and adolescents with major depression based on brain scans from 20 cohorts worldwide in the ENIGMA Major Depressive Disorder Working Group. Molecular psychiatry 6, 900–909 (2016).

16. Hibar, D. P. et al. Novel genetic loci associated with hippocampal volume. Nature communications 8, 13624 (2017).

17. Franke, B. et al. Genetic influences on schizophrenia and subcortical brain volumes: large-scale proof of concept. Nature neuroscience 19, 420–431 (2016).

18. Hibar, D. P. et al. Common genetic variants influence human subcortical brain structures. Nature 520, 224–229 (2015).

19. Thinking big in mental health. Nature medicine 24, 1 (2018).

20. Iglesias, J. E. et al. Bayesian segmentation of brainstem structures in MRI. NeuroImage 113, 184–195 (2015).

21. Fischl, B. et al. Whole brain segmentation: automated labeling of neuroanatomical structures in the human brain. Neuron 33, 341–355 (2002).

22. Sudlow, C. et al. UK biobank: an open access resource for identifying the causes of a wide range of complex diseases of middle and old age. PLoS Medicine 12, e1001779 (2015).

23. Purcell, S. et al. PLINK: a tool set for whole-genome association and population-based linkage analyses. American journal of human genetics 81, 559–575 (2007).

24. van der Meer, D. et al. Brain scans from 21,297 individuals reveal the genetic architecture of hippocampal subfield volumes. Molecular psychiatry 10.1038/s41380-41018-40262-41387 (2018).

25. Bulik-Sullivan, B. et al. An atlas of genetic correlations across human diseases and traits. Nature genetics 47, 1236–1241 (2015).

26. Watanabe, K., Taskesen, E., van Bochoven, A., Posthuma, D. Functional mapping and annotation of genetic associations with FUMA. Nature communications 8, 1826 (2017).

27. Ernst, J., Kellis, M. ChromHMM: automating chromatin-state discovery and characterization. Nature methods 9, 215–216 (2012).

28. Kircher, M. et al. A general framework for estimating the relative pathogenicity of human genetic variants. Nature genetics 46, 310–315 (2014).

29. Pickrell, J. K. et al. Detection and interpretation of shared genetic influences on 42 human traits. Nature genetics 48, 709–717 (2016).

30. de Leeuw, C. A., Mooij, J. M., Heskes, T., Posthuma, D. MAGMA: generalized gene-set analysis of GWAS data. PLoS computational biology 11, e1004219 (2015).

31. Garcia-Fernandez, J. The genesis and evolution of homeobox gene clusters. Nature reviews Genetics 6, 881–892 (2005).

32. Trainor, P. A., Krumlauf, R. Patterning the cranial neural crest: hindbrain segmentation and Hox gene plasticity. Nature reviews Neuroscience 1, 116–124 (2000).

33. Kamburov, A., Stelzl, U., Lehrach, H., Herwig, R. The ConsensusPathDB interaction database: 2013 update. Nucleic acids research 41, D793–800 (2013).

34. Andreassen, O. A. et al. Improved detection of common variants associated with schizophrenia by leveraging pleiotropy with cardiovascular-disease risk factors. Am J Hum Genet 92, 197–209 (2013).

35. Liu, J. Z. et al. Dense genotyping of immune-related disease regions identifies nine new risk loci for primary sclerosing cholangitis. Nature genetics 45, 670–675 (2013).

36. Schork, A. J. et al. New statistical approaches exploit the polygenic architecture of schizophrenia--implications for the underlying neurobiology. Current opinion in neurobiology 36, 89–98 (2016).

37. Smeland, O. B. et al. Genome-wide analysis reveals extensive genetic overlap between schizophrenia, bipolar disorder, and intelligence. Molecular psychiatry 10.1038/s41380-41018-40332-x (2019).

38. Smeland, O. B. et al. Identification of Genetic Loci Jointly Influencing Schizophrenia Risk and the Cognitive Traits of Verbal-Numerical Reasoning, Reaction Time, and General Cognitive Function. JAMA psychiatry 74, 1065–1075 (2017).

39. R Core Team. R: A language and environment for statistical computing. R Foundation for Statistical Computing, Vienna, Austria (2013).

40. Folstein, M. F., Folstein, S. E., McHugh, P. R. “Mini-mental state”. A practical method for grading the cognitive state of patients for the clinician. Journal of psychiatric research 12, 189–198 (1975).

41. Kurtzke, J. F. Rating neurologic impairment in multiple sclerosis: an expanded disability status scale (EDSS). Neurology 33, 1444–1452 (1983).

42. Pedersen, G., Karterud, S. The symptom and function dimensions of the Global Assessment of Functioning (GAF) scale. Comprehensive psychiatry 53, 292–298 (2012).

43. Kay, S. R., Fiszbein, A., Opler, L. A. The positive and negative syndrome scale (PANSS) for schizophrenia. Schizophrenia bulletin 13, 261–276 (1987).

44. Elliott, L. T. et al. Genome-wide association studies of brain imaging phenotypes in UK Biobank. Nature 562, 210–216 (2018).

45. Wang, Y., Mandelkow, E. Tau in physiology and pathology. Nature Reviews Neuroscience 17, 22 (2015).

46. Chang, D. et al. A meta-analysis of genome-wide association studies identifies 17 new Parkinson’s disease risk loci. Nature genetics 49, 1511–1516 (2017).

47. Hoglinger, G. U. et al. Identification of common variants influencing risk of the tauopathy progressive supranuclear palsy. Nature genetics 43, 699–705 (2011).

48. Lyons, D. A., Naylor, S. G., Scholze, A., Talbot, W. S. Kif1b is essential for mRNA localization in oligodendrocytes and development of myelinated axons. Nature genetics 41, 854–858 (2009).

49. Zhao, C. et al. Charcot-Marie-Tooth disease type 2A caused by mutation in a microtubule motor KIF1Bbeta. Cell 105, 587–597 (2001).

50. Ahmad, F. J., Yu, W., McNally, F. J., Baas, P. W. An essential role for katanin in severing microtubules in the neuron. The Journal of cell biology 145, 305–315 (1999).

51. Toyo-Oka, K. et al. Recruitment of katanin p60 by phosphorylated NDEL1, an LIS1 interacting protein, is essential for mitotic cell division and neuronal migration. Human molecular genetics 14, 3113–3128 (2005).

52. Checler, F., Vincent, J. P., Kitabgi, P. Purification and characterization of a novel neurotensin-degrading peptidase from rat brain synaptic membranes. The Journal of biological chemistry 261, 11274–11281 (1986).

53. Boules, M. et al. Diverse roles of neurotensin agonists in the central nervous system. Frontiers in endocrinology 4, 36 (2013).

54. Sharma, R. P., Janicak, P. G., Bissette, G., Nemeroff, C. B. CSF neurotensin concentrations and antipsychotic treatment in schizophrenia and schizoaffective disorder. The American journal of psychiatry 154, 1019–1021 (1997).

55. Vuong, T. A. et al. SGTb regulates a surface localization of a guidance receptor BOC to promote neurite outgrowth. Cellular signalling 55, 100–108 (2019).

56. Webb, B. D. et al. HOXB1 founder mutation in humans recapitulates the phenotype of Hoxb1−/− mice. American journal of human genetics 91, 171–179 (2012).

57. Kohler, M. et al. Small-conductance, calcium-activated potassium channels from mammalian brain. Science (New York, NY) 273, 1709–1714 (1996).

58. Zollino, M. et al. Mutations in KANSL1 cause the 17q21.31 microdeletion syndrome phenotype. Nature genetics 44, 636–638 (2012).

59. Frei, O. et al. Bivariate causal mixture model quantifies polygenic overlap between complex traits beyond genetic correlation. Nature communications 10, 2417 (2019).

60. Mirsky, A. F., Duncan, C. C. Pathophysiology of mental illness: a view from the fourth ventricle. International journal of psychophysiology: official journal of the International Organization of Psychophysiology 58, 162–178 (2005).

61. Schildkraut, J. J. The catecholamine hypothesis of affective disorders: a review of supporting evidence. The American journal of psychiatry 122, 509–522 (1965).

62. Ashok, A. H. et al. The dopamine hypothesis of bipolar affective disorder: the state of the art and implications for treatment. Molecular psychiatry 22, 666–679 (2017).

63. Murray, G. K. et al. Substantia nigra/ventral tegmental reward prediction error disruption in psychosis. Molecular psychiatry 13, 239, 267–276 (2008).

64. Rimol, L. M. et al. Cortical thickness and subcortical volumes in schizophrenia and bipolar disorder. Biological psychiatry 68, 41–50 (2010).

65. Alnaes, D. et al. Brain Heterogeneity in Schizophrenia and Its Association With Polygenic Risk. JAMA psychiatry 76, 739–748 (2019).

66. Ledig, C. et al. Structural brain imaging in Alzheimer’s disease and mild cognitive impairment: biomarker analysis and shared morphometry database. Scientific reports 8, 11258 (2018).

67. Lee, J. H. et al. Brainstem morphological changes in Alzheimer’s disease. Neuroreport 26, 411–415 (2015).

68. Nigro, S. et al. Fully automated segmentation of the pons and midbrain using human T1 MR brain images. PloS one 9, e85618 (2014).

69. Daams, M. et al. Unraveling the neuroimaging predictors for motor dysfunction in long-standing multiple sclerosis. Neurology 85, 248–255 (2015).

70. Lee, C. Y. et al. Differential brainstem atrophy patterns in multiple sclerosis and neuromyelitis optica spectrum disorders. Journal of magnetic resonance imaging: JMRI 47, 1601–1609 (2018).

71. Liu, C. et al. Three dimensional MRI estimates of brain and spinal cord atrophy in multiple sclerosis. Journal of neurology, neurosurgery, and psychiatry 66, 323–330 (1999).

72. Pagonabarraga, J. et al. Neural correlates of minor hallucinations in non-demented patients with Parkinson’s disease. Parkinsonism & related disorders 20, 290–296 (2014).

73. Rektorova, I. et al. Grey matter changes in cognitively impaired Parkinson’s disease patients. PloS one 9, e85595 (2014).

74. Sawczak, C. M., Barnett, A. J., Cohn, M. Increased Cortical Thickness in Attentional Networks in Parkinson’s Disease with Minor Hallucinations. Parkinsons Dis 2019, 5351749 (2019).

75. Goetz, C. G. et al. Movement Disorder Society-sponsored revision of the Unified Parkinson’s Disease Rating Scale (MDS-UPDRS): scale presentation and clinimetric testing results. Movement disorders: official journal of the Movement Disorder Society 23, 2129–2170 (2008).

76. Hoehn, M. M., Yahr, M. D. Parkinsonism: onset, progression and mortality. Neurology 17, 427–442 (1967).

77. Boyle, A. P. et al. Annotation of functional variation in personal genomes using RegulomeDB. Genome research 22, 1790–1797 (2012).

78. Kundaje, A. et al. Integrative analysis of 111 reference human epigenomes. Nature 518, 317–330 (2015).

79. Zhu, Z. et al. Integration of summary data from GWAS and eQTL studies predicts complex trait gene targets. Nature genetics 48, 481–487 (2016).

80. Subramanian, A. et al. Gene set enrichment analysis: a knowledge-based approach for interpreting genome-wide expression profiles. Proceedings of the National Academy of Sciences of the United States of America 102, 15545–15550 (2005).

81. Demontis, D. et al. Discovery of the first genome-wide significant risk loci for attention deficit/hyperactivity disorder. Nature genetics 51, 63–75 (2019).

82. Ripke, S. et al. Biological insights from 108 schizophrenia-associated genetic loci. Nature 511, 421–427 (2014).

83. Meta-analysis of GWAS of over 16,000 individuals with autism spectrum disorder highlights a novel locus at 10q24.32 and a significant overlap with schizophrenia. Molecular autism 8, 21 (2017).

84. Stahl, E. A. et al. Genome-wide association study identifies 30 loci associated with bipolar disorder. Nature genetics 51, 793–803 (2019).

85. Wray, N. R. et al. Genome-wide association analyses identify 44 risk variants and refine the genetic architecture of major depression. Nature genetics 50, 668–681 (2018).

86. Hyde, C. L. et al. Identification of 15 genetic loci associated with risk of major depression in individuals of European descent. Nature genetics 48, 1031–1036 (2016).

87. Lambert, J. C. et al. Meta-analysis of 74,046 individuals identifies 11 new susceptibility loci for Alzheimer’s disease. Nature genetics 45, 1452–1458 (2013).

88. Sawcer, S. et al. Genetic risk and a primary role for cell-mediated immune mechanisms in multiple sclerosis. Nature 476, 214–219 (2011).

89. Nalls, M. A. et al. Large-scale meta-analysis of genome-wide association data identifies six new risk loci for Parkinson’s disease. Nature genetics 46, 989–993 (2014).

90. Andreassen, O. A. et al. Improved detection of common variants associated with schizophrenia and bipolar disorder using pleiotropy-informed conditional false discovery rate. PLoS genetics 9, e1003455 (2013).

91. Nichols, T. et al. Valid conjunction inference with the minimum statistic. NeuroImage 25, 653–660 (2005).

## Supplementary references

1. Di Martino A, Yan CG, Li Q, Denio E, Castellanos FX, Alaerts K et al. The autism brain imaging data exchange: towards a large-scale evaluation of the intrinsic brain architecture in autism. Molecular psychiatry 2014; 19: 659–667.

2. Di Martino A, O’Connor D, Chen B, Alaerts K, Anderson JS, Assaf M et al. Enhancing studies of the connectome in autism using the autism brain imaging data exchange II. Sci Data 2017; 4: 170010.

3. Maglanoc LA, Landro NI, Jonassen R, Kaufmann T, Cordova-Palomera A, Hilland E et al. Data-Driven Clustering Reveals a Link Between Symptoms and Functional Brain Connectivity in Depression. *Biological psychiatry Cognitive neuroscience and neuroimaging* 2018.

4. Liu Y, Paajanen T, Zhang Y, Westman E, Wahlund LO, Simmons A et al. Combination analysis of neuropsychological tests and structural MRI measures in differentiating AD, MCI and control groups--the AddNeuroMed study. Neurobiol Aging 2011; 32: 1198–1206.

5. Lovestone S, Francis P, Strandgaard K. Biomarkers for disease modification trials--the innovative medicines initiative and AddNeuroMed. J Nutr Health Aging 2007; 11: 359–361.

6. Brown MR, Sidhu GS, Greiner R, Asgarian N, Bastani M, Silverstone PH et al. ADHD-200 Global Competition: diagnosing ADHD using personal characteristic data can outperform resting state fMRI measurements. Front Syst Neurosci 2012; 6: 69.

7. Consortium HD. The ADHD-200 Consortium: A Model to Advance the Translational Potential of Neuroimaging in Clinical Neuroscience. Front Syst Neurosci 2012; 6: 62.

8. Guadalupe T, Mathias SR, vanErp TGM, Whelan CD, Zwiers MP, Abe Y et al. Human subcortical brain asymmetries in 15,847 people worldwide reveal effects of age and sex. Brain Imaging Behav 2017; 11: 1497–1514.

9. Conzelmann A, Mucha RF, Jacob CP, Weyers P, Romanos J, Gerdes AB et al. Abnormal affective responsiveness in attention-deficit/hyperactivity disorder: subtype differences. Biological psychiatry 2009; 65: 578–585.

10. Weiner MW, Aisen PS, Jack CR, Jr., Jagust WJ, Trojanowski JQ, Shaw L et al. The Alzheimer’s disease neuroimaging initiative: progress report and future plans. Alzheimers Dement 2010; 6: 202–211 e207.

11. Wyman BT, Harvey DJ, Crawford K, Bernstein MA, Carmichael O, Cole PE et al. Standardization of analysis sets for reporting results from ADNI MRI data. Alzheimers Dement 2013; 9: 332–337.

12. Nilsson L-G, Adolfsson R, Bäckman L, de Frias CM, Molander B, Nyberg L. Betula: A Prospective Cohort Study on Memory, Health and Aging. *Aging*, Neuropsychology, and Cognition 2004; 11: 134–148.

13. Taylor JR, Williams N, Cusack R, Auer T, Shafto MA, Dixon M et al. The Cambridge Centre for Ageing and Neuroscience (Cam-CAN) data repository: Structural and functional MRI, MEG, and cognitive data from a cross-sectional adult lifespan sample. Neuroimage 2017; 144: 262–269.

14. Shafto MA, Tyler LK, Dixon M, Taylor JR, Rowe JB, Cusack R et al. The Cambridge Centre for Ageing and Neuroscience (Cam-CAN) study protocol: a cross-sectional, lifespan, multidisciplinary examination of healthy cognitive ageing. BMC Neurol 2014; 14: 204.

15. Eisenacher S, Rausch F, Ainser F, Mier D, Veckenstedt R, Schirmbeck F et al. Investigation of metamemory functioning in the at-risk mental state for psychosis. Psychological medicine 2015; 45: 3329–3340.

16. Rausch F, Mier D, Eifler S, Esslinger C, Schilling C, Schirmbeck F et al. Reduced activation in ventral striatum and ventral tegmental area during probabilistic decision-making in schizophrenia. Schizophrenia research 2014; 156: 143–149.

17. Zuo XN, Anderson JS, Bellec P, Birn RM, Biswal BB, Blautzik J et al. An open science resource for establishing reliability and reproducibility in functional connectomics. Scientific data 2014; 1: 140049.

18. Lu H, Xu F, Rodrigue KM, Kennedy KM, Cheng Y, Flicker B et al. Alterations in cerebral metabolic rate and blood supply across the adult lifespan. Cereb Cortex 2011; 21: 1426–1434.

19. Gorgolewski KJ, Durnez J, Poldrack RA. Preprocessed Consortium for Neuropsychiatric Phenomics dataset. F1000Res 2017; 6: 1262.

20. Poldrack RA, Congdon E, Triplett W, Gorgolewski KJ, Karlsgodt KH, Mumford JA et al. A phenome-wide examination of neural and cognitive function. Sci Data 2016; 3: 160110.

21. Repovs G, Barch DM. Working memory related brain network connectivity in individuals with schizophrenia and their siblings. Frontiers in human neuroscience 2012; 6: 137.

22. Repovs G, Csernansky JG, Barch DM. Brain network connectivity in individuals with schizophrenia and their siblings. Biological psychiatry 2011; 69: 967–973.

23. Velanova K, Wheeler ME, Luna B. Maturational changes in anterior cingulate and frontoparietal recruitment support the development of error processing and inhibitory control. Cereb Cortex 2008; 18: 2505–2522.

24. Lepping RJ, Ruth AA, Cary R. Development of a validated emotionally provocative musical stimulus set for researc. Psychology of music 2016; 44.

25. Van Schuerbeek P, Baeken C, De Mey J. The Heterogeneity in Retrieved Relations between the Personality Trait ‘Harm Avoidance’ and Gray Matter Volumes Due to Variations in the VBM and ROI Labeling Processing Settings. PloS one 2016; 11: e0153865.

26. Van Schuerbeek P, Baeken C, De Raedt R, De Mey J, Luypaert R. Individual differences in local gray and white matter volumes reflect differences in temperament and character: a voxel-based morphometry study in healthy young females. Brain Res 2011; 1371: 32–42.

27. FitzGerald THB, Hammerer D, Friston KJ, Li SC, Dolan RJ. Sequential inference as a mode of cognition and its correlates in fronto-parietal and hippocampal brain regions. PLoS Comput Biol 2017; 13: e1005418.

28. Van Essen DC, Smith SM, Barch DM, Behrens TE, Yacoub E, Ugurbil K et al. The WU-Minn Human Connectome Project: an overview. Neuroimage 2013; 80: 62–79.

29. Haukvik UK, Schaer M, Nesvag R, McNeil T, Hartberg CB, Jonsson EG et al. Cortical folding in Broca’s area relates to obstetric complications in schizophrenia patients and healthy controls. Psychol Med 2012; 42: 1329–1337.

30. Haberg AK, Hammer TA, Kvistad KA, Rydland J, Muller TB, Eikenes L et al. Incidental Intracranial Findings and Their Clinical Impact; The HUNT MRI Study in a General Population of 1006 Participants between 50-66 Years. PloS one 2016; 11: e0151080.

31. Krokstad S, Langhammer A, Hveem K, Holmen TL, Midthjell K, Stene TR et al. Cohort Profile: the HUNT Study, Norway. Int J Epidemiol 2013; 42: 968–977.

32. Liu K, Yao S, Chen K, Zhang J, Yao L, Li K et al. Structural Brain Network Changes across the Adult Lifespan. Front Aging Neurosci 2017; 9: 275.

33. Collste K, Plaven-Sigray P, Fatouros-Bergman H, Victorsson P, Schain M, Forsberg A et al. Lower levels of the glial cell marker TSPO in drug-naive first-episode psychosis patients as measured using PET and [(11)C]PBR28. Molecular psychiatry 2017; 22: 850–856.

34. Orhan F, Fatouros-Bergman H, Goiny M, Malmqvist A, Piehl F, Cervenka S et al. CSF GABA is reduced in first-episode psychosis and associates to symptom severity. Molecular psychiatry 2018; 23: 1244–1250.

35. Elvsashagen T, Westlye LT, Boen E, Hol PK, Andreassen OA, Boye B et al. Bipolar II disorder is associated with thinning of prefrontal and temporal cortices involved in affect regulation. Bipolar Disord, vol. doi: 10.1111/bdi.121172013.

36. Elvsashagen T, Westlye LT, Boen E, Hol PK, Andersson S, Andreassen OA et al. Evidence for reduced dentate gyrus and fimbria volume in bipolar II disorder. Bipolar disorders 2013; 15: 167–176.

37. Espeseth T, Christoforou A, Lundervold AJ, Steen VM, Le Hellard S, Reinvang I. Imaging and cognitive genetics: the Norwegian Cognitive NeuroGenetics sample. Twin Res Hum Genet 2012; 15: 442–452.

38. von Rhein D, Mennes M, van Ewijk H, Groenman AP, Zwiers MP, Oosterlaan J et al. The NeuroIMAGE study: a prospective phenotypic, cognitive, genetic and MRI study in children with attention-deficit/hyperactivity disorder. Design and descriptives. Eur Child Adolesc Psychiatry 2015; 24: 265–281.

39. Doan NT, Engvig A, Zaske K, Persson K, Lund MJ, Kaufmann T et al. Distinguishing early and late brain aging from the Alzheimer’s disease spectrum: consistent morphological patterns across independent samples. Neuroimage 2017; 158: 282–295.

40. Buckner RL, Head D, Parker J, Fotenos AF, Marcus D, Morris JC et al. A unified approach for morphometric and functional data analysis in young, old, and demented adults using automated atlas-based head size normalization: reliability and validation against manual measurement of total intracranial volume. Neuroimage 2004; 23: 724–738.

41. Fotenos AF, Snyder AZ, Girton LE, Morris JC, Buckner RL. Normative estimates of cross-sectional and longitudinal brain volume decline in aging and AD. Neurology 2005; 64: 1032–1039.

42. Jernigan TL, Brown TT, Hagler DJ, Jr., Akshoomoff N, Bartsch H, Newman E et al. The Pediatric Imaging, Neurocognition, and Genetics (PING) Data Repository. Neuroimage 2016; 124: 1149–1154.

43. Satterthwaite TD, Connolly JJ, Ruparel K, Calkins ME, Jackson C, Elliott MA et al. The Philadelphia Neurodevelopmental Cohort: A publicly available resource for the study of normal and abnormal brain development in youth. Neuroimage 2016; 124: 1115–1119.

44. Satterthwaite TD, Elliott MA, Ruparel K, Loughead J, Prabhakaran K, Calkins ME et al. Neuroimaging of the Philadelphia neurodevelopmental cohort. Neuroimage 2014; 86: 544–553.

45. The Parkinson Progression Marker Initiative (PPMI). Progress in neurobiology 2011; 95: 629–635.

46. Sowa P, Harbo HF, White NS, Celius EG, Bartsch H, Berg-Hansen P, et al. Restriction spectrum imaging of white matter and its relation to neurological disability in multiple sclerosis. 2018; 1352458518765671.

47. Wei D, Zhuang K, Chen Q, Yang W, Luiu W, Wang K et al. Structural and functional MRI from a cross-sectional Southwest University Adult lifespan Dataset (SALD). bioRxiv 2018.

48. Bustillo JR, Jones T, Chen H, Lemke N, Abbott C, Qualls C et al. Glutamatergic and Neuronal Dysfunction in Gray and White Matter: A Spectroscopic Imaging Study in a Large Schizophrenia Sample. Schizophr Bull 2017; 43: 611–619.

49. Cetin MS, Christensen F, Abbott CC, Stephen JM, Mayer AR, Canive JM et al. Thalamus and posterior temporal lobe show greater inter-network connectivity at rest and across sensory paradigms in schizophrenia. Neuroimage 2014; 97: 117–126.

50. Gollub RL, Shoemaker JM, King MD, White T, Ehrlich S, Sponheim SR et al. The MCIC collection: a shared repository of multi-modal, multi-site brain image data from a clinical investigation of schizophrenia. Neuroinformatics 2013; 11: 367–388.

51. Kogan A, Alpert K, Ambite JL, Marcus DS, Wang L. Northwestern University schizophrenia data sharing for SchizConnect: A longitudinal dataset for large-scale integration. Neuroimage 2016; 124: 1196–1201.

52. Wang L, Alpert KI, Calhoun VD, Cobia DJ, Keator DB, King MD et al. SchizConnect: Mediating neuroimaging databases on schizophrenia and related disorders for large-scale integration. Neuroimage 2016; 124: 1155–1167.

53. Ambite JL, Tallis M, Alpert K, Keator DB, King M, Landis D et al. SchizConnect: Virtual Data Integration in Neuroimaging. Data Integr Life Sci 2015; 9162: 37–51.

54. Borgwardt S, Koutsouleris N, Aston J, Studerus E, Smieskova R, Riecher-Rossler A et al. Distinguishing prodromal from first-episode psychosis using neuroanatomical single-subject pattern recognition. Schizophr Bull 2013; 39: 1105–1114.

55. Dukart J, Smieskova R, Harrisberger F, Lenz C, Schmidt A, Walter A et al. Age-related brain structural alterations as an intermediate phenotype of psychosis. Journal of psychiatry & neuroscience: JPN 2017; 42: 307–319.

56. Wang Y, Wei D, Li W, Qiu J. Individual differences in brain structure and resting-state functional connectivity associated with type A behavior pattern. Neuroscience 2014; 272: 217–228.

57. Zhu W, Chen Q, Tang C, Cao G, Hou Y, Qiu J. Brain structure links everyday creativity to creative achievement. Brain Cogn 2016; 103: 70–76.

58. Dorum ES, Alnaes D, Kaufmann T, Richard G, Lund MJ, Tonnesen S et al. Age-related differences in brain network activation and co-activation during multiple object tracking. Brain Behav 2016; 6: e00533.

59. Kaufmann T, Alnaes D, Brandt CL, Doan NT, Kauppi K, Bettella F et al. Task modulations and clinical manifestations in the brain functional connectome in 1615 fMRI datasets. Neuroimage 2016; 147: 243–252.

60. Kaufmann T, Skatun KC, Alnaes D, Doan NT, Duff EP, Tonnesen S et al. Disintegration of Sensorimotor Brain Networks in Schizophrenia. Schizophr Bull 2015.

61. Skåtun KC, Kaufmann T, Tonnesen S, Biele G, Melle I, Agartz I et al. Global brain connectivity alterations in patients with schizophrenia and bipolar spectrum disorders. Journal of psychiatry & neuroscience: JPN 2016; 41: 150159.

62. Brandt CL, Kaufmann T, Agartz I, Hugdahl K, Jensen J, Ueland T et al. Cognitive Effort and Schizophrenia Modulate Large-Scale Functional Brain Connectivity. Schizophr Bull 2015.

63. Heck A, Fastenrath M, Ackermann S, Auschra B, Bickel H, Coynel D et al. Converging genetic and functional brain imaging evidence links neuronal excitability to working memory, psychiatric disease, and brain activity. Neuron 2014; 81: 1203–1213.

64. Alfaro-Almagro F, Jenkinson M, Bangerter NK, Andersson JLR, Griffanti L, Douaud G et al. Image processing and Quality Control for the first 10,000 brain imaging datasets from UK Biobank. Neuroimage 2018; 166: 400–424.

65. Pergola G, Trizio S, Di Carlo P, Taurisano P, Mancini M, Amoroso N et al. Grey matter volume patterns in thalamic nuclei are associated with familial risk for schizophrenia. Schizophrenia research 2017; 180: 13–20.

66. Maglanoc LA, Landro NI, Jonassen R, Kaufmann T, Cordova-Palomera A, Hilland E et al. Data-Driven Clustering Reveals a Link Between Symptoms and Functional Brain Connectivity in Depression. Biol Psychiatry Cogn Neurosci Neuroimaging 2019; 4: 16–26.

67. Collste K, Plaven-Sigray P, Fatouros-Bergman H, Victorsson P, Schain M, Forsberg A et al. Lower levels of the glial cell marker TSPO in drug-naive first-episode psychosis patients as measured using PET and [(11)C]PBR28. Molecular psychiatry 2017; 22: 850–856.

68. Orhan F, Fatouros-Bergman H, Goiny M, Malmqvist A, Piehl F, Karolinska Schizophrenia Project C et al. CSF GABA is reduced in first-episode psychosis and associates to symptom severity. Molecular psychiatry 2017.

69. Elvsashagen T, Westlye LT, Boen E, Hol PK, Andreassen OA, Boye B et al. Bipolar II disorder is associated with thinning of prefrontal and temporal cortices involved in affect regulation. Bipolar disorders 2013; 15: 855–864.

70. Sowa P, Harbo HF, White NS, Celius EG, Bartsch H, Berg-Hansen P et al. Restriction spectrum imaging of white matter and its relation to neurological disability in multiple sclerosis. Mult Scler 2018; 1352458518765671.

